# Tracing the evolutionary histories of ultra-rare variants using variational dating of large ancestral recombination graphs

**DOI:** 10.64898/2026.01.07.698223

**Authors:** Nathaniel S. Pope, Sam Tallman, Ben Jeffery, Duncan Robertson, Yan Wong, Savita Karthikeyan, Peter L. Ralph, Jerome Kelleher

**Author notes:** Joint first author (Nathaniel S. Pope, Sam Tallman, Ben Jeffery). Joint senior author (Peter L. Ralph, Jerome Kelleher).

## Abstract

Ultra-rare variants dominate whole-genome sequencing datasets, yet their interpretation is limited by allele frequency, which provides little information at very low counts and is highly sensitive to uneven ancestry representation. Allele age offers an ancestry-agnostic alternative but existing methods do not scale to biobank-sized cohorts. Here we present a scalable variational algorithm for dating Ancestral Recombination Graphs (ARGs), implemented in tsdate, together with new distributed methods enabling practical biobank-scale ARG inference using tsinfer. Applied to 47,535 genomes from the Genomics England 100,000 Genomes Project, we infer contiguous ARGs spanning 206 Mb and estimate ages for 23.2 million variants, including 11.8 million singletons. ARG-based allele ages remain accurate under extreme sampling imbalance and, in real data, reveal signatures of purifying selection and clinically relevant heterogeneity among variants with identical observed frequencies. Estimates for recent mutations are precise only at large sample sizes, highlighting the information accessible in the haplotype structure of large datasets. Biobank-scale ARGs therefore enable robust, ancestry-agnostic age estimation for ultra-rare variation with broad utility for statistical and clinical genomics.

## Introduction

Allele frequency is one of the most widely used descriptors of genetic variation, informing applications that range from clinical variant interpretation^1^ and association testing^2^ to evolutionary inference and models of natural selection.^3–5^ However, the interpretation of allele frequency is fundamentally constrained by how datasets are assembled. In contemporary human datasets, frequency depends not only on evolutionary processes but also on sample size and the geographic, cultural, or clinical contexts in which individuals are recruited, and there are fundamental issues with stratification by ancestry grouping.^6^ Classical population-genetic guidance typically assumes discrete, well-defined populations, an assumption increasingly inconsistent with the continuum of human genetic diversity.^7–9^ Many individuals cannot be unambiguously assigned to a single ancestry group^10^, and large cohorts are usually sampled in highly unbalanced ways.^11^ As a result, allele frequency remains informative for some questions, but for the vast majority of variants (which are ultra rare) its dependence on dataset composition and arbitrary or unstable group definitions makes interpretation difficult and often misleading.^12^ Furthermore, tools designed for discrete population groups often exclude individuals whose ancestry falls into a mixture of standard population labels.

Allele age offers a complementary and, in key respects, more direct summary of evolutionary history. Defined through genealogical relationships rather than population labels, the age of a mutation does not depend on which individuals happen to be included in a particular dataset, and is in principle comparable across cohorts with very different sampling schemes. Classical theory links age, frequency and selection, showing that strongly deleterious alleles are much less likely to be old than neutral alleles.^13, 14^ Early empirical work estimated ages for individual disease-causing mutations using locus-specific data.^15^ More recent approaches have produced tens of millions of genome-wide age estimates,^16–18^ but cannot scale to the billions of overwhelmingly ultra-rare variants now present in modern whole-genome sequencing (WGS) datasets.^19–21^ To obtain informative age estimates across the full allele frequency spectrum—especially for the ultra-rare variants that dominate WGS and are enriched for functional and clinical effects—we need methods that aggregate information along the genome, explicitly model local genealogical relationships, and remain computationally feasible at biobank scale.

Ancestral recombination graphs (ARGs) provide a principled framework for estimating allele ages because they place each mutation in its full genealogical context. For a given set of sampled DNA sequences, an ARG describes the haplotypes and inheritance relationships of all genetic ancestors of the sample, and consequently defines the genealogical trees for those samples at each position along the genome.^22, 23^ Mutation ages correspond to node times within these trees, so ARGs link neighbouring sites through shared ancestry and integrate information over long genomic regions rather than treating variants independently. This property is especially important for rare variation, where haplotype sharing carries most of the available signal. Recent methodological advances have made genome-wide ARG inference increasingly practical, fueling a rapid expansion of applications in population and statistical genetics.^24–28^ Broadly, inference methods fall into two classes. Model-based approaches^29–31^ sample from an explicit population-genetic model and can be highly accurate on simulation benchmarks,^32–35^ but are limited to at most a few hundred genomes and typically require specification of demographic parameters.^36^ Heuristic approaches^37–40^ scale to tens or hundreds of thousands of samples without strong parametric assumptions and separate ARG topology inference from dating. A key feature of these large-scale heuristic methods is that they are ancestry-agnostic: latent population structure is expressed in the inferred ARG without requiring predefined population labels or assumptions about group boundaries.

However, dating ultra-rare alleles requires ultra-large, highly contiguous ARGs, at a scale that has not yet been produced. To obtain accurate ages for very recent mutations, ARG inference must simultaneously encompass large sample sizes and long, continuous genomic regions. Close relatives share extended haplotypes that are identical by descent, manifesting as recent ancestors in the ARG, and artificially truncating these haplotypes by inferring in short chunks (e.g., to distribute computation across a cluster) removes much of the information that constrains their ages and biases date estimates upwards. Similarly, existing general-purpose dating methods,^41, 42^ cannot scale to ultra-large ARGs with tens of millions of nodes and mutations. As a result, there is currently no practical way to obtain accurate, uncertainty-aware ages for the billions of mostly ultra-rare variants in contemporary WGS cohorts.

Here we address this gap by developing a scalable variational ARG-dating algorithm, implemented in tsdate, together with new distributed methods in tsinfer that enable practical biobank-scale ARG inference. The variational algorithm uses expectation propagation to approximate posterior distributions for node times in large ARGs, providing principled, uncertainty-aware age estimates with substantially improved accuracy and efficiency over previous implementations. The distributed tsinfer pipeline infers ARG topologies over long, contiguous genomic segments for tens of thousands of high-coverage genomes. Applied to 47,535 genomes from the Genomics England 100,000 Genomes Project,^19^ we use these methods to infer ARGs spanning 206 Mb and to estimate ages for 23.2 million variants, including 11.8 million singletons. We show that these allele ages remain accurate under extreme sampling imbalance, reveal genealogical structure that is invisible to allele frequency alone, capture signatures of purifying selection from individual variants to regional constraint, and stratify clinically classified variants even at identical observed frequencies. Together, these results demonstrate that scalable ARG inference and dating can now deliver robust, ancestry-agnostic ages for ultra-rare human variation at biobank scale, providing a temporal framework that links evolutionary history, functional impact, and clinical interpretation.

## Results

### Variational Gamma ARG dating

We developed an efficient and accurate method to estimate the age of each ancestral node in an inferred ARG based on a Variational Gamma (VG) approximation of their joint posterior distribution, that is implemented in tsdate version 0.2 (Fig. 1). This implementation uses a Bayesian message-passing algorithm called Expectation Propagation^43^ (EP) to rapidly estimate variational posteriors for both nodes and mutations (Methods §1). These approximate posteriors are compact and capture uncertainty in a principled manner. Posteriors are calibrated by rescaling the ARG to match the average mutation count per sample (Methods §2), and we regularise the ages of unconstrained roots via an exponential prior to avoid unrealistically deep node ages (Methods §3). The method also incorporates a principled approach to coestimating the phase and age of singletons (Methods §4), facilitating the analysis of this large and phenotypically important class of variant.^44^

**Figure 1:**
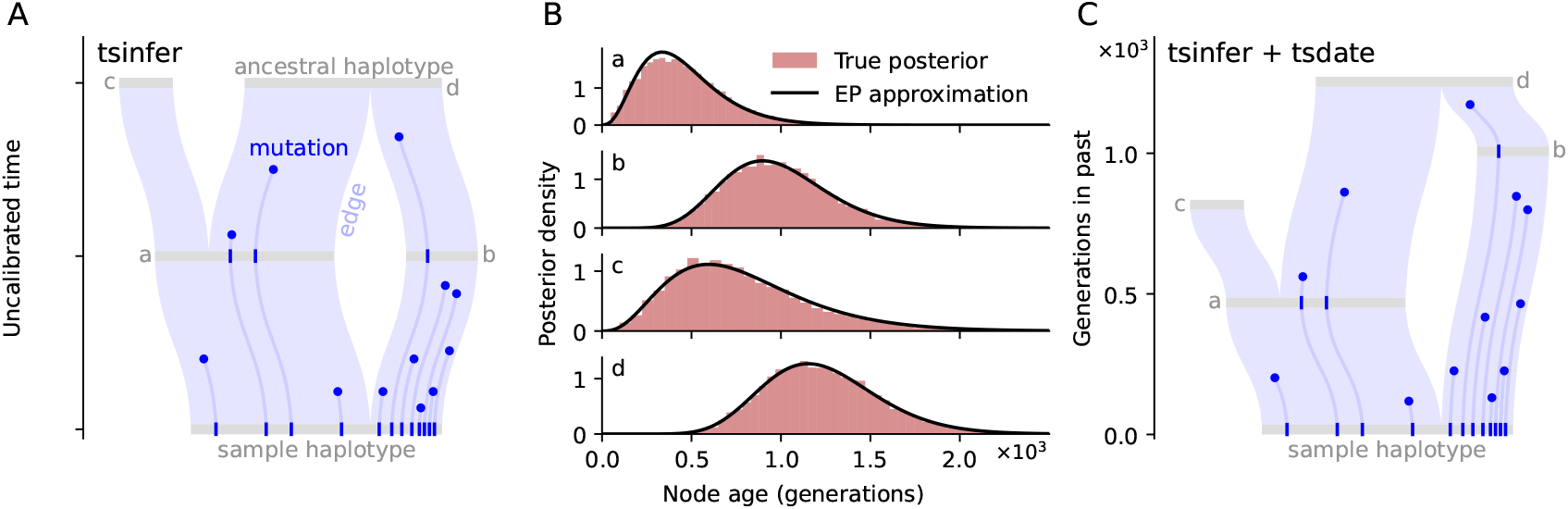
Tsinfer and tsdate use scalable approximations to infer variant ages from biobank-scale sequence data. From left to right: (A) ARGs consists of nodes (ancestral haplotypes) connected by edges (patterns of inheritance) that are localized in time (y-axis) and along the reference sequence (x-axis). Tsinfer infers an ARG topology that is consistent with observed genetic variation in contemporary samples. (B) The density of mutations along edges in the inferred ARG is used by tsdate to estimate (variational) posterior distributions for the ages of nodes. The variational posteriors closely approximate the true posteriors, calculated here by rejection sampling for the ARG shown in (A). (C) The posterior means are used to time-calibrate the ages of nodes in the ARG. Posteriors for mutation ages are derived from those of node ages by assuming that the age of a mutation is equally likely over the length of its edge, and integrating over the ages of the parent and child.

The new VG algorithm is a major improvement over the “inside-outside” (IO) method implemented in tsdate version 0.1^41^ in terms of both accuracy (Fig. S1, Tab. S1, Methods §5) and scalability (Fig. S2). Implemented in Python, leveraging the efficient tskit data model^46, 47^ and Numba acceleration,^48^ the VG algorithm can be applied to the largest available ARGs, requiring only 15 minutes and 7.7GB of RAM to date a simulated ARG of 1 million human chromosomes^45^ on a single thread of an AMD EPYC 7742 CPU. In addition to this scalability, Fig. 2 shows that the accuracy of the VG algorithm improves as sample size grows (in contrast to IO; Fig. S2). Accuracy is comparable to a recently-developed Markov chain Monte Carlo (MCMC) algorithm^42^ that requires orders of magnitude longer runtime for large sample sizes (Fig. S2, Methods §6). Benchmarking indicates that the expectation propagation scheme used by the VG algorithm converges within a small number of iterations (regardless of sample size, Fig. S3) to approximate posteriors that provide well-calibrated uncertainty estimates (Fig. S4).

**Figure 2:**
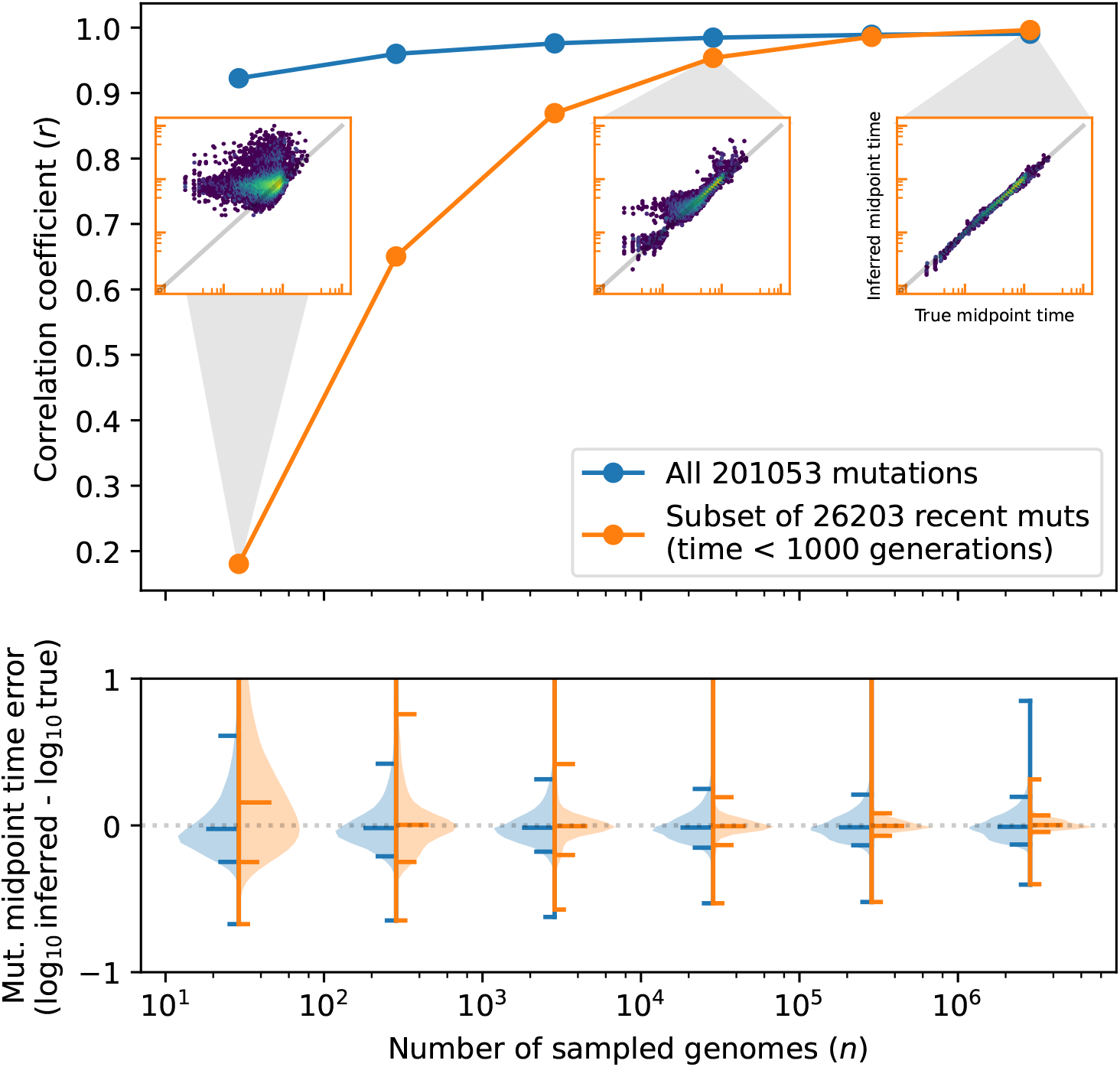
Accuracy of tsdate inference on a pedigree-based simulated tree sequence of 2.85 million samples of human chromosome 17.^45^ Successively smaller subsamples were extracted by simplification, resulting in a smallest tree sequence of 29 samples (0.001% of the original) and 201053 mutations, and tsdate was run on each. In both plots, results are shown for either those 201053 mutations found in all subsamples (blue), or on the subset of these with true time more recent than 1000 generations (orange). Upper: correlation between true and inferred mutation times as a function of sample size (distributions on which *r* is based shown as inset scatterplots); true times are taken as midpoints of node dates above and below the mutation. Lower: error distributions, with bars showing median, 5% and 95% quantiles for errors in log_10_ dates.

To evaluate performance on real data, we ran the combination of tsinfer and the VG algorithm (tsinfer+tsdate, hereafter) on WGS data from the 1000 Genomes Project^49^ (1KGP, Methods §7) and validated against previous results providing estimates of the ages of young, intermediate and ancient variants. We found that inferred ages of > 99% of recent variants were consistent with their first appearance in the ancient DNA record^50^ (Fig. S5, Methods §8); strong agreement with a recent estimate^51^ of the time and magnitude of the Out-of-Africa bottleneck (Fig. S6, Methods §9); and that age estimates of an ancient inversion on chromosome 17q closely correspond to previous work^34^ (Fig. S7, Methods §10). We also evaluated the dependence of mutation age on sample size, showing that mutation ages inferred from subsets of the 1KGP cohort correlate well (Fig. S8). To validate singleton dating, we dated 106,863 singletons in a subset of 1,500 individuals using their estimated phase^49^ and found that the resulting ages correlate more strongly with those from the phase-agnostic method (*r* = 0.807) than with ages obtained under random phasing (*r* = 0.550; Fig. S9). Finally, we compared four published allele age estimates (CoalNN,^18^ GEVA,^17^ Relate,^37^ and SINGER^31^) for 1KGP with tsinfer+tsdate (Fig. S10, Methods §11). Overall, there is considerable variation between the methods, with correlation coefficients ranging from 0.683 (CoalNN/SINGER) to 0.835 (Relate/tsinfer+tsdate), consistent with previous comparisons.^52^

Crucially, tsinfer+tsdate are robust to the biased sampling common in contemporary datasets. To illustrate this we performed a large realistic forwards-time population genetic simulation incorporating negative selection (Methods §12). After running tsinfer and tsdate on the simulated data, the allele ages estimated are accurate and unbiased, even under the highly unbalanced sampling used to mimic the GEL dataset discussed in later sections (Fig. S11). The simulation also illustrates a more fundamental point. It has been shown that allele age offers little information about selection advantage above allele frequency with uniform sampling.^53^ However, the simulation shows that biased sampling induces complex and geographically structured distortions in allele frequency, while allele age remains straightforwardly interpretable. Additionally, in this simulation the combination of allele age and frequency has significantly more power to predict deleteriousness than frequency alone (SI §9).

### Inferring large contiguous ARGs for GEL

Although ARGs have been inferred for hundreds of thousands of samples using genotype array^38, 39^ and imputed^40^ data, there is a considerable gap to the variant densities required for chromosome-scale WGS data. The most recent method, Threads, was applied to approximately 10 million imputed variants genome wide for 487,409 samples, split into 136 chunks,^40^ each of size 30cM (an average of ~ 74,000 variants per chunk). Current biobank-scale WGS datasets, however, consist of around 1 *billion* variants.^21, 54^ In addition to the challenge posed by the number of variants, dating recent, rare alleles requires ARG inference to be performed on long, contiguous regions. For example, close relatives will share long haplotypes that are identical by descent and manifest as recent ancestors (nodes) in the ARG. Artificially truncating the mutational area beneath these recent ancestors will bias their estimated ages upwards (Fig. S12). Thus, to infer accurate dates for recent mutations, we must have both large sample sizes *and* large sections of genome.

To enable such large-scale contiguous ARG inference we added the ability to perform inference distributed over a cluster in tsinfer version 0.4 (Methods §13). The approach distributes work in a deterministic manner, yielding the same results as a single-machine run. We applied this new pipeline to high coverage WGS data for 47,535 individuals in the Genomics England (GEL) 100,000 Genomes Project aggregated (aggV2) dataset^54^ (Methods §14). A central element of tsinfer’s algorithm infers ancestral haplotypes using heuristics. This approach does not perform well in variant-poor regions of the genome, often resulting in ancient haplotypes artefactually spanning such regions. To avoid this, we split chromosome arms into segments based on a site-density filter, yielding 15 segments, ranging from 1.36 to 33Mb, over chromosomes 16– 22 (Tab. S2). Totalling 206Mb (around 14% of the ungapped genome), these regions have a somewhat higher recombination rate and gene density than the genome-wide average because of the bias towards short chromosomes, but are typical in terms of previous allele age estimates (Tab. S2, Fig. S13, SI §8). We performed ARG inference over a total of 11,445,983 non-singleton sites, which required about 16 CPU years (Tab. S3). We then dated the ARGs and phased and dated an additional 11,778,316 singletons, resulting a total of 16.8 million nodes, 141 million edges, 23 million mutations and 2.5GB file size (Tab. S4). We now focus on analysing the final dataset of 23,224,276 allele ages, and in particular on chromosome 17 for several analyses as the largest and most complete chromosome (Methods §14).

### Allele ages in GEL

The ages of the 23 million alleles studied span from very recent to deeply ancient, with 60.8% (*n* = 14, 043, 512) estimated to have occurred within the past 100 generations and 2.3% (*n* = 540, 318) exceeding 10,000 generations (Fig. S14). The frequency spectrum is dominated by ultra-rare variants (Fig. S15), with 94.8% (*n* = 21, 886, 991) having derived allele frequency (DAF) < 0.1% and 41.4% (*n* = 9, 617, 785) absent from gnomAD v4.1.^55^ Allele age correlates positively with DAF (Spearman’s *ρ* = 0.594, *p* < 10^−300^), increasing from a geometric mean of ~51 generations for singletons to ~20,931 generations for common variants (DAF > 10%) (Figs. S16, S17).

As in previous studies,^17, 56^ we find that allele age also reflects functional consequence (Methods §15; Figs. S18, S19). Missense mutations (*n* = 336,127, geometric mean age 81 generations) are younger than non-coding (*n* = 22,517,141, 100 generations) and synonymous variants (*n* = 180,982, 104 generations), whilst the youngest consequence classes are protein-truncating (*n* = 9,994, 63 generations) and splice-altering mutations (*n* = 5,332, 58 generations), consistent with their typically higher functional impact and stronger selective constraint.

### Allele age and ancestral diversity

The GEL aggV2 dataset is a clinically recruited cohort of rare-disease and cancer patients^57^ derived from the 100,000 Genomes Project.^19^ The dataset broadly reflects the UK patient population, with participants predominantly of European descent.^58, 59^ For downstream descriptive analyses, we grouped participants using a PCA-based classifier trained on gnomAD v4.1 (Methods §16) which indicates a strong imbalance of geographic ancestries: 83.7% of individuals (*n* = 39, 806) were assigned to the “non-Finnish European” (NFE) label with high confidence (classification probability > 0.8; Fig. 3a). These groupings are used only for interpretation, and do not enter the ARG inference or dating procedures. We refer to these as “PCA groups” throughout.

**Figure 3:**
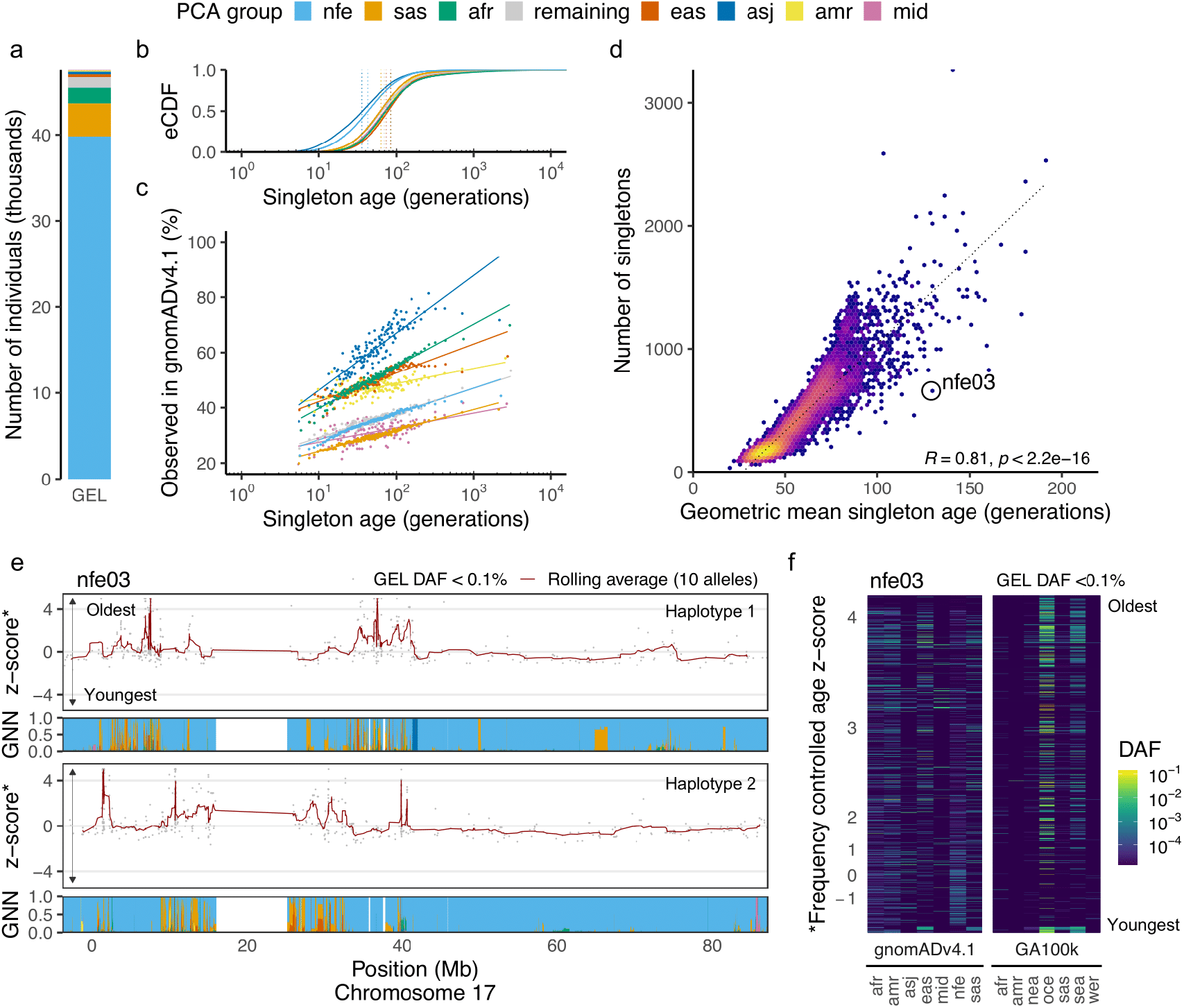
Mutation ages capture ancestral diversity in the 100,000 Genomes Project. (a) Representation of continental “PCA groups” assigned using a classifier trained on gnomADv4.1; see text for abbreviations. (b) Distribution of tsdate inferred ages of singleton alleles (GEL DAC = 1) per PCA group. Dotted lines show the geometric mean of each distribution. (c) Per PCA group, the proportion of singleton mutations at a given mutation age that were observed in gnomADv4.1 (100 quantiles per PCA group). (d) Relationship between number and geometric mean age of singleton variants observed per individual. A participant, nfe03 (ID anonymised for privacy), is highlighted as an example of a member of the NFE PCA group with an excess of older singletons compared to the majority of other members. (e) Frequency-controlled mutation age z-scores (normalised per discrete DAC bin) for ultra-rare (GEL AF < 0.1%) alleles along the two chromosome 17 haplotypes of nfe03. Local GNN proportions colored by PCA group are also shown. We note that the apparent mosaic of divergent lineages inherited from both parents is indicative of switch errors that occurred during phasing. (f) Heatmap depicting external group-specific frequencies of ultra-rare variants observed in nfe03 across gnomADv4.1 (genomes) and GenomeAsia 100k. The y-axis is arranged by age z-score from lowest (bottom) highest (top).

The unequal representation of geographical ancestries in the dataset is reflected in the genealogical relationships in the inferred ARGs and the distribution of mutation ages within discrete derived allele count (DAC) bins. Individuals assigned to the NFE PCA group exhibit a greater density of recent coalescence events (Methods §17, Fig. S20). In contrast, participants assigned to less well-represented PCA groups—including South Asian (SAS, *n* = 3, 837), African (AFR, *n* = 1, 827), and East Asian (EAS, *n* = 335)—show fewer recent coalescence events, consistent with older average times to shared ancestry within the sample. As expected, given their closer genealogical proximity on average to others in the dataset, the NFE PCA group carry singletons (Methods §18) that are on average 28 generations younger than those carried across the remaining individuals (geometric mean age: 43 generations vs. 71 generations for NFE and non-NFE, respectively; Fig.3b). Similar patterns occur in higher DAC bins (Fig. S21).

Inferred mutation age reveals additional biological signal within frequency bins. For example, among singleton variants, we observe a strong positive relationship between the estimated age of a mutation and its probability of being observed in gnomADv4.1 (Fig 3c). This aligns with expectations: older mutations, having had more time to accumulate across more distantly related individuals, are more likely to be shared with external cohorts.

We also analysed how per-individual aggregated singleton ages varied across the cohort. We find that the number of singletons per individual is strongly correlated with the geometric mean age of those singletons (Spearman’s *ρ* = 0.81, *R* = 0.81, *p* < 10^−16^; Fig. 3d), consistent with the expectation that deeper genealogies accumulate more mutations on average. Notably, this relationship is not only apparent across the full dataset but also persists within PCA groups (Fig. S22), highlighting the continuous nature of ancestral relationships apparent within these groups.

We next asked whether these signals could reveal finer-scale ancestry structure across genomic regions within individual genomes. To extend our analysis beyond discrete allele counts and capture variation among all ultra-rare (DAF < 0.1%) mutations, we normalized log10-transformed mutation ages by z-scoring within discrete DAC bins (Methods §19), thereby controlling for the inherent relationship between allele frequency and age. This normalization enabled the identification of genomic regions where mutation ages deviated substantially from expectations conditioned on frequency. We find that these deviations correlate with tracts of differentiated ancestries in an individual, as inferred using the genealogical nearest neighbours (GNN)^38^ proportions. A notable example (Fig. 3e) highlights an individual assigned to the NFE PCA group with local ancestry tracts enriched for mutations with large positive age z-scores. Many of these variants are common polymorphisms (derived allele frequency > 1%) in the Oceanian subset of GenomeAsia 100K^60^ (Fig. 3f). Individuals of Oceanian descent have extremely limited representation in the 100,000 Genomes Project cohort. Consequently, older variants that are common in Oceania are likely to appear superficially rare due to sampling gaps.^61^ Indeed, across the full dataset, the 48,010 DAF < 0.1% variants common in Oceania are 12.9-fold enriched among the most ancient age-frequency outliers (*z* > 4; logistic regression *p* < 10^−300^). This illustrative example shows that genealogically informed metrics such as mutation age can expose informative patterns of rare variant history that may otherwise be obscured when assessing allele frequencies alone. Importantly, however, these signals are dataset-relative: the same genome analysed in a different cohort could exhibit substantially different patterns, reflecting the interaction between allele frequency and genetic ancestries.

### Allele age and patterns of negative selection

Deleterious variants are expected to have younger mutation ages compared to benign variants at the same frequency.^62–64^ To explore whether mutation age could provide information regarding deleterious variant effects, we began by evaluating the 326,121 ultra-rare (DAF < 0.1%) missense variants in our study dataset (Methods §15). Of these, 18.8% (*n* = 61, 402) are classified as likely pathogenic, 71.2% (*n* = 232, 346) as likely benign, and 10% (*n* = 32, 373) as ambiguous using AlphaMissense^65^ with default classification thresholds. In line with expectations of negative selection against functionally disruptive variants, likely pathogenic variants are significantly younger on average (geometric mean: 60 generations) than those classified as likely benign (80 generations; permutation test *p* < 10^−4^). The strongest effect of negative selection is expected to be a deficit of old deleterious alleles. Indeed, the quantiles of the age distribution of likely pathogenic alleles are lower than those of likely benign alleles (Methods §20, Fig. 4a.i), particularly at the ancient tail of the distribution. Frequency-controlled mutation age z-score distributions (Fig. 4a.ii, Methods §21) show that this shift is observed not just in singletons, but across the ultra-rare allele frequency spectrum. Here, we find that likely pathogenic missense variants are depleted for mutations that are older than expected given their frequency (positive z-score bins) and enriched for those that are younger than expected (negative z-score bins). Consistent results are observed when using PrimateAI-3D^66^ to classify variant deleteriousness (Fig. S23).

**Figure 4:**
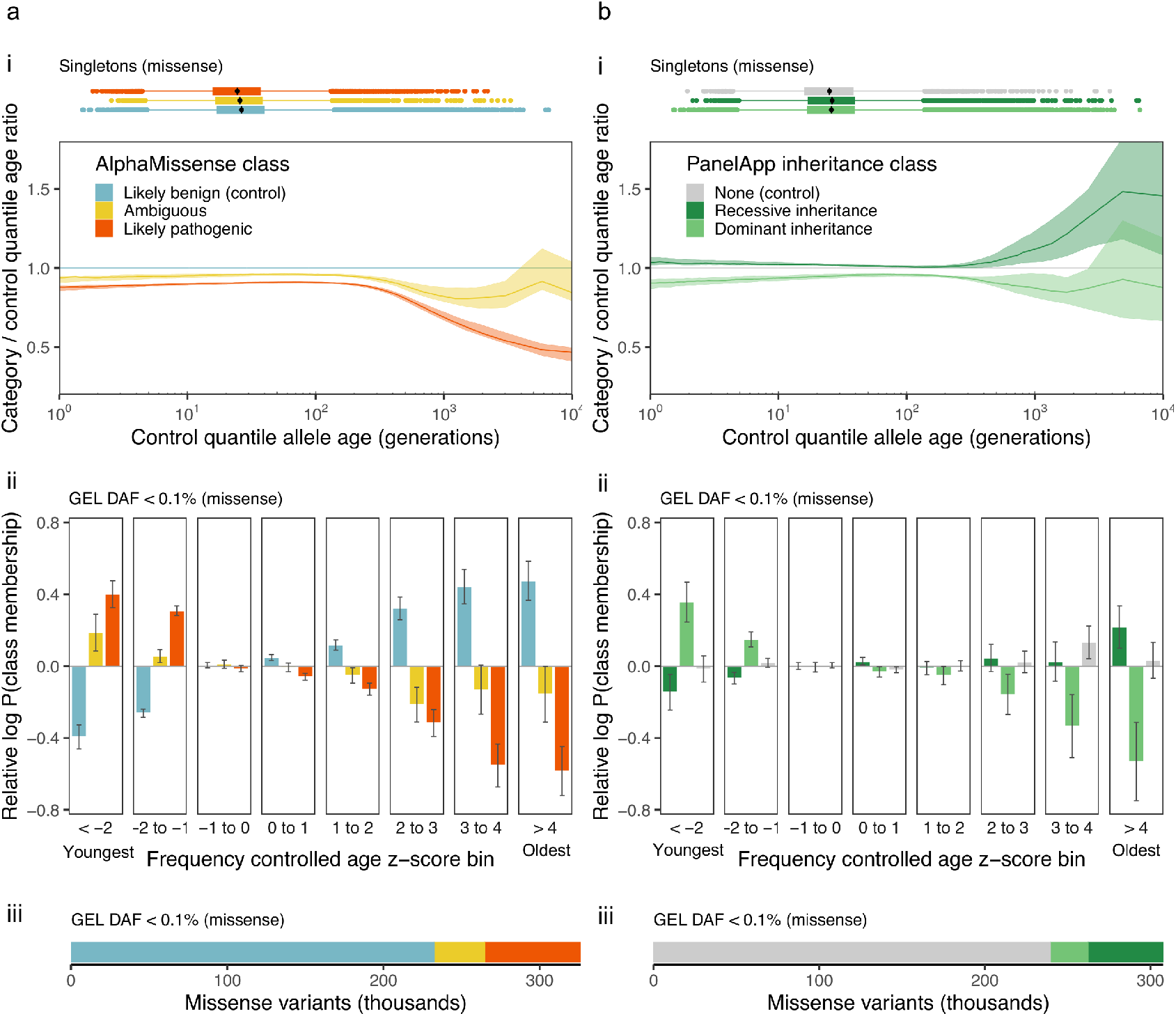
Allele age patterns indicate negative selection within coding regions. **(A) AlphaMis-sense.**(i) Quantile–quantile analysis of singleton missense variants (DAC=1): lines show the ratio of age quantiles for each AlphaMissense category relative to the corresponding quantiles for *likely benign*, with bootstrapped 95% confidence bands. For instance, the median age of *likely pathogenic* missense singletons is about 90% of the median age of *likely benign* missense singletons (which is around 15 generations ago). (ii) For ultra-rare missense variants (GEL DAF < 0.1%), the relative probability (odds ratios) of a variant being classified as *likely pathogenic* or *ambiguous* across frequency-controlled allele-age *z*-score bins (youngest < − 2 to oldest > 4), estimated via logistic regression with 95% confidence intervals. (iii) Counts of ultra-rare missense variants by AlphaMissense class. **(B) PanelApp inheritance**. (i) Quantile–quantile analysis of singleton missense variants contrasting genes annotated as *dominant* or *recessive* against genes with *no disease annotation* (control), with bootstrapped 95% confidence bands. (ii) For ultra-rare missense variants (GEL DAF < 0.1%), the relative probability (odds ratios) of a variant falling in *dominant* or *recessive* disease genes across allele-age *z*-score bins (youngest <− 2 to oldest > 4), from logistic regression with 95% confidence intervals. (iii) Counts of ultra-rare missense variants stratified by PanelApp inheritance class.

Rare recessive deleterious alleles are largely masked from the effects of selection, while dominant alleles are more strongly affected. So, we next investigated whether stratification by clinically annotated modes of inheritance shows similar patterns (Methods §22). Specifically, we compared the distributions of ultra-rare (DAF < 0.1%) missense mutation ages within exons of Mendelian disease-associated genes, stratified into dominant (*n* = 22, 542 variants, 6.9%), recessive (*n* = 45, 502 variants, 14.8%), and non-Mendelian (unannotated; *n* = 239, 638 variants, 73.5%) classes based on expert assessed PanelApp^67^ annotations (Fig. 4b; Methods §15). As predicted, dominant disease genes are depleted for older-than-expected mutations and enriched for younger-than-expected mutations relative to overall missense background (Fig. 4b.i), while recessive disease genes show the opposite pattern.

Extending our analysis beyond missense variants, we examined all coding and non-coding ultrarare variants (DAF < 0.1%; *n* = 7, 552, 186) to assess how mutation age distributions relate to regional selective constraint. Using gnocchi^55^ constraint metrics derived from gnomADv3.1 mapped to 10 kb windows reveals a clear pattern (Fig. 5a). Mutations older than expected for their frequency are significantly enriched in the least constrained regions (gnocchi z < − 4) and depleted in the most constrained regions (gnocchi *z* > 4) (Fig. 5b).

**Figure 5:**
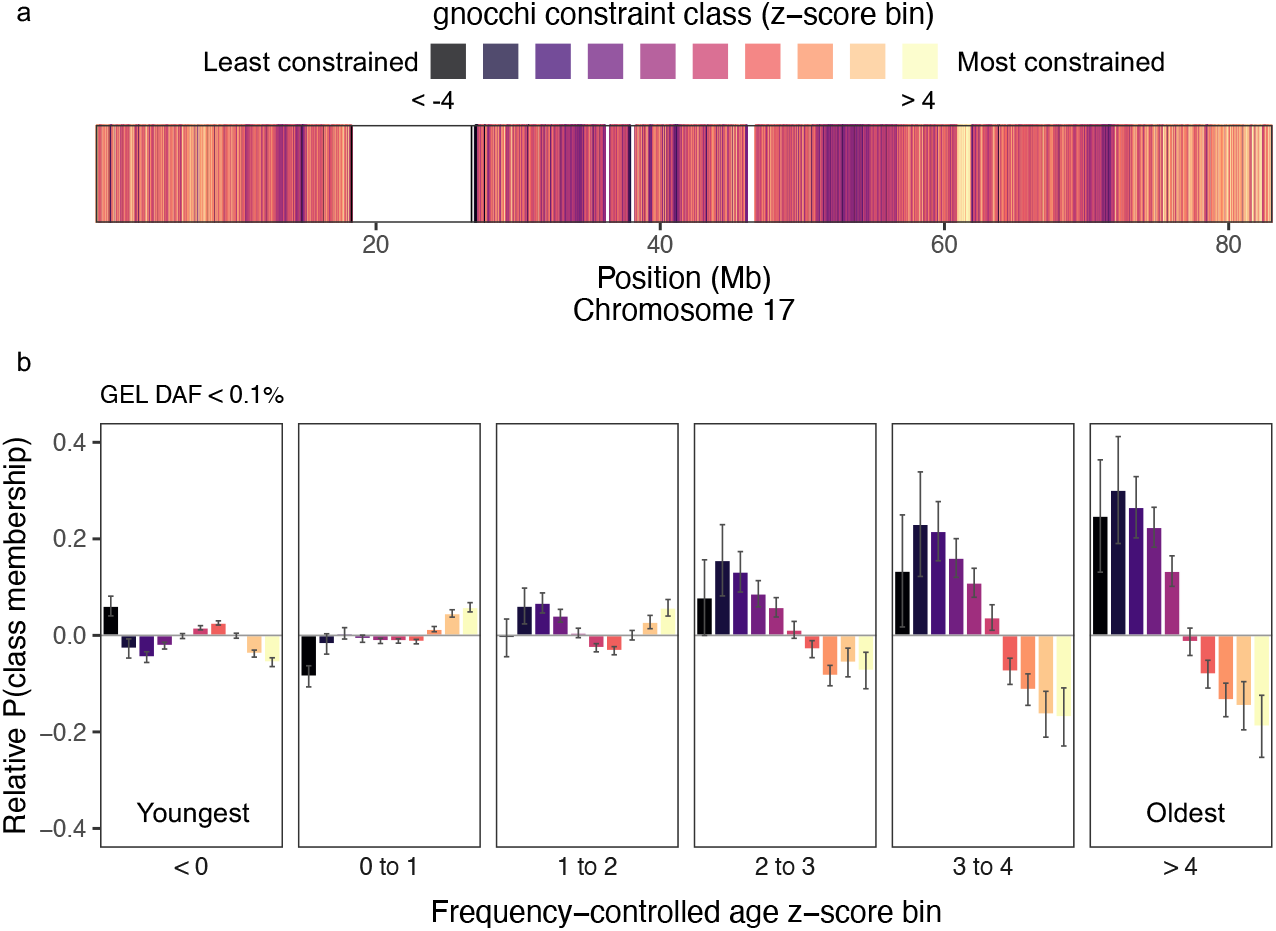
Allele age distributions reflects chromosome-wide patterns of selective constraint. (A) 10kb regions across chromosome 17 coloured by gnocchi constraint z-score bin (< − 4 to > 4). (B) For all ultra-rare (DAF < 0.1%) variants across chromosome 17, the relative probability of a variant appearing within a given constraint z-score bin (10kb region) per frequency-controlled mutation age z-score bin (< 0 to > 4). Relative probabilities (odds ratios) were calculated using logistic regression, with error bars showing 95% confidence intervals.

Collectively, these analyses demonstrate that mutation age distributions capture biologically meaningful signatures of purifying selection across scales—from individual coding variants to broader patterns of regional constraint.

### Age of clinically classified variants

We analysed clinically classified missense or loss-of-function (LoF) variants in the GEL dataset (Methods §22). Among singletons, pathogenic and likely pathogenic (P/LP) variants were significantly younger than both benign/likely benign (B/LB) variants and variants of uncertain significance (VUS) (permutation test all *p* < 0.0001; Tab. S5), consistent with strong purifying selection acting on pathogenic alleles. Notably, P/LP singletons reported exclusively within the 100,000 Genomes Project are younger than those also reported in ClinVar (*p* = 0.0052), supporting the idea that older pathogenic variants are more likely to be shared across independent clinical datasets. We observed the same pattern among clinically classified doubletons: after removing likely recurrent mutations using a genealogical consistency filter (Methods §23, Fig. S24), P/LP variants again exhibit significantly younger ages than both B/LB variants (*p* = 0.0012) and VUS (*p* = 0.0066; Tab. S5).

We next asked whether integrating allele age could identify variants with different evolutionary histories despite being matched by allele frequency, PCA group, and clinical classification (as above, PCA groups are used only for descriptive stratification; age estimation itself is label-free). For this, we focussed on non-recurrent doubletons from the four largest PCA groups in the GEL dataset (NFE, AFR, SAS, AFR; Fig. S25). Within this set, the oldest P/LP variant is estimated to be 339 generations old (95% CI: 49-900). In contrast, we identified six VUS in dominant genes with ages more than an order of magnitude older (4,827-35,340 generations; Tab. S6), which is likely inconsistent with the expectation of strong purifying selection on pathogenic Mendelian variants. One notable example was a missense mutation in *DCGR8*, a gene associated with early-onset nervous system tumours, estimated to be ~ 21,100 generations old (95% CI: 4,900–59,300 generations; Fig. 6a). This variant is present in Denisovans but absent from Neanderthals and lies on a 46 kb (0.06 cM), identity-by-descent (IBD) haplotype shared among two EAS carriers, consistent with proposed dates of Denisovan introgression ~ 50,000 years ago.^68^ Within the EAS PCA group, we further identified 22 VUS at the same allele count (DAC = 2) with ages ranging from 30 and 3,035 generations (geometric mean age = 74 generations). The youngest example was a missense mutation in *SLC2A10* (95% CI: 5–130 generations) located on a 1,129 kb (2.14 cM) shared IBD haplotype (Fig. 6b), indicative of recent shared ancestry among carriers.

**Figure 6:**
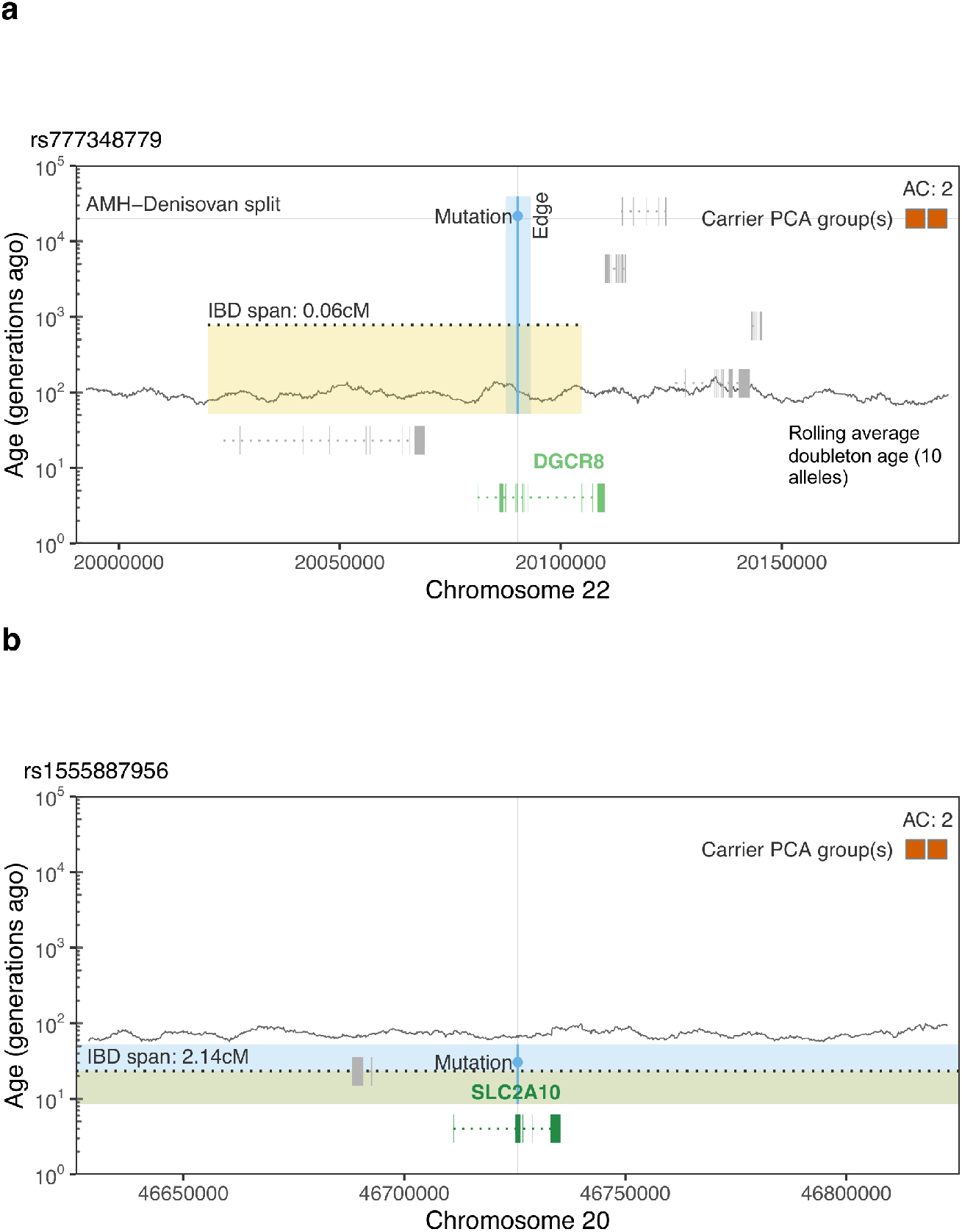
Distinct genealogical contexts underlying two variants of unknown clinical significance (VUS) at the same frequency. (a) Example of EAS doubleton (DAC = 2) VUS with deep inferred age. Allele age is contextualised using approximate dates of the AMH (anatomically modern human) Denisovan split (20,000 generations). (b) Example of EAS doubleton VUS with recent inferred age. Blue shaded areas represent edges upon which the mutations reside, with coordinates bounded by the edge span (x) and the node ages above and below the mutation (y). Orange shaded areas represent IBD haplotypes, defined here as the span where the node below the mutation represents the most recent common ancestor (MRCA) of both carriers, with y-coordinates bounded by the estimated node age and the recombination-based haplotype age (dotted black line). cM: centiMorgans, estimated using HapMapII recombination maps.

Such findings illustrate that even among variants at the same apparent frequency, mutation age and genealogical structure can reveal substantial differences in evolutionary history, offering complementary evidence for variant interpretation.

## Discussion

Molecular dating of phylogenies enabled evolutionary processes to be interpreted on an absolute timescale, revealing patterns of divergence and speciation that were previously inaccessible. In human genomics, however, inference at comparably fine temporal resolution within the recent past has remained challenging, particularly for ultra-rare variants whose interpretation is poorly served by allele frequency alone. Here we show that scalable, uncertainty-aware estimation of allele ages from large ARGs can recover temporal structure within rare variation, providing a complementary perspective on recent mutational history with the potential to inform clinical practice. This was enabled by two key methodological developments. Firstly, we developed an accurate and efficient method to estimate the ages of nodes and mutations in an ARG, which captures uncertainty through a principled variational approximation of their posterior distributions. Secondly, we developed the methods to perform large contiguous ARG topology inference (necessary to accurately date ultra-rare alleles) distributed over a cluster. We then applied these advances to infer highly contiguous ARGs for nearly 50,000 individuals over 14% of the genome, estimating allele ages for 23 million variants. Our analyses show that these ages capture deep biological signals of demography and selection, inaccessible to allele frequency alone. Importantly, because our ARG topology and dating methods do not require any labelling or partitioning of individuals, these results are stable and interpretable under the biased sampling common to modern datasets.

Several factors affect the reliability of these age estimates. First, we assume that each observed allele arose from a single mutational event. While recurrent mutation is relatively rare in humans, it becomes increasingly relevant as analyses focus on ultra-rare variation at very large sample sizes,^69^ particularly in mutational hotspots and at those clinically important loci where multiallelic sites are common. Incorporating calibrated, context-dependent mutation rates^70, 71^ and extending ARG methods to better accommodate multiallelic sites are therefore important directions for future work. Second, our analyses focus primarily on rare, and typically recent, variants. Recent ancestry is often tightly constrained by long shared haplotypes, providing strong information for dating, whereas deeper genealogical structure is more weakly informed by the data. As a consequence, absolute ages in the deep past are sensitive to timescale calibration and to the regularisation required to stabilise root ages, and should be interpreted with appropriate caution. Importantly, relative age comparisons among rare variants are relatively unaffected.

We have shown that age contains information about the likelihood of deleteriousness, and so has potential to inform clinical prioritization of alleles in rare genetic disease diagnosis. The examples in Figs. 3 and 6 give a glimpse of how this might work. This is likely most helpful in narrowing the search: candidate variants who are inferred to be old might be deprioritized. This could greatly increase power in some cases: for instance, an individual who has inherited a small portion of the genome from an undersampled portion of the globe could have a large number of candidate private alleles; however, these would also be inferred to be old (and with high uncertainty), so attention could be directed to the smaller number of confidently recent variants.

While ARG inference has made major leaps in recent years and interest in applications is burgeoning,^35, 72–80^ there remains considerable work to do before it becomes a routine component of clinical and statistical genetics pipelines. Scalability is one axis: even with distributed implementations, WGS data at biobank scale remain a substantial challenge. Another is uncertainty: the credible intervals reported by tsdate are expected to be well-calibrated *given* the ARG, but do not incorporate uncertainty in ARG topology itself. At present, Bayesian methods that sample from a posterior on ARGs^29, 31^ are limited to much smaller sample sizes. Hybrid inference schemes that reconstruct the tightly constrained recent past using non-parametric, heuristic approaches and then apply model-based methods to the deep past are therefore attractive.^81^ As ARG methods mature, it will be important to develop systematic approaches to quality control and evaluation. Standardised simulations,^82–84^ richer comparison metrics that quantify differences between inferred and true genealogies,^23, 34^ and new visualisation tools^85–87^ are important steps. Here we evaluated our inferred ARGs by comparing tsdate ages and tree structures to multiple independent lines of evidence. A systematic collection of such benchmarks across multiple species would be a valuable resource for the field, ensuring that methods are robust to the vagaries of real data.

Inferred ages of common alleles have found diverse applications.^63, 64, 88–93^ Our work shows that accurate and uncertainty-aware allele ages can now be obtained for ultra-rare alleles. These ages provide a coherent temporal interpretation of ultra-rare variation and link genealogical structure with functional and clinical consequences across a heterogeneous cohort. As sequencing efforts expand to larger and more diverse populations^11, 94–96^ and genealogical methods continue to develop, allele age is likely to become an important statistic in human genomics, informing evolutionary, functional, and clinical analyses. By avoiding reliance on population labels, allele age provides a stable, transferable measure of variation that is well suited to large, ancestrally mixed datasets and to cross-cohort synthesis of rare variant information.

## Online methods

### 1 Variational dating algorithm

We develop an efficient and accurate method of estimating the ages of each ancestral node in an inferred ARG, using a Bayesian message-passing algorithm (Fig. 1). Each edge in an ARG represents the inheritance of a particular segment of genome by a *child* node from another, more ancestral *parent* node. The *area* of an edge is its genomic span in base pairs, multiplied by the time between parent and child nodes in generations. We assume that the number of mutations falling on this edge is Poisson distributed with a mean given by its area multiplied by the mutation rate. The conjugate prior to the Poisson is the Gamma distribution, and therefore if the edge lengths were unconstrained, a Gamma prior on each edge length would lead to a Gamma posterior. ARGs impose strong constraints on node ages,^41^ however, and the exact posterior of node times is therefore intractable. We use Expectation Propagation^43^ (EP) to compute an efficient and principled variational approximation of the posterior node times in an ARG.

More precisely, we place exponential priors on node times and obtain a Gamma distribution for each node that approximates its marginal posterior. Properties of exponential families^97^ allow us to decompose the marginal posterior of each node time into a product of per-edge log-linear “factors” that are refined iteratively by EP updates (Fig. S26). For a given edge, EP proceeds by removing the associated factors from the parent and child variational posteriors, forming a bivariate “cavity” distribution (Fig. S26B, blue contours). The cavity is multiplied by the Poisson likelihood for the edge, forming the target or “surrogate” distribution (Fig S26B, black contours). Finally, the variational posteriors for the parent and child are updated to match the mean and variance of the surrogate (Fig S26B, red contours). A key factor in the computational efficiency of this algorithm is that the moments of the surrogate distribution are expressed in terms of hypergeometric functions and efficiently updated using saddlepoint approximations.^98^ A single round of EP updates requires traversing all edges in the ARG from leaves-to-roots and then roots-to-leaves. Expectation propagation is not guaranteed to converge in general, but in practice we find rapid convergence to accurate variational posteriors for node ages (Figs. S3, S4).

Formally, let 𝒱 and ℰ be the sets of nodes and edges in the ARG, respectively, and let *t*_*i*_ be the age of the *i*th node measured as generations in the past. We know the ages of a subset of sampled nodes 𝒮, and wish to infer ages for the remaining set of internal nodes 𝒩. For simplicity, assume that a given pair of nodes is connected by at most one edge and use *ij* ∈ ℰ to denote the edge connecting parent *i* to child *j* (i.e., *i* is ancestral to *j* over a single contiguous genomic segment). This assumption is for notational convenience only, and the underlying model can accommodate an arbitrary number of edges between a given ancestor and descendant. Each edge has two dimensions: time *t*_*i*_ − *t*_*j*_ measured in generations, and genomic span *s*_*ij*_ measured in base pairs. If the per-generation, per-base probability of mutation is a constant *µ* ≪ 1 and independent between bases, then the probability of *y*_*ij*_ mutations occurring along the edge is Poisson,

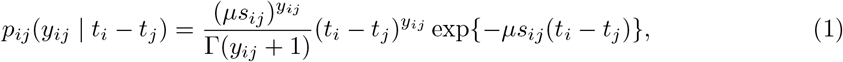

subject to the constraint *t*_*j*_ < *t*_*i*_ for all *ij* ∈ ℰ. Let *p*(***t*** | ***η***) be a joint prior distribution on the ages of the internal nodes, with hyperparameters *η*. The posterior of node ages is then,

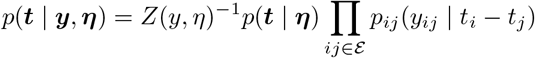

where *Z*(*y, η*) is the appropriate normalizing constant.

Exact posterior inference of node ages is not tractable because *Z* is an integral over a high-dimensional polytope. Instead, we approximate the true posterior *p* by a product distribution *q*_***θ***_,

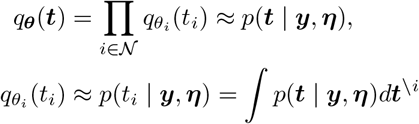

where 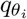 is an approximate Gamma marginal for the *i*th node with variational parameters *θ*_*i*_, and ***t***^\*i*^ is the vector of node ages without the *i*th node. The advantage of this approximation scheme is that the integration problem becomes an optimization problem: find the values of ***θ*** that produce the best approximation according to some distributional measure of discrepancy, in our case the Kullback-Leibler (KL) divergence. Since Gamma distributions form an exponential family, minimizing KL divergence is equivalent to moment matching of sufficient statistics *ϕ*(*t*) = {*t*, log *t*}:

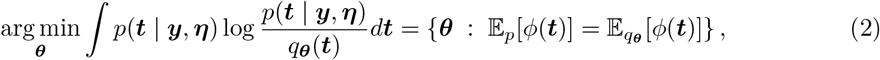

where 𝔼_*p*_[·] denotes expectation with respect to *p*(***t*** | ***y, η***), and *ϕ*(***t***) is the concatenation of Gamma sufficient statistics for each node.

The minimization in Equation (2) is still intractable, as 𝔼_*p*_[*ϕ*(***t***)] also involves a high-dimensional integral. We use “expectation propagation”^43^ to approximate the global optimization problem with a series of tractable local problems. Because the approximate marginals 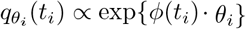 are log-linear in the variational parameters *θ*_*i*_ (up to a normalizing constant), each may be factorized into

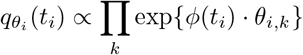

subject to ∑_*k*_ *θ*_*i,k*_ = *θ*_*i*_. Note that the individual factors have the same form as the original distribution, but are not required to have a finite integral (as long as their product can be normalized). Therefore, we reparameterize the approximate joint posterior as

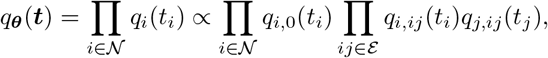

with the shorthand *q*_*i,ij*_ = exp *ϕ*(*t*_*i*_) · *θ*_*i,ij*_, and requiring that *θ*_*i*_ = *θ*_*i*,0_ + ∑_*j*:*ij* ∈ *ℰ*_*θ*_*ij*_ + ∑_*j*:*ji* ∈ℰ_*θ*_*ji*_ defines a valid (normalizable) distribution for all *i*. This aligns the form of the posterior to match that of the likelihood (1), allowing us to work with terms grouped by edge, not by node. In words, the variational marginals for each node are split into “messages” that correspond to the prior (*q*_*i*,0_) and edges (*q*_*i,ij*_), and each has the same form as the unnormalized posterior.

Expectation propagation works by iteratively refining the messages, one at a time. For edge *ij*, the “cavity” distribution is the approximation without the *ij*-th factor,

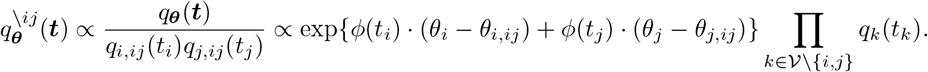

The “surrogate” is the product of the cavity and the Poisson edge factor from the exact unnormalized posterior,

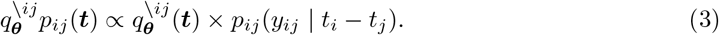

Intuitively, the surrogate is an approximation of the posterior that is exact locally (for the edge being refined) and approximate elsewhere. The variational parameters can be updated by matching moments (minimizing KL divergence) against the surrogate: given a current set of parameters ***θ***, we wish to find an updated set of parameters ***θ***^′^ such that

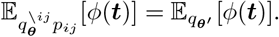

The crucial difference with the global problem (2) is that the surrogate is *mostly factorized*, so that 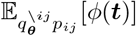 and 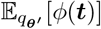 only differ for nodes *i* and *j* (i.e., for *ϕ*(*t*_*i*_) and *ϕ*(*t*_*j*_)). Thus, the only variational marginals that need modification are *q*_*i*_ and *q*_*j*_. The messages are updated by dividing the new approximate posterior by the cavity (equivalent to subtracting natural parameters), so the only changes between ***θ*** and ***θ***^′^ are that 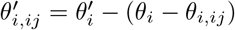 and 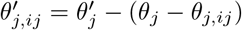.

What remains is to calculate moments of the surrogate (3). Rather than the sufficient statistics 𝔼[*t*_*i*_] and 𝔼[log *t*_*i*_], it turns out to be more numerically stable to match 𝔼[*t*_*i*_] and 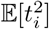, and in practice gives a nearly equivalent fit. To find the moments, first note that the density of the surrogate is proportional to

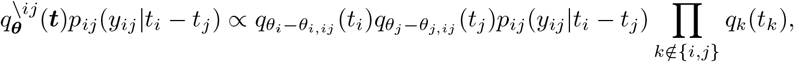

and so terms related to *t*_*k*_ with *k* ∉ {*i, j*} do not appear. At this point, it is useful to switch to the canonical parametrization of the Gamma, so that *q*_*i*_(*t*_*i*_) ∝ exp{−*t*_*i*_*β*_*i*_ + (*α*_*i*_ − 1) log *t*_*i*_}. The message is then parameterized as *q*_*i,ij*_ ∝ exp{−*t*_*i*_*β*_*i,ij*_ + *γ*_*i,ij*_ log *t*_*i*_}, and the cavity parameters as 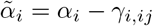 and 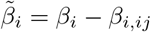, so that

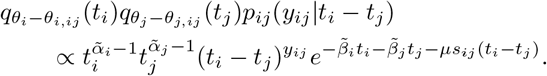

Defining

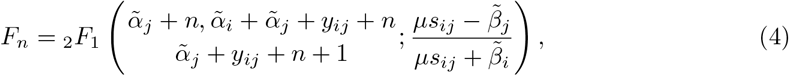

where _2_*F*_1_ is the Gauss hypergeometric function, we can write the first two moments as

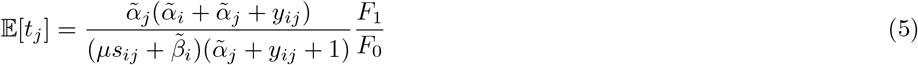

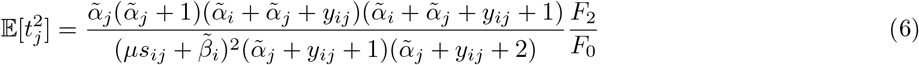

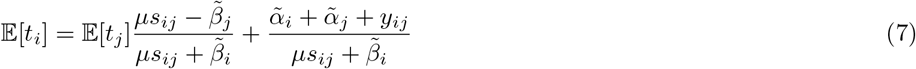

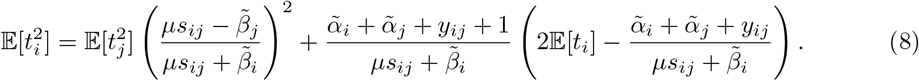

Or, when node *j* is a contemporary sample (*t*_*j*_ = 0),

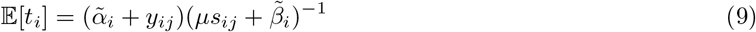

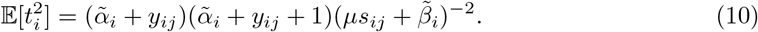

The full derivation of Equations (4) through (10) is given in SI §1. In practice, we use a fast and numerically stable saddlepoint approximation^98^ to evaluate the hypergeometric series (SI §2). Equivalent moments are derived for noncontemporary samples in SI §3. The updated natural parameters are obtained from the moments by

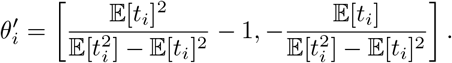

If necessary, 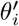 is projected onto the feasible range (such that the marginals can be normalized).

Expectation propagation converges to a fixed point that is often very close to the global minimum of (2).^99^ We find that iterating over edges in increasing then decreasing time order (i.e., from leaves to roots and then roots to leaves) provides the most rapid convergence, although in theory the updates could be done in parallel. Finally, variational approximations to the posteriors of mutation ages are calculated from those of node ages after convergence (SI §4).

### 2 Timescale calibration via root-to-leaf distance

The expectation propagation update described in Methods §1 is local in scope; taking a noisy observation of the area of an edge and updating our posterior belief regarding the ages of the two attached nodes. However, the assumption that mutation follows a Poisson process implies a strong global property: that the number of mutations in a given time interval is equal in expectation to the sum of all edge areas intersecting that interval (multiplied by the per-base mutation rate). Therefore, after the per-node posteriors have been obtained by expectation propagation we rescale time globally to enforce an approximate molecular clock.

Unfortunately, we cannot expect this property to hold for ARGs produced by tsinfer. The reason is that tsinfer encodes uncertainty about the topology of genealogical trees via polytomies. If the true genealogies are actually binary across all positions, then these “spurious” polytomies will create an excess of edge area relative to mutational density, which will introduce bias into estimates of node ages.

Instead, we enforce a match between the average number of mutations carried by each sample in a given time interval and the length of the interval multiplied by the mutation rate. Intuitively, the number of neutral mutations carried by a sample measures the average distance between it and the roots, a quantity that is not impacted by spurious polytomies. Specifically, we find a piecewise-constant, monotonic transformation of time that results in an ARG satisfying this constraint. To do so, we use equally-spaced quantiles of mutation ages to determine a time discretization, use the posterior means of node ages to calculate edge areas, and integrate over the ages of mutations on each edge.

A second variational approximation is used to transform the posteriors themselves: we transform a lower and an upper quantile of each posterior, then find the gamma distribution that matches these transformed quantiles. This strategy only requires a binary search over time intervals and a few Newton iterations, and is far more efficient than matching mean and variance (that would be calculated by piecewise-constant integration). The algorithm requires a single pass over edges and a single pass over nodes, and is described in detail in SI §7.

This approach is inspired by the rescaling algorithm implemented in SINGER^31^ and POLE-GON,^42^ that matches observed and expected numbers of mutations rather than the average count per sample. To assess the impact of spurious polytomies, we implemented both rescaling approaches in tsdate, and applied them to a 100 Mb simulated ARG containing 20,000 samples after collapsing branches without mutational support. When the ARG contains strictly binary trees, both approaches are similarly unbiased (Fig S27). However, when spurious polytomies are present, rescaling based on absolute mutation counts and edge area introduces a strong, time-dependent bias in node ages (Fig S27, “area rescaling”), that is largely eliminated by weighting mutations by the number of carriers (Fig S27, “path rescaling”).

### 3 Regularization of root ages

Information about the ages of nodes comes from noisy measurements of edge lengths—mutational density—that are propagated up the ARG from a fixed reference point (the sample ages). Typically, the oldest ancestral segments will each have very short spans, because of the cumulative action of recombination. Mutational density is sparse on these ancient haplotypes, and there is relatively little ancestral material that can constrain their ages. As a consequence, the posteriors of the oldest nodes typically have extremely high variance, and this is exacerbated in the variational approximations by the local nature of expectation propagation. Thus, we regularize the ages of “ultimate” roots (the nodes with no parents) via an exponential prior. This scheme acts as a soft constraint on the maximum height of the ARG.

Specifically, if the *i*th node has no parents, then it is given the prior *p*(*t*_*i*_ | *η*) ∝ exp { − *ηt*_*i*_ }. As this is a special case of the Gamma distribution, the prior incorporates directly into the variational posterior as *α*_*i*,0_ = 0 and *β*_*i*,0_ = *η*. Rather than specify *η a priori*, we fit it via expectation maximization (EM) at the end of each round of EP updates. Let *δ*_*i*_ be a binary indicator of whether node *i* is without parents, and *q*_*i*,\0_ be the variational posterior without the contribution of the prior (i.e., *α*_*i*,\0_ = *α*_*i*_ and *β*_*i*,\0_ = *β*_*i*_ − (1 − *δ*_*i*_)*η*). The surrogate model is Π_*i*_ *q*_*i*,\0_(*t*_*i*_) exp(−*ηt*_*i*_*δ*_*i*_). The associated EM objective for *η* is 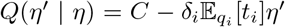 where *C* is a constant that does not depend on *η*, leading to the update

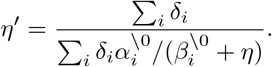

For the applications in this paper involving very recent mutations, this simple regularization scheme works well while adding little additional overhead. However, it may be an oversimplification if the root ages themselves are of interest, especially if a large heterogeneous sequence is being dated. In this case, the regularization scheme may be generalized to a mixture of Gamma distributions, at the expense of computational cost (described in detail in SI §6).

### 4 Ambiguous singleton phasing

ARG inference methods require input genotypes to be phased, most often achieved by statistical methods.^100, 101^ While phasing tends to be quite accurate for common variants and ARG inference methods robust to switch errors in simulations,^102^ singletons require special consideration and are often omitted from analysis. Support for phasing singletons has only recently been introduced in SHAPEIT5,^101^ and is based on choosing the shorter of the two possible carrier haplotypes, as this is likely to be the older copy (and thus the more likely to have mutated at a given base). A similar rationale is used by Runtc,^16, 103^ where ancestor age is estimated directly from shared haplotype length.

Singleton mutations have no impact on the tree topologies produced by tsinfer. So, the analogous procedure with an inferred but undated ARG is to choose the shorter of the two edges that connect to the carrier individual as the correct phase. However, the choice of phase determines the allocation of mutations across terminal edges, and may therefore impact the estimated ages of ancestors and of the singleton mutations themselves. Thus, we adapted the model described in Methods §1 to integrate over both possible phases for each singleton mutation. By doing so, we propagate uncertainty about choice of phase into the variational posteriors for the ages of nodes and mutations, and therefore leverage information from the entire graph rather than just singleton edges. This modification requires splitting the edges above each leaf so that any given individual has at most two immediate ancestors (nodes) over the span of every attached edge. See the Supplement (SI §5) for a detailed description of the “phase-agnostic” EP algorithm.

In Fig. S28, we show the impact of phasing for singleton age estimates in a 100Mb inferred ARG of 20,000 diploids, simulated from an equilibrium population with msprime. Choosing singleton phase based on inferred edge span largely mitigates error due to incorrect phase, relative to a worst-case scenario where phase is randomly assigned. The phase-agnostic algorithm marginally improves upon this shortest-edge heuristic. See also Fig. S9 for a benchmark on 1KGP data.

### 5 Accuracy of mutation age estimates from inferred ARGs

To measure the accuracy of variational posteriors for mutation age under realistic inferences, we simulated ARGs with 15,000 diploids sampled uniformly across populations using stdpopsim^84^ (version 0.2.1) and msprime.^104^ The simulated ARGs are modelled after 40Mb of human chromosome 17, using the HapMapII recombination map^105^ and four different demographic models: `AmericanAdmixture_4B18`,^106^ `OutOfAfricaArchaicAdmixture_5R19`^107^ `OutOfAfrica_3G09`,^108^ and `Zigzag_1S14`.^109^ We inferred ARG topologies using tsinfer with default paramters, and used the tsdate VG (new) and IO (previous) algorithms to date mutations. We measured bias and RMSE in estimated mutation age for both methods, across frequencies of very rare mutations (< 0.001 in global frequency; Fig. S1, Tab. S1).

To measure the calibration of the posteriors produced by the VG algorithm, we calculated empirical versus observed coverage of true mutation ages for posterior intervals of various widths. Because we expect ARG topology inference by tsinfer to introduce errors that are not modelled by the VG algorithm, we dated both true and inferred ARGs with the expectation that posteriors from the latter would underestimate uncertainty (Fig. S4).

### 6 Dating algorithm benchmarking

To assess the performance of the variational gamma algorithm versus the previous version of tsdate (“inside-outside”),^41^ we used an ARG of human chromosome 22 from a large realistic simulation.^45^ We randomly subsampled the ARG (by simplification^110^) to sample sizes ranging from 10 to 1 million human chromosomes, for three replicates at each sample size. We also benchmarked POLEGON^42^ (version 0.1.3, commit 0c088b3). We used the default settings for POLEGON (100 MCMC samples with a thinning interval of 10, followed by 3 rescaling iterations).

For all three methods, we measured wall time using the `time` module of the python standard library, and peak memory usage using `memray` for python code. For POLEGON, the additional memory usage of the compiled binary used for MCMC sampling was profiled via GNU Time, and peak memory usage was taken as the maximum over the parent python process and subprocess. Wall time and peak memory were measured separately, to avoid additional overhead from memory profiling. Finally, we measure accuracy as the root mean squared error in log_10_ estimated node age.

### 7 1,000 Genomes Project ARG inference

We downloaded the phased 1000 Genomes dataset from https://ftp.1000genomes.ebi.ac.uk/vol1/ftp/data_collections/1000G_2504_high_coverage/working/20220422_3202_phased_SNV_INDEL_SV/. Notably, singletons are absent from this resource.^49^ We converted the VCF to VCF Zarr^111^ format using bio2zarr version 0.1.5, and used ancestral state information from Paten et al.^112^ to polarise alleles. We performed ARG inference using tsinfer following the same approach as the GEL dataset (Methods §14). Details of the regions inferred are given in Tab. S7. All 3202 samples were included.

We extracted singletons from the unphased high-coverage 1000G data (https://ftp.1000genomes.ebi.ac.uk/vol1/ftp/data_collections/1000G_2504_high_coverage/working/20201028_3202_raw_GT_with_annot/) using bcftools.^113^ We added SNPs to the inferred ARGs if they were biallelic, passed quality control, had no missing genotypes and had a derived allele present in a single sample. We then used tsdate 0.2.4 to phase singletons and date the ARGs using a mutation rate of 1.29 × 10^−8^. Summary statistics of the resulting ARGs are given in Tab. S7.

To evaluate sensitivity to sample size, we also inferred ARGs using tsinfer+tsdate on the long arm of chromosome 20 for randomly-sampled sets of 100, 300 and 1500 samples, ensuring that the proportion of every super-population in each sample was conserved. Results are shown in Fig. S8.

To test the effectiveness of tsdate’s phase-agnostic singleton dating, we identified mutations that are singletons in the 1500 sample subset but of higher frequency in the full 3202 sample dataset. Results are shown in Fig. S9.

### 8 Comparison to ancient DNA dates

We used ancient DNA data from the 1240K Allen Ancient DNA Resource (AADR)^50^ v62.0 for Fig S5. We converted genome data for chromosome 20 to VCF using convertf^114^ and PLINK,^115^ then lifted over from hg19 to hg38 using the bcftools^113^ liftover plugin together with the chain file from https://hgdownload.soe.ucsc.edu/goldenPath/hg19/liftOver/. We set ancient DNA sample ages using the AADR field “Date mean in BP in years before 1950 CE” and minimum site ages from the oldest aDNA sample containing the derived allele at that site. We compared these minimum site ages to the time of node ancestral to the mutation in the inferred 1KGP ARGs (Methods §7).

We define a site as consistent with the aDNA date if the node time converted to years is older than the minimum aDNA site age, using both a conservative assumption of an average human generation of time of 25 years, and a more lenient assumption of 27 years.^116^

### 9 Comparison to PHLASH

We compared the inferred times of human effective population size (*N*_*e*_) changes as derived from the 1KGP inferred ARGs (Methods §7) with independent estimates from a recent study.^51^ Fig S6 shows data from Fig. 3a of Terhorst^51^ (obtained from https://github.com/jthlab/phlash_paper/blob/master/data/fig7b.csv) along with the inverse of the instantaneous coalescence rate (equivalent to *N*_*e*_) computed from chromosome 20 of the inferred ARGs using the tskit `pair coalescence rates` function. Coalescent times are comparable, as both methods use the same mutation rate (1.29 × 10^−8^). For ease of display we focus on three continental groupings (AFR, EUR and EAS) present both in the PHLASH analysis and in our 1KGP inferred ARGs.

### 10 Dating the 17q21.31 inversion

To estimate the age of the 17q21.31 inversion with tsinfer+tsdate, we extracted the identifiers and associated H1 and H2 alleles of 21 marker SNPs from Table 2 of Donnelly et al.,^117^ where H1 and H2 correspond to non-carriers and carriers of the inversion, respectively. We excluded the SNP labeled `E_TAUIVS11_10` because we could not confirm its hg38 genomic coordinates. For the remaining 20 SNPs, we used their RSids to retrieve hg38-mapped positions from https://myvariant.info/.Of these, 18 were present in the inferred 1KGP ARG (Methods §7); three additional markers were excluded because the derived or ancestral allele in the ARG did not match either the H1 or H2 allele, rendering carrier status ambiguous. We assigned a haploid sample in the ARG to the H1 or H2 group if it carried the H1 or H2 allele, respectively, at all 15 retained marker sites. We computed cross-coalescence rates between the H1 and H2 groups using the `pair_coalescence_rates` function in tskit, with 200 evenly spaced, non-overlapping genomic windows from ~45.6–46.2 Mb and 50 log-spaced time windows from approximately 10^3^ to 3 × 10^5^ generations ago.

Relate-based^37^ upper-bound estimates for the age of the inversion by genomic region were obtained from https://github.com/a-ignatieva/dolores-paper/blob/main/1KGP/data, as published by Ignatieva et al.^34^ We lifted over genomic region coordinates from hg19 to hg38 using liftover^118^ (v1.3.2) with a chain file from https://hgdownload.soe.ucsc.edu/goldenPath/hg19/liftOver/, determining that the total region used for inversion age inference was from ~45.6–46.2 Mb. The results are compared with tsinfer+tsdate estimates in Fig. S7.

### 11 Comparison to published allele age estimates

To compare allele age estimates produced by tsinfer+tsdate on real data with those of other methods, we assembled a dataset of four previously published estimates for 1KGP data. We developed a Snakemake^119^ pipeline to run tsdate on chromosome 20 of the 1KGP ARGs (Methods §7) and to perform the necessary data manipulation over the existing datasets. We ran tsdate using a mutation rate of *µ* = 1.29 × 10^−8^.

We obtained allele age estimates from four external methods (Relate, CoalNN, SINGER, and GEVA) from publicly available resources. We downloaded Relate^37^ data from https://zenodo.org/records/3234689, where results are provided separately for each population in the 1KGP; we calculated the age of each mutation as a population size weighted average of the branch midpoint estimates for each population. Site positions were lifted over from hg19 to hg38 using liftover^118^ (version 1.3.2) with a chain file obtained from https://hgdownload.soe.ucsc.edu/goldenPath/hg19/liftOver/. We obtained CoalNN^18^ allele age data for each super population in the 1KGP from https://github.com/PalamaraLab/coalNN_data/releases/tag/v1.0; we applied the same population size weighted averaging approach across super populations to obtain a single estimate per mutation. We downloaded ARGs estimated from 200 African individuals by SINGER^31^ from https://zenodo.org/records/10467509. The dataset consists of 100 ARGs sampled from the posterior, and we extraced allele ages from the first sample. Mean allele ages from GEVA^17^ were downloaded from https://human.genome.dating/, and site positions were lifted over from hg19 to hg38 using the same procedure as for Relate.

After aggregating allele age estimates, we identified a subset of 162,556 sites that were present in all datasets and had consistent derived and ancestral allele assignments. No allele-based filtering was applied to SINGER because nucleotide alleles are not specified in the ARG; ancestral and derived alleles are simply encoded as 0 and 1, respectively. The results are shown in Fig. S10.

### 12 Simulating sampling bias

To perform simulations modelling the sampling bias in the GEL dataset we used stdpopsim^84^ (version 0.2.1) to run forward-time SLiM^120, 121^ (version 4.3) simulations of a 40Mb segment of the human genome, chosen to match the longest inference region (17q4, Tab. S2), and using the HapMapII recombination map.^105^ We imposed selection by simulating neutral and deleterious mutations, applying the `Mixed_K23` distribution of fitness effects (DFE)^122^ to exonic regions within the chosen segment, and used the `OutOfAfrica_3G09` model,^108^ which describes a history in which European, East Asian, and African populations evolved separately with limited migration over the past 30-60,000 years.

For balanced sampling, we took an equal number of individuals from each simulated population; in the unbalanced case we sampled from these populations in a ratio of 95:1:5, which approximately matches sampling proportions in the GEL dataset. Note that despite the simulation terminating with a global population of only ~ 100,000 individuals, the majority of variants still have a DAF < 0.1% (with 76% of variants counted as ultra-rare in the unbalanced case versus 78% in the balanced case). We then ran tsinfer+tsdate on the simulated data, using default parameters.

Mutation ages can only be estimated on the basis of node times above and below a mutation. For a fair comparison, we therefore set the “true” age of each mutation as the midpoint of the simulated dates of its child and parent nodes, comparing this to the inferred midpoint age taken from the topologically unconstrained posterior mean node times provided by tsdate. Mutation ages were only used from sites without recurrent mutations (corresponding to 98.2% or 1,419,535 sites in the simulation with balanced sampling and 98.7% or 1,000,412 sites in the simulation with unbalanced sampling).

The results are shown in Fig S11, and additional analysis is provided in SI §9.

### 13 Distributed ARG inference

Briefly, tsinfer operates in three phases: first, we generate putative ancestral haplotypes from the genotype matrix using heuristics; then we infer how ancestors descend from older ancestors using the Li and Stephens^123^ (LS) Hidden Markov Model (HMM); then we infer how samples descend from the inferred ancestors, using the same LS HMM machinery. Making inference of the large contiguous ARGs fundamental to this study feasible with the computing resources available required two key updates to tsinfer.

A significant bottleneck in earlier versions of tsinfer occurs during the second phase, in which we infer genetic inheritance among inferred ancestors. Groups of ancestors for which the LS HMM was conducted in parallel were determined only by their temporal ordering. This resulted in many small sets of ancestors, greatly limiting parallelism. As ancestors that do not overlap genomically are independent and can be placed in the same matching group this approach is too conservative. Tsinfer version 0.4 uses a linesweep algorithm to detect overlaps, encoding the result in a directed acyclic graph that represents the minimal constraints on parallelism. Parallel ancestor groups are then formed by progressively removing ancestors with no inbound edges until no ancestors remain.

The second key advance in tsinfer version 0.4 is to allow matching under the LS HMM for these groups of independent ancestors (and samples) to be distributed over a cluster. The overall inference job is arranged into a directed graph of tasks which output either partial HMM results or intermediate tree sequence files. The core count and memory requirements for each job are dynamically set from the number and length of ancestors in each group, allowing high cluster utilisation. In addition to enabling greater parallelism, this architecture also provides resilience to the failures that are inevitable in a busy compute cluster, along with checkpointing and mid-run analysis and monitoring. We developed a Snakemake^119^ pipeline to fully orchestrate this process, allowing large-scale tsinfer inference to be performed on a range of job schedulers.

### 14 Genomics England dataset

We inferred ARGs for Genomics England’s `aggv2` dataset. This dataset contains 722 million variants for 78,195 rare disease and cancer participants recruited as part of the 100,000 Genomes Project.^19, 124^ 69 of these participants had withdrawn consent for the use of their data in research as per the 100,000 Genomes Project main programme release 17 and were therefore removed from consideration. We excluded a further 30,591 individuals who were identified as parents of other participants and phased by direct transmission information^54^ for two reasons. Firstly, tsinfer does not currently support the use of pedigree information to enforce ancestral relationships between individuals, meaning that the haploid genomes of parents and children would always be contemporaries in the ARG. Secondly, the probands in each pedigree are of primary interest for genetic diagnosis purposes, such that removing them in favour of their parents would limit the clinical applications of inferred mutation ages. The remaining subset of 47,535 putatively unrelated individuals was used for ARG inference.

We converted the `aggv2` VCF data to VCF-Zarr format using bio2zarr (version 0.0.8).^111^ We then filtered as above to 47,535 diploid samples, and added ancestral state information.^112^ Sites were then subject to a series of quality control filters which removed sites at duplicated positions, non-SNP variants, non-biallelic sites, and sites with either missing or low quality ancestral state assignments. Additionally, we implemented allele frequency-based filtering, requiring a minimum of 2 derived and 3 ancestral alleles at each site. To avoid problematic flanking regions we applied a sliding window approach with a window size of 100,000 base pairs, requiring that a window-average density of 50 sites per kilobase needed to be met before sites would be used from the 5’ and 3’ ends. We then performed distributed ARG inference using tsinfer, with ancestor truncation disabled, path compression enabled and with a mismatch ratio of zero (ensuring that there is one mutation per site).^38, 41^ We then fully simplified the resulting tree sequences, and used tsdate (version 0.2.1) to split disjoint nodes and then assign ages to nodes and mutations using the variational gamma method with a SNP mutation rate of 1.29 × 10^−8^ per base pair per generation.^82, 125, 126^

As singleton variants were excluded from the phased `aggv2`, we extracted all biallelic, singleton SNPs and the corresponding individual in which they occur from the unphased `aggv2` VCF files using bcftools v1.12, applying the same site-level quality control filters as used by Shi et al.^54^ to construct the phased dataset. We then appended unphased singleton mutations in their derived state to the terminal branches. We re-ran tsdate (version 0.2.1) to phase singleton mutations and date the updated tree sequences using the same parameters as specified above. Allele ages were taken as the posterior mean times inferred using the variational gamma method.

We applied this pipeline to the 15 genome regions detailed in Tab. S2. Statistics for the resulting 15 ARGs are listed in Tab. S4 and the CPU resources required detailed in Tab. S3. The combined dataset of 11,445,983 sites used for primary inference with tsinfer and the 11,778,293 singletons (23 could not be dated due to numerical instability during posterior rescaling) added by tsdate yielded a final dataset of 23,224,276 allele ages.

### 15 Annotation of dated mutations

To characterize how estimated mutation ages vary across diverse genomic contexts, we annotated all variants in the GEL dated ARGs using multiple external resources.

Publicly available ancestry-stratified allele frequencies were obtained from gnomAD v4.1 (genomes and exomes; https://gnomad.broadinstitute.org/data#v4) and GenomeAsia100k^60^ (genomes; https://browser.genomeasia100k.org/#tid=download) as composite VCF files. For each variant overlapping the GEL ARG, alleles were polarised according to the ancestral and derived states in the ARG. Where the external datasets reported the opposite reference/alternate encoding, frequencies were flipped accordingly to yield derived allele frequencies consistent with the GEL ancestral polarisation.

Variant consequences within canonical gene transcripts were predicted using ENSEMBL VEP^127^ v111. AlphaMissense^65^ scores for all missense variants in canonical transcripts were extracted from public Google Cloud buckets (https://console.cloud.google.com/storage/browser/_details/dm_alphamissense/AlphaMissense_hg38.tsv.gz) and, following AlphaMissense nomenclature, classified as ‘likely benign’, ‘ambiguous’, or ‘likely pathogenic’ using default thresholds. Variants were also annotated with PrimateAI-3D^66^ scores downloaded from https://primad.basespace.illumina.com/download, under controlled access for research-only use. Scores > 0.8 were classified as ‘likely pathogenic’, < 0.60 as ‘likely benign’ with all others classified as ‘ambiguous’.

Gene-specific modes of disease inheritance were extracted from 450 virtual gene panels in Pan-elApp^67^ (https://panelapp.genomicsengland.co.uk/panels/518/, accessed 11 November 2024), restricted to clinical-grade ‘Green’ genes (confidence = 3). Missense variants were classified as ‘recessive’ if the gene was annotated as biallelic in every panel, and ‘dominant’ if annotated as monoallelic in every panel. Genes with multiple or conflicting annotations (both recessive and dominant possible) were excluded. Genes not present in any panel were labelled ‘none’.

Genomic constraint *z*-scores for chromosome 17, stratified into 1Kb regions, were obtained using the gnocchi method^55^ based on the gnomADv3 sequencing dataset (https://storage.googleapis.com/gcp-public-data--gnomad/release/3.1/secondary_analyses/genomic_constraint/constraint_z_genome_1kb.qc.download.txt.gz). Only scores within quality-controlled regions were used. Variants were assigned constraint classes between < − 4 (least constrained) and > 4 (most constrained) if they overlapped a scored region.

### 16 PCA group assignment

Individuals in the GEL dataset were assigned continental group labels according to their genetic distance in PC space to a set of nine reference groups released as part of gnomADv4.1.^55^ Following the gnomAD protocol, the `pc project` function from Hail^128^ was used to project sample genotypes and reference group allele frequencies onto the top 20 gnomADv4.1 PC axes (`gs://gcp-public-data--gnomad/release/4.0/pca/gnomad.v4.0.pca_loadings.ht`). The `assign population pcs` function from the gnomAD Hail utilities was then used to predict group assignments using the trained `sklearn ONNX` random forest model, with a minimum assignment probability threshold of 0.8. Individuals with a fitted probability of < 0.8 to all reference groups were labelled as “Remaining individuals”. We refer to the groupings of individuals as “PCA groups” throughout.

### 17 Individual vs all coalescence counts

To quantify how uneven ancestry representation in the GEL dataset influences the density and timing of genealogical coalescences we randomly subsampled up to 500 focal individuals per PCA group (or included all individuals when *n* < 500). For each focal individual, coalescence events were counted against all other individuals within the largest contiguous inferred ARG segment (17q4; Tab. S2). Coalescence counts were computed using `tskit`’s `pair_coalescence_counts` in evenly spaced, logarithmically scaled time windows. Results are shown in Fig. S20.

### 18 Cryptic relatives in the GEL dataset

Although parents from duo and trio families within the broader `aggV2` cohort were excluded from ARG inference, singleton variant counts remain highly sensitive to the presence of close biological relatives. To mitigate this effect, we used the genetically inferred kinship matrix previously generated by the Genomics England bioinformatics team (https://re-docs.genomicsengland.co.uk/principal_components/) to identify cryptic relatedness. From the total cohort of 47,535 individuals in the ARG, we excluded 4,849 with at least one genetically inferred < 3^rd^ degree relative from all analyses involving singleton variants.

### 19 Z-scoring allele ages within allele count bins

Allele age is correlated with allele frequency, with younger alleles generally occurring at lower frequencies (e.g., Fig. S17). To control for this relationship, we normalized allele ages by *z*-scoring within discrete derived allele count (DAC) bins. For each allele *i* with estimated age *t*_*i*_ and DAC bin *g*(*i*), we applied a log_10_ transformation followed by within-bin standardization:

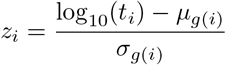

where *µ*_*g*(*i*)_ and *σ*_*g*(*i*)_ represent the mean and standard deviation of log_10_(*t*) within DAC bin *g*(*i*). This yields a *z*-score *z*_*i*_ that reflects how old or young an allele is relative to others of the same frequency, enabling frequency-controlled comparisons across annotations and individuals.

### 20 Quantile-quantile (QQ) analysis

To compare missense allele age distributions stratified by AlphaMissense or PrimateAI-3D pathogenicity class or PanelApp mode of inheritance, 100 samples from the gamma posterior of each variant were pooled to compute quantiles for each category of variant. For each of Figs. 4a.i and 4b.i, a “control” class is chosen, and then for each percentile, the ratio of that percentile in the comparison class to the value in the control class is shown, plotted against the percentile in the control class. In other words, for each value of 0 < *p* < 1, let *x*(*p*) and *y*(*p*) be the *p*-th quantiles of the “likely benign” and “likely pathogenic” age distributions, respectively; then each point on the red curve in Fig. 4a.i is of the form (*x*(*p*), *y*(*p*)*/x*(*p*)), for some *p*. For each non-control group, this procedure was repeated 100 times using bootstrap sampling with replacement to generate 95% confidence intervals in QQ plots.

### 21 Logistic regression analysis

To quantify allele enrichment within frequency-controlled allele age *z*-score bins (Methods §19) we used logistic regression to compare the frequency of variants assigned to each annotation class against all other variants with valid annotations. For AlphaMissense, PrimateAI-3D and PanelApp mode of inheritance classes, comparisons were restricted to missense variants in the GEL dataset across all contigs with assigned class labels. For constraint, comparisons were made using *all* variants within regions with an annotated constraint level on chromosome 17.

### 22 Clinically classified variants

Variants in the GEL ARG were annotated with clinical classifications based on the ACMG/AMP framework.^129^ Classifications were obtained from ClinVar^130^ (January 2024 release; https://ftp.ncbi.nlm.nih.gov/pub/clinvar/vcf_GRCh38/weekly/clinvar_20240107.vcf.gz) and Genomics England’s internal clinical scientist rare diseases outcomes questionnaire (main programme release v19; https://re-docs.genomicsengland.co.uk/exit_questionnaire/). Variants with discordant classifications, either within or across sources, were excluded prior to down-stream analyses. Pathogenic and likely pathogenic variants were grouped as P/LP, and benign and likely benign variants were grouped as B/LB. Clinically classified variants (DAC > 1) whose child nodes showed extreme uncertainties in ages were excluded.

### 23 ARG-informed heuristic to identify likely recurrent mutations

Recurrent mutations—independent mutational events at the same genomic site—violate a core assumption of the ARG inference method we used that each mutation originates once and is inherited along a single ancestral branch. Such violations can distort local genealogies and bias allele age estimates, and hence pose a particular challenge when analysing individual variants. To mitigate the effect of recurrent mutations biasing our analysis of clinically classified variants, we devised a heuritic identify mutations more likely to be recurrent by examining the local genealogical structure of each ultra-rare clinically classified variant with DAC > 1 and DAF < 0.1%.

For each variant, we computed the edge span (the genomic interval over which the ancestral branch carrying the mutation is inherited) and the local identity-by-descent (IBD) span (the region where all carriers share a most recent common ancestor at the node immediately below the mutation).

Under the assumption of a single origin, the mutation-bearing edge would typically fall within a broader region of shared ancestry. In contrast, recurrent or misinferred mutations are more likely to generate short, isolated branches or apparent discontinuities in the inferred ARG. Accordingly, we flagged mutations as more likely to be recurrent if (a) the mutation coincided with the boundary of its edge span, suggesting the introduction of an artificial recombination breakpoint during inference to accommodate inconsistent mutation patterns; or (b) the edge span was shorter than or equal to the local IBD span, indicating genealogical inconsistency with the surrounding ancestry.

Following this, we annotated each mutation as a CpG>TpG transition (or its reverse complement) using the ancestral reference sequence. We first identified CpG dinucleotides by scanning the ancestral FASTA sequence for consecutive “CG” bases and storing their genomic coordinates. Mutations were then labelled as CpG>TpG transitions if they occurred at one of these CpG dinucleotide positions and involved a C to T or G to A substitution.

We then evaluated our recurrence heuristic across DAC bins and by mutation type. As expected, mutations classified as CpG>TpG transitions—known hotspots for recurrent events due to methylation-mediated deamination—were more frequently flagged as likely recurrent than other mutation types (Fig. S24) We also observed an increasing proportion of flagged variants at higher DAC values, consistent with the expectation that variants with more derived alleles have a higher likelihood of including at least one recurrent event.

#### Acknowledgements

DR is supported by Wellcome (DPhil in Genomic Medicine and Statistics, 228319/Z/23/Z). JK, NSP and PLR acknowledge support from research grant R01 HG012473 from the National Institutes of Health NHGRI. JK acknowledges support from the Robertson Foundation.

We gratefully acknowledge the participants of the National Genomic Research Library (NGRL), whose contributions made this research possible. Secure access to the NGRL under project ID RR690 was provided by Genomics England, which delivers the NGRL in partnership with NHS England, and is wholly owned by the UK Department of Health and Social Care. The NGRL contains participants’ health data collected by the NHS as part of their care, along with samples and data from their participation in research, for which fully informed consent has been obtained. This includes genomic and clinical data provided through the NHS Genomic Medicine Service, as well as data obtained through research studies, including the 100,000 Genomes Project and the Generation Study, both of which are delivered in partnership with the NHS, and from other research cohorts involving external collaborators.

Data from the National Genomic Research Library (NGRL) used in this research are available within the secure Genomics England Research Environment. Access to NGRL data is restricted to adhere to consent requirements and protect participant privacy. Data used in this research include: phased genotypes from the Aggregated Variant Calls (AggV2) dataset provided by the University of Oxford (`/gel_data_resources/main_programme/aggregation/aggregate_gVCF_strelka/aggV2/phased_data/genotypes/`) and singleton genotypes extracted from the unphased AggV2 dataset (`/gel_data_resources/main_programme/aggregation/aggregate_gVCF_strelka/aggV2/genomic_data/`). Access to NGRL data is provided to approved researchers who are members of the Genomics England Research Network, subject to institutional access agreements and research project approval under participant-led governance. For more information on data access, visit: https://www.genomicsengland.co.uk/research

Computation used the Oxford Biomedical Research Computing (BMRC) facility, a joint development between the Wellcome Centre for Human Genetics and the Big Data Institute supported by Health Data Research UK and the NIHR Oxford Biomedical Research Centre. The views expressed are those of the author(s) and not necessarily those of the NHS, the NIHR or the Department of Health.

## Supplementary Information

This section contains the supplementary information, which we integrate into the main document for ease of cross referencing during review.

### 1 Expectation propagation update

Here we derive the moments used in the expectation propagation update described in Methods §1. Recall that the “surrogate” distribution for which we wish to compute moments has the unnormalized form,

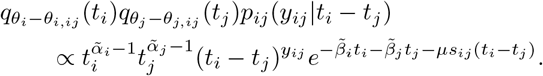

Then, if we define

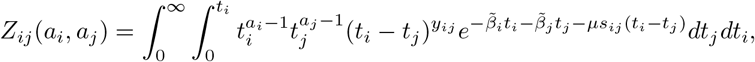

then we can obtain the moments through

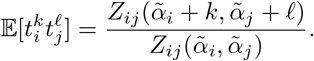

To evaluate this, rewrite the integral with *T* = *t*_*i*_ and *uT* = *t*_*j*_, so that

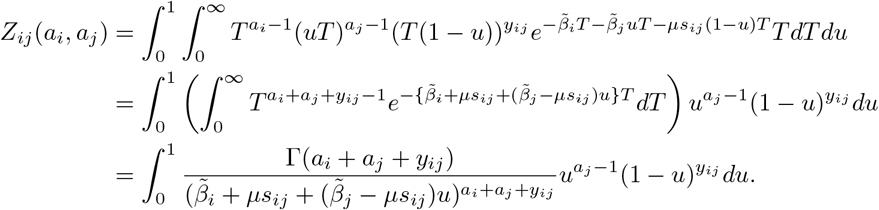

Next we use the fact that (ref^131^ §3.197.3)

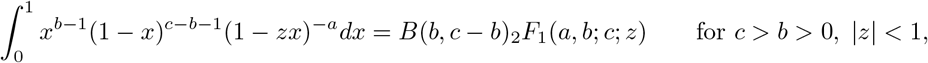

where *B* is the beta function and _2_*F*_1_ is the Gaussian hypergeometric function. With *b* = *a*_*j*_, *c* = 1 + *a*_*j*_ + *y*_*ij*_, *a* = *a*_*i*_ + *a*_*j*_ + *y*_*ij*_, and 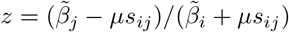, this is

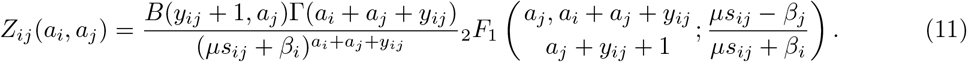

If node *j* is a contemporary sample (*t*_*j*_ = 0) this simplifies to

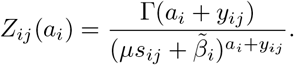

Defining

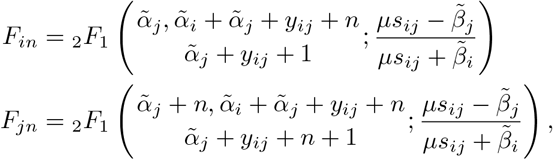

we can thus write the first two moments as

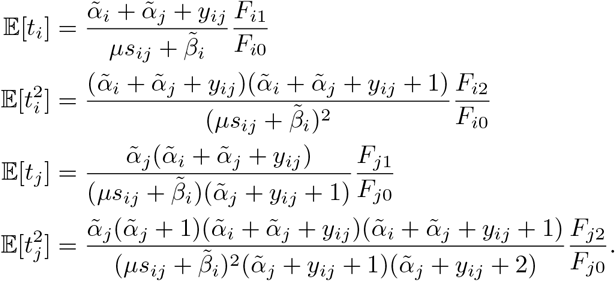

Or, when node *j* is a contemporary sample (*t*_*j*_ = 0),

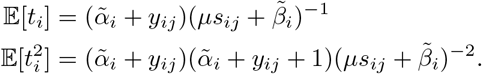

The updated natural parameters are obtained from the moments by

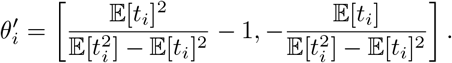

As a final note: the computational bottleneck in the fitting procedure will be computing values of _2_*F*_1_, so it is helpful to reduce the number of distinct values we need to compute. Using Gauss’s contiguous relation that *z*(*ab/c*)_2_*F*_1_(*a* + 1, *b* + 1; *c* + 1; *z*) = *b*(_2_*F*_1_(*a, b* + 1; *c* + 1; *z*) − _2_*F*_1_(*a, b*; *c*; *z*)), we can write *F*_*i*1_*/F*_*i*0_ in terms of *F*_*j*1_*/F*_*j*0_, and similarly reduce *F*_*i*2_*/F*_*i*0_, obtaining that

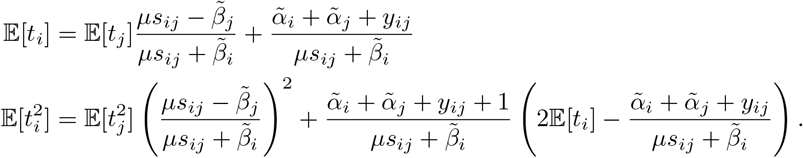

**2 Numerically stable approximations of** _2_*F*_1_

The Gaussian hypergeometric function _2_*F*_1_ can be difficult to compute in a numerically stable fashion for arbitrary parameter regimes without relying on multiprecision floating point arithmetic, due to catastrophic cancellation between alternating terms in the defining hypergeometric series.

A numerically stable alternative is to approximate _2_*F*_1_ via Laplace’s method, using the scheme in^98^ (derived here for completeness). The idea is to use Euler’s integral identity

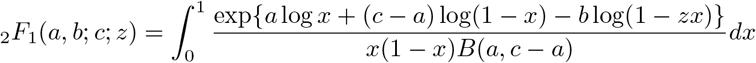

and change variables such that the transformed integrand is well-approximated by a Gaussian function. Note that *f* (*x*) = − *a* log *x* − (*c* − *a*) log(1 − *x*) + *b* log(1 − *zx*) has a minimum in (0, 1) given by

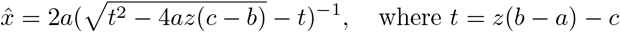

provided *c* > *a* > 0 and *z* > 0. The first condition will always be satisfied in our application; and if *z* < 0 then the Pfaff transform _2_*F*_1_(*a, b*; *c*; *z*) = (1 − *z*)^−*b*^_2_*F*_1_(*c* − *a, b*; *c*; *z*(*z* − 1)^−1^) is used instead.

Making the logit change of variables by setting *x*^′^ = *g*(*x*) = log(*x/*(1 − *x*)) (and so *dx*^′^ = *x*^−1^(1 − *x*)^−1^*dx* and *x*^′^ = *g*(*x*) = log *x* − log(1 − *x*), 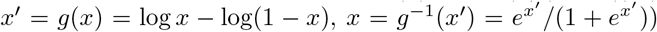 and taking a second-order Taylor expansion of *f* (*g*^−1^(*x*^′^)) around 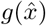 leads to:

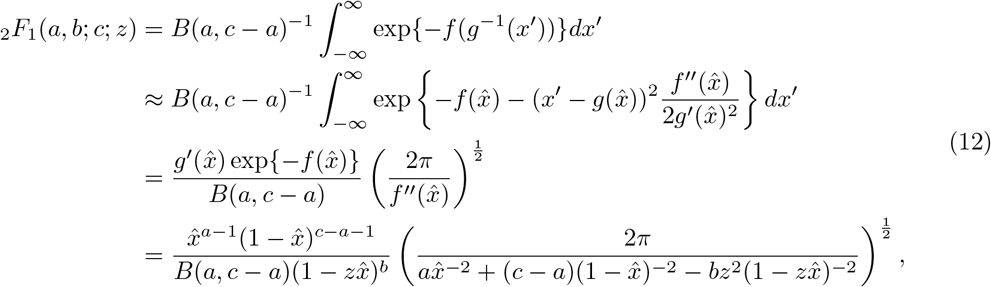

where the first-order term has disappeared from the Taylor approximation on the second line because 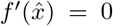. Finally,^98^ suggest using the identity _2_*F*_1_(*a, b*; *c*; 0) = 1 to calibrate the approximation, via

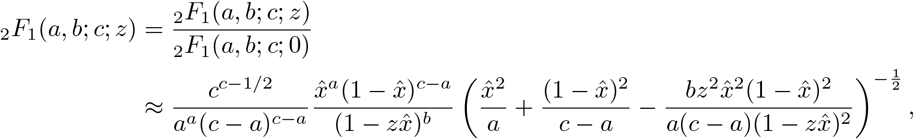

which follows from applying (12) separately to the numerator and denominator and simplifying the result.

### 3 Non-contemporary samples and internal constraints

In previous sections we have assumed that sample nodes are contemporary (fixed to time zero), in which case the expectation propagation surrogate for singleton edges reduces to a Gamma distribution and moments follow directly. However, if an edge *ij* has one end fixed to a particular nonzero time, the solution is more complicated. As above we used the hypergeometric function _2_*F*_1_, here we will encounter the hypergeometric functions of Tricomi and Kummer, *U* and *M*, with integral representations

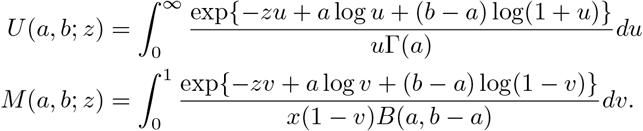

Following the notation of SI §1 for the edge *ij*, define for later use

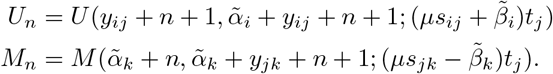

First suppose the child *j* has fixed time *t*_*j*_. Then the normalizer analogous to *Z*_*ij*_ in (11) is

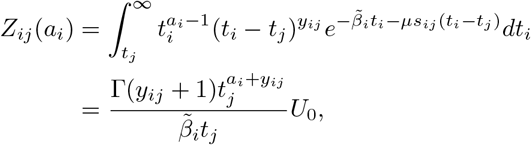

and so

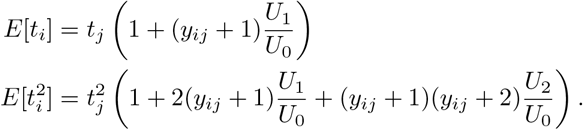

Now suppose that the edge is *jk* with fixed time *t*_*j*_ (where *j* is now the parent). In this case, the normalizer is

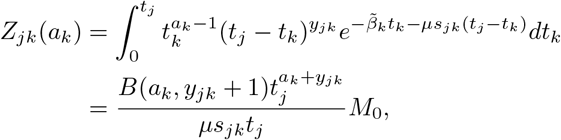

and so

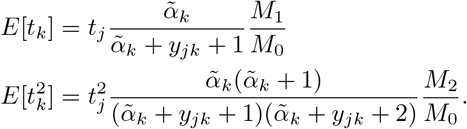

Calculating these moments in practice requires evaluation of the Tricomi hypergeometric function *U* (*a, b*; *z*) or the Kummer hypergeometric function *M* (*a, b*; *z*). As for _2_*F*_1_, direct evaluation of the defining hypergeometric series can suffer from numerical instability under certain parameter regimes. Given the integral identities,

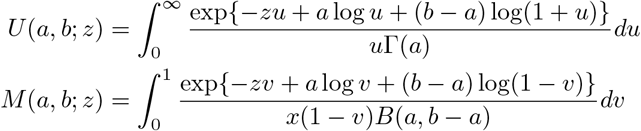

changing variables to *du*^′^ = *u*^−1^*du* and *dv*^′^ = *v*^−1^(1 − *v*)^−1^*dv*, then using Laplace’s method leads to the numerically stable approximations,

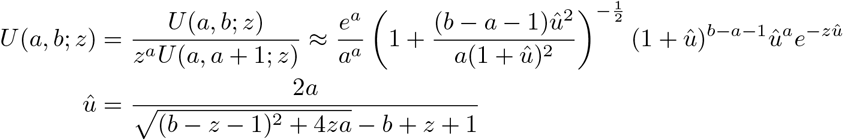

and

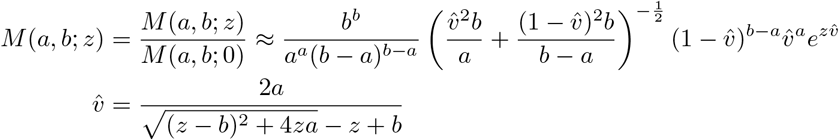

where we have used the identities *z*^*a*^*U* (*a, a* + 1; *z*) = 1 and *M* (*a, b*; 0) = 1 to calibrate the approximations, analogously to SI §2.

### 4 Mutation ages

Mutations are mapped to edges in the ARG, but we cannot know more precisely where a particular mutation is located in time. Implicit in the Poisson mutation process from Methods §1 is the assumption that mutations are uniformly distributed along the time dimension of an edge. Thus the *n*th conditional moment of the age of a mutation on edge *ij* (with parent *i*, child *j*) is 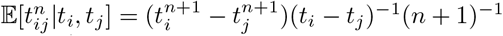. Let 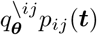 be the variational surrogate for edge *ij* (i.e. the “cavity” distribution for edge *ij* multiplied by its Poisson likelihood), defined in equation (3). As described previously, the surrogate distribution is globally approximate (in that it replaces the joint distribution of node ages by a factorized variational approximation) but locally exact (in that it models the dependence between the age of parent *i* and child *j* that is due to the likelihood of edge *ij*). Under this approximation, the age *t*_*ij*_ of a mutation on the edge has the *n*th unconditional moment,

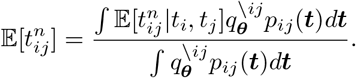

Using the precise definition of the variational gamma surrogate and following the rationale and notation of SI §1, let

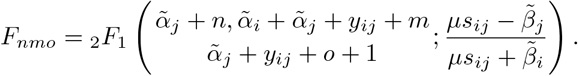

Then the first two moments of the mutation’s age are,

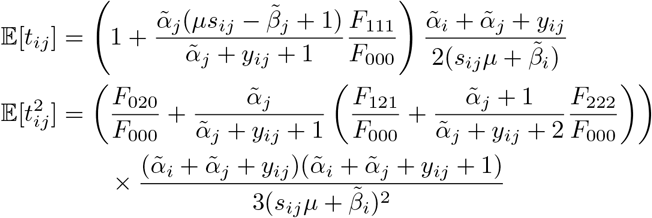

and may be used to construct a variational gamma approximation, as was done for node ages.

### 5 Ambiguous singleton phasing

In diploid data, mutations with frequency one are likely to have ambiguous phase. In this case, the model in SI §1 may be adjusted so as to integrate over both possible phases for each singleton mutation. This requires splitting the edges above each leaf so that any given individual has at most two immediate ancestors (nodes) over the span of every attached edge. For example, imagine we have a sequence of length *L* containing a diploid individual where the first haplotype has immediate ancestor *i* over interval [0, *a*) and ancestor *j* over interval [*a, L*); and similarly the second haplotype has ancestor *k* over interval [0, *b*) and ancestor *l* over interval [*b, L*). Each of these intervals is represented by an edge in the tree sequence data structure, and in general *a* ≠ *b*; that is, edges can overlap between the two haplotypes. After splitting leaf edges as described above for this example, if *a* < *b*, the individual is subtended by a pair of edges leading to ancestors *i* and *k* over the interval [0, *a*), a pair leading to *j* and *k* over [*a, b*), and a pair leading to *j* and *l* over [*b, L*).

Consider an individual whose haplotypes are *u* and *v*, and let ℰ_*uv*_ be the edges that end in *u* or *v* (i.e., ℰ_*uv*_ = {*ij* ∈ ℰ: *j* = *u* or *j* = *v*}). After splitting as described above, these edges come in concurrent pairs of the form (*iu, jv*), where *i* and *j* are the immediate ancestors of the individual on a given genomic segment (and possibly *i* = *j*). Each pair is associated with an interval of span *s*_*iu,jv*_, and these intervals are nonoverlapping. If a singleton mutation over this interval is unphased, then it cannot be assigned to either edge *iu* or edge *jv*. If the individual (*u, v*) is contemporary (*t*_*u*_ = *t*_*v*_ = 0) then the total branch area of the edge pair is *s*_*iu,jv*_(*t*_*i*_ + *t*_*j*_). Assuming all singleton mutations on the edge pair are unphased, the likelihood for the total count *y*_*iu,jv*_ of such mutations is

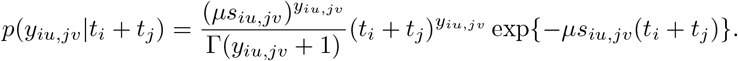

Let ℐ denote the set of individuals, ℰ_ℐ_ = ∪_*uv*∈ℐ_ ℰ_*uv*_ the set of all edges subtended by individuals, and 𝒫_*uv*_ the set of concurrent edge pairs for individual *uv*. Then the joint distribution of node ages becomes

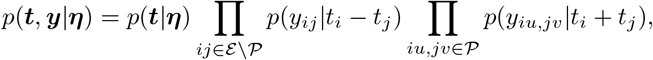

where 𝒫 are all split leaf edges and ℰ \ 𝒫 are all internal edges (e.g., edges carrying phased, non-singleton mutations).

Following the rationale in SI §1 leads to an expectation propagation update for the ages of the two ancestors *t*_*i*_ and *t*_*j*_ involved in a given singleton edge pair:

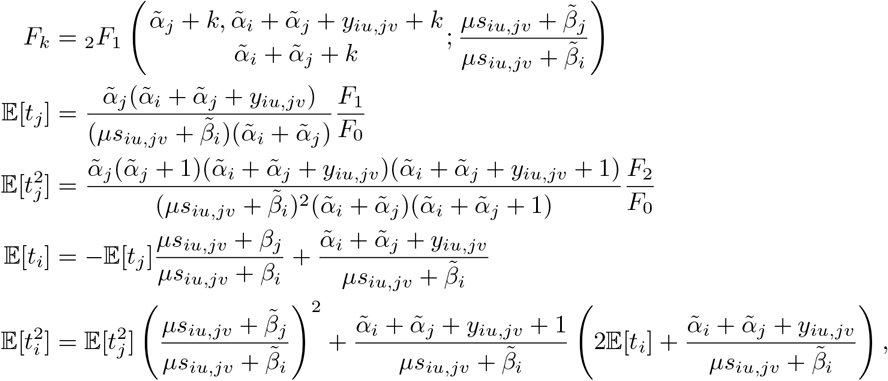

assuming these ancestors are distinct (*i* ≠ *j*). If instead a single ancestor is attached to both edges of the pair (so *i* = *j*), the update becomes

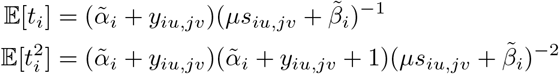

because the surrogate reduces to a Gamma distribution.

## 6 Mixture prior on root ages

In SI §1, we assumed generic gamma priors for particular nodes without specifying how these are determined. Information about the ages of nodes comes from noisy measurements of edge lengths – mutational density – that are propagated up the ARG from a fixed reference point (the sample ages). Typically, the oldest ancestral segments will each have very short spans, because of the cumulative action of recombination. Mutational density is sparse on these ancient haplotypes, and there is relatively little ancestral material that can act as constraint on their ages. As a consequence, the posteriors of the oldest nodes typically have extremely high variance, and this is exacerbated in the variational approximations by the local nature of expectation propagation. Thus, we regularise the ages of “ultimate” roots (the nodes with no parents) via a prior, and use flat priors for all other nodes. This scheme acts as a soft constraint on the maximum height of the ARG. Because the choice of prior introduces another decision for the user with potentially large consequences on the quality of inference, we fit an arbitrarily flexible prior via an Empirical Bayes method.

To this end, we employ a mixture of gamma distributions for a prior on the ages of ultimate roots, with hyperparameters that are estimated during each EP iteration from the current variational approximation. Explicitly, let the hyperparameters *η*_*k*_ = {*ω*_*k*_, *γ*_*k*_, *κ*_*k*_} be the mixture weight, shape, and rate for the *k*th mixture component. The prior for the *i*th node is

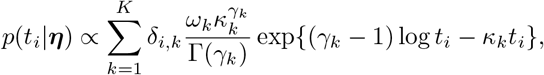

where the node-specific weights *δ*_*i,k*_ are assumed to be specified *a priori*, so that the choice *δ*_*i,k*_ = 1 if node *i* has no parents and *δ*_*i,k*_ = 0 otherwise leads to an i.i.d. prior across ultimate roots. Other choices of *δ* allow stratification of the prior across frequency classes or windows along the genome, or extension of the prior to non-root nodes. As before, the messages due to the prior are removed from the variational approximation to create the cavity 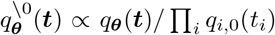 with natural parameters 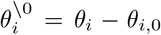 and then updated by matching moments against the surrogate *q*^\0^(***t***) Π_*i*_ *p*(*t*_*i*_ |***η***).

We extend this standard EP update with an expectation-maximization (EM) procedure that finds optimal hyperparameters for the surrogate. To perform EM, we augment the model with per-node binary variables *z*_*i,k*_ that indicate if node *i* belongs to mixture component *k*, so that under this model ∑_*k*_ *z*_*i,k*_ = 1 and ∑*z*_*i,k*_ = 1 with probability *ω*_*k*_, and then if *z*_*i,k*_ = 1 then *t*_*i*_ is drawn from the distribution with parameters (*γ*_*k*_, *κ*_*k*_). The joint posterior of ***t*** and ***z*** is then

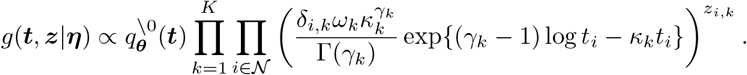

Now our goal is, given a current set of parameters *η*^(*n*)^, to find *η*^(*n*+1)^ to maximize the expected value of log *g*, averaging ***t*** and ***z*** across the posterior. To do this, let *g*_*n*_ ≡ *g*(***t, z***|***η***^(*n*)^) denote the augmented surrogate at the *n*th EM iteration. The expectation step yields the objective function,

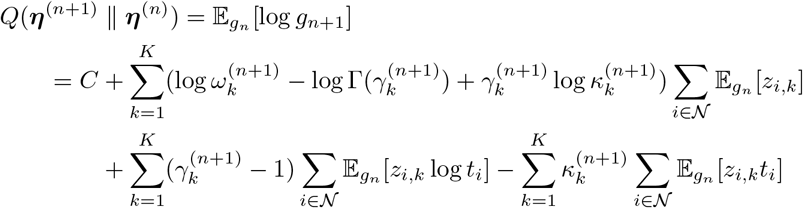

where the necessary expectations involve standard manipulations of gamma distributions,

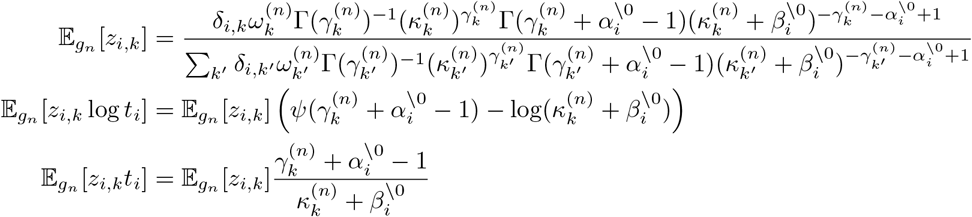

and we’ve used the canonical parameterization 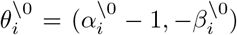. The maximization step ***η***^(*n*+1)^ = arg max_***η***_*′ Q*(***η***^′^ ∥ ***η***^(*n*)^) gives the updates

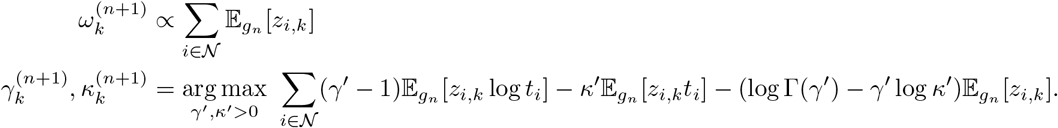

Note that the second line is equivalent to finding maximum likelihood estimates of gamma parameters from sufficient statistics Upon convergence, integrating over the optimized surrogate yields the approximate posterior moments for the ages of individual nodes, 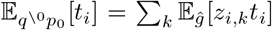 and 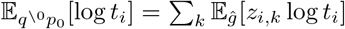, that are used to calculate updated variational parameters 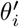 and prior messages 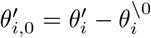.

## 7 Timescale calibration algorithm

Here we describe the time-rescaling algorithm used by tsdate. First we describe a simple approach that matches mutation counts and edge area over time, and then an extension that is robust to spurious polytomies in the genealogies.

Under the mutational clock, the expected total number of mutations that fall on an ARG in some time window [*a, b*) is equal to the mutation rate *µ* multiplied by the total area of all edges contained in that window. Suppose an edge *e* between parent *p*_*e*_ and child *c*_*e*_ has span *s*_*e*_ and *y*_*e*_ mutations; the area of this edge that intersects [*a, b*) is 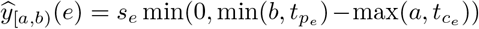. The corresponding observed number of mutations on edge *e* is obtained by integrating over mutations’ temporal positions on their edge: 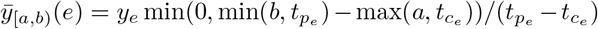, The total expected and observed counts are obtained by summing over all edges: *ŷ* _[*a,b*)_ = ∑_*e*∈E_ *y* _[*a,b*)_(*e*) and 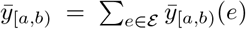. Then, the interval length is rescaled so that expected and observed mutational counts match, mapping *a* and *b* to *a*^′^ and *b*^′^ where 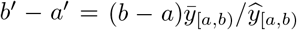. The endpoints of all intervals are adjusted given the new lengths and the constraint that the first interval starts at zero, so that the age of a given node *k* located in [*a, b*) is updated using the rescaled length and lower bound, to 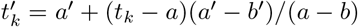.

There are two problems with applying this core idea. First, tsinfer encodes topological ambiguity in the ARG by polytomies. That is, a given node may have multiple children in a given inferred marginal tree when underlying true genealogy is binary. A consequence of these artefactual polytomies is, roughly, that the mutational area is overcounted, and rescaling based on observed counts of mutations will bias dates downwards. While the impact of polytomies on total branch area is minimal when the ARG contains relatively few samples, the bias can become extreme for large datasets when polytomies may contain hundreds or thousands of nodes.

To see the issue, consider a single tree in which a branch from parent *i* to child *j* has no mutations. In the inferred tree, *i* and *j* will be merged; but what should the “inferred” time of the resulting node be? There is no *a priori* best answer, but an argument can be made that the parent time *t*_*i*_ is a better choice: suppose that samples *a* and *b* have node *j* as their MRCA, and sample *c* has node *i* as its MRCA with *a* and *b*. Then if we were to place the merged node “*ij*” more recently than *t*_*i*_, this would imply that *a, b*, and *c* share a common ancestor more recently than they actually do. Furthermore, the time *t*_*ij*_ of the merged node provides the upper bound on the time of any events occurring on the edge between *i* and *j*; so if *t*_*ij*_ is less than the true time *t*_*i*_, we will incorrectly constrain the times of those events. (These “events” might be a stand-in for how information in this tree affects dating of nodes and mutations in adjacent trees.) On the other hand, setting the merged node’s time *t*_*ij*_ to *t*_*i*_ results in an overestimate of the TMRCA of *a* and *b* (unless perhaps we interpret it as the “most recent inferrable ancestor”), but provides an accurate upper bound (given available information) on the times of those events.

The solution is to choose a statistic for the rescaling numerator and denominator that is not impacted by artefactual polytomies. Consider sampling a path from a randomly selected sample to the root in a given marginal tree. The number of mutations and length of the path is unaffected by any missing intermediate nodes (e.g., nodes which have been collapsed into their parent). Taking the average over all possible paths and summing across marginal trees leads to similar statistics as *ŷ* and 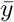, but weighted by the number of descendant samples. For a tree 𝕋 let *s* _𝕋_ denote the span of the tree, *d* _𝕋_ (*e*) the number of samples descending from edge *e* in tree 𝕋, and *y* _𝕋_ (*e*) the number of mutations on edge *e* in the tree. Then, define

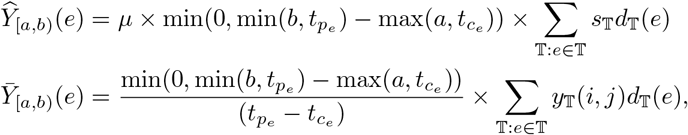

where the summations are over marginal trees that contain the edge *e*. As above, define *ŷ* _[*a,b*)_ and 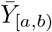 as the sum of these over all edges, which may be computed efficiently in 𝒪 (*N* +*T* log *N*) time using the incremental strategy of,^47^ where *N* is the number of samples in the ARG and *T* is the number of marginal trees. Subsequently, node ages are rescaled as described above for segregating sites but with *Y*_[*a,b*)_*/Y*_[*a,b*)_ as the rescaling factor.

The second challenge we need to solve is that rather than rescaling point estimates of node age we need to rescale variational posteriors, which implies integrating a distribution over a piecewise constant function to calculate sufficient statistics for moment matching. This is costly for large ARGs.

A solution is to rescale posteriors by matching quantiles rather than moments. We will choose new parameters for each node *i* to match two things: the median and the ratio between the third and first quantiles. Denote the desired piecewise linear map to the transformed timescale as *t*^′^ = *F* (*t*). The value *F* (*t*) can be computed in 𝒪 (log *K*) time given precomputed rescaling factors for *K* intervals, by using a binary search to find the interval bracketing *t*. Let *Q*^−1^(*p*|*α, β*) (for *p* ∈ [0, 1]) denote the inverse CDF for the Gamma distribution with shape *α* and scale *β*. Recall that the Gamma distribution is a scale family with the property *Q*^−1^(*p*|*α, β*) = *β*^−1^*Q*^−1^(*p*|*α*, 1). Fix two quantiles *p*_1_ < *p*_2_ and define the function

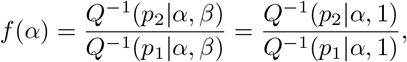

which is monotonically decreasing in *α* > 0, and therefore has an inverse that may be found numerically with a small number of iterations of Newton’s method. Now suppose the variational posterior of node *i* is Gamma(*α*_*i*_, *β*_*i*_); we wish to find 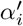 and 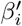 such that 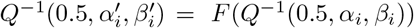 and (with the arbitrary choice *p*_1_ = 0.25 and *p*_2_ = 0.75),

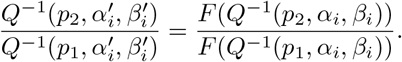

To do this, we first solve 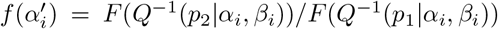 for 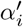, and then set 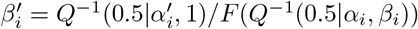.

## 8 Genomic properties of regions used for inference

We compared the genic content and recombination rate of regions selected for ARG inference of the GEL dataset (Methods §14) to those of randomly sampled autosomal regions of identical sizes, and used genome-wide allele age estimates to assess whether these genomic properties were associated with differences in allele age. All analyses were restricted to autosomes (chr1–22).

Inference regions were defined as the fixed set of autosomal segments used for ARG construction (listed in Tab S2). For all comparisons, region lengths were preserved exactly, and summary statistics were aggregated across all inference regions.

Protein-coding gene annotations were obtained from Ensembl (GRCh38; https://ftp.ensembl.org/pub/release-115/gtf/homo_sapiens). Gene intervals annotated as protein-coding were restricted to autosomes and reduced to a non-overlapping union. For each region set, the proportion of bases overlapping protein-coding genes was calculated as the total number of genic bases divided by the total genomic span of the regions. For inference regions, genic content was calculated per region and aggregated across all regions.

Recombination rates were estimated using fine-scale genetic maps from HapMap II lifted to GRCh38 coordinates. For each region, genetic map positions at the start and end coordinates were obtained by linear interpolation, and genetic distance was calculated as the difference between these values. Recombination rate was expressed as centimorgans per megabase (cM/Mb). Aggregate recombination rates were computed by dividing the total genetic distance by the total physical length across all regions.

Mean allele ages were obtained from genome-wide Relate^37^ age estimates inferred using the 1,000 Genomes Project Yoruba (YRI) population (https://zenodo.org/records/3234689). For each region, Relate age estimates overlapping that interval were averaged, and an overall mean allele age was computed as a length-weighted mean across all regions.

To generate null distributions, 100 random region sets were sampled sampled from the callable autosomal genome, defined as autosomes excluding centromeric regions and annotated assembly gaps. Centromeres were identified using UCSC cytoband annotations (acen and gvar bands; https://hgdownload.soe.ucsc.edu/goldenPath/hg38/database/cytoBand.txt.gz), and assembly gaps were excluded using UCSC gap annotations for GRCh38 (https://hgdownload.soe.ucsc.edu/goldenPath/hg38/database/gap.txt.gz). Each replicate consisted of one region per inference segment, matched exactly by length. Regions were sampled uniformly across all valid genomic placements within the callable genome. To avoid positional bias, callable intervals were weighted by the number of valid start positions they contained for a given region length. For each replicate, genic content, recombination rate, and mean allele ages were recomputed using the same procedures applied to the observed inference regions.

The inference regions showed clear deviations from random expectations in terms of genic content and recombination rate (Supp Fig S13.a,b) — consistent with broad-scale chromosomal variation in gene density and recombination rate, given that the inference regions are concentrated on chromosomes 17–22, which include some of the most gene-dense human chromosomes (notably chromosomes 17^132^ and 19^133^) and are characterised by elevated recombination rates per unit physical length owing to the inverse relationship between chromosome size and recombination rate.^134^ In contrast, the mean allele age estimated from Relate YRI data for the inference regions lay well within the distribution obtained from random, length-matched regions (Supp Fig S13.c), with no strong deviation from the null expectation. This indicates that, despite being enriched for genic sequence and higher recombination the inference regions are not detectably biased with respect to ARG-derived genome-wide allele age estimates.

## 9 Age vs frequency as a predictor of selection

Using data simulated from a model of human evolution under purifying selection (Methods §12), we assessed the ability of mutation age versus allele frequency to predict the known selection coefficients, *s*, associated with each mutation. We considered simulations under balanced versus unbalanced sampling, which generated *n* = 1,419,535 vs *n* = 1,000,412 non-recurrent mutations respectively. We also repeated the analysis restricting these two datasets to ultra-rare variants only (DAF < 0.001; *n* = 1,110,527 vs *n* = 761,867).

Treating the selection coefficient *s* as a response variable, we fitted an additive linear model in R^135^ using the logarithm of age plus the logarithm of frequency as predictors, and an additional model which also included the mutation’s population of origin as a separate additive factor. The simulated DFE generates a highly skewed distribution of selection coefficients, with a majority of mutations being neutral (*s* = 0), resulting in residuals which do not conform to a simple statistical distribution. We therefore did not calculate p-values, but simply measure the variance contribution (pseudo *R*^2^) of the age versus frequency term by dropping each independently from the full model. Pseudo *R*^2^ values were calculated as the proportional reduction in deviance, 1 − *D*_reduced_*/D*_full_, where *D*_reduced_ is the deviance of the model with the dropped term. We report the *R*^2^ contributions of the age and frequency terms, and also calculate the ratio of the two values, to produce a measure of the relative predictive ability of age versus frequency in each model. Finally, we fitted models in which mutation age was taken either as the true (*t*_true_) or the tsinfer+tsdate inferred (*t*_tsdate_) midpoint time.

The results in Tab. S8 show that under a balanced sampling scheme, age and frequency are roughly equivalent in predictive power, to within a factor of 2. If true (midpoint) age is known, then over the whole dataset, age provides about twice as much information as frequency when predicting the selection coefficient, even if population-of-origin is taken into account. However, for ultra-rare variants, the relative power is reduced, and using inferred age results in frequency having slightly more predictive power. Nevertheless, over all the results, even the lowest *R*^2^ for age is 6.539e-6, which is comparable in magnitude to the highest *R*^2^ for frequency (1.684e-5).

Under the unbalanced sampling regime, allele age consistently explains orders of magnitude more variation in the selection coefficient than allele frequency. This is true both using inferred age or the true age as a predictor: in the latter case, age accounts for the entire prediction: the additional contribution of frequency is negligible. Accounting for population-of-origin somewhat reduces this difference, nevertheless in our models age always remains a better predictor of (negative) selection when sampling is unbalanced, even for ultra-rare alleles. We also note that the higher predictive power of age over frequency is maintained when changing the model to use the raw (unlogged) frequency values as a predictor (data not shown).

This simulation and analysis illustrates the potential predictive power of allele age; further work is required, however, to understand more fully these properties over a range of evolutionary scenarios. In practice, of course, both frequency (in the sample) and (estimated) age are available.

## Supplementary Tables and Figures

In this section we include all supplementary tables and figures.

**Table S1:** Accuracy of the new Variational Gamma (VG) algorithm compared to the original Inside-Outside (IO) algorithm in tsdate. Accuracy was assessed by comparing true (simulated) and inferred (`tsinfer` and `tsdate`) ARGs across four human `stdpopsim` models, over a 40 Mb section of chromosome 17 with 15000 diploids sampled evenly across populations. Root mean square error (RMSE) and bias are calculated for log_10_ mutation ages (estimated as edge midpoints), for all mutations and for low-frequency mutations segregating in < 0.1% of sample haplotypes (shown in parentheses). Statistics are shown separated by frequency in Fig. S1.

| stdpopsim model | VG |  | IO |  |
| --- | --- | --- | --- | --- |
|  | Bias | RMSE | Bias | RMSE |
| AmericanAdmixture_4B18 | −0.01 (−0.01) | 0.27 (0.27) | −0.32 (−0.38) | 0.48 (0.53) |
| OutOfAfricaArchaicAdmixture_5R19 | −0.03 (−0.04) | 0.32 (0.33) | −0.38 (−0.46) | 0.58 (0.65) |
| OutOfAfrica_3G09 | −0.01 (−0.01) | 0.27 (0.27) | −0.30 (−0.36) | 0.47 (0.51) |
| Zigzag_1S14 | −0.00 (0.00) | 0.31 (0.31) | −0.16 (−0.21) | 0.40 (0.44) |

**Table S2:** Inference regions. Details of the 15 regions used for inference, including coordinates, genic proportion, and average recombination rate (cM/Mb).

| region | start | end | length | gene content | recomb rate |
| --- | --- | --- | --- | --- | --- |
| 16p1 | 16,689 | 14,637,672 | 14,620,983 | 0.712 | 2.236 |
| 16p2 | 22,661,538 | 28,286,126 | 5,624,588 | 0.531 | 2.237 |
| 17p1 | 281,784 | 18,276,569 | 17,994,785 | 0.661 | 2.387 |
| 17q1 | 26,718,757 | 36,149,680 | 9,430,923 | 0.670 | 1.142 |
| 17q2 | 36,458,468 | 37,824,094 | 1,365,626 | 0.633 | 2.059 |
| 17q3 | 38,221,639 | 45,432,621 | 7,210,982 | 0.643 | 0.833 |
| 17q4 | 45,586,151 | 83,087,382 | 37,501,231 | 0.594 | 1.636 |
| 18p | 148,918 | 15,357,916 | 15,208,998 | 0.479 | 2.626 |
| 19p | 75,006 | 24,174,775 | 24,099,769 | 0.714 | 2.044 |
| 20p | 227,205 | 26,274,525 | 26,047,320 | 0.487 | 1.951 |
| 20q1 | 31,254,980 | 64,237,737 | 32,982,757 | 0.524 | 1.733 |
| 21q1 | 34,495,543 | 42,960,010 | 8,464,467 | 0.726 | 2.275 |
| 21q2 | 44,253,397 | 46,680,226 | 2,426,829 | 0.702 | 1.713 |
| 22q1 | 16,439,646 | 18,110,684 | 1,671,038 | 0.560 | 3.230 |
| 22q2 | 18,897,685 | 21,045,540 | 2,147,855 | 0.715 | 2.980 |
| Total | N/A | N/A | 206,798,151 | 0.600 | 1.942 |

**Table S3:** CPU resources for ARG inference. For each region listed in Tab. S2 we show the CPU time consumed by different phases of inference. Note that this is the cumulative user time across all threads and processes and not the elapsed wall-clock time.

| region | gen_ancestors | match_ancestors | match_samples | tsdate | total |
| --- | --- | --- | --- | --- | --- |
| 16p1 | 8.28h | 6.37d | 1.54y | 14.03m | 1.56y |
| 16p2 | 1.16h | 2.65d | 88.04d | 6.37m | 90.74d |
| 17p1 | 17.96h | 14.13h | 2.62y | 19.45m | 2.63y |
| 17q1 | 2.23h | 5.29d | 176.03d | 7.58m | 181.42d |
| 17q2 | 13.89m | 3.44h | 15.54d | 2.69m | 15.70d |
| 17q3 | 1.54h | 3.92d | 112.25d | 5.79m | 116.24d |
| 17q4 | 1.06d | 6.23d | 1.89y | 41.56m | 1.91y |
| 18p | 5.92h | 9.51d | 1.49y | 16.19m | 1.52y |
| 19p | 12.92h | 6.63d | 2.15y | 14.85m | 2.17y |
| 20p | 1.29d | 11.72h | 3.29y | 22.99m | 3.29y |
| 20q1 | 1.12d | 1.89d | 1.34y | 39.86m | 1.35y |
| 21q1 | 2.65h | 5.72d | 196.69d | 9.85m | 202.53d |
| 21q2 | 30.34m | 18.85h | 36.83d | 1.42m | 37.64d |
| 22q1 | 8.26m | 5.71h | 14.66d | 3.18m | 14.90d |
| 22q2 | 13.16m | 6.70h | 36.47d | 3.53m | 36.76d |
| Total | 5.71d | 50.74d | 16.18y | 3.49h | 16.33y |

**Table S4:** Tree sequence statistics by region in the GEL dataset. For each region listed in Tab. S2 we show the number of variant sites used in inference (sites); the number of trees, nodes, edges, and mutations in the final inferred tree sequence; and the size of tszip file.

| region | length (bp) | sites | trees | nodes | edges | muts | file size |
| --- | --- | --- | --- | --- | --- | --- | --- |
| 16p1 | 14,620,983 | 2,207,614 | 708,992 | 1,793,817 | 14,582,277 | 2,207,614 | 247.3MB |
| 16p2 | 5,624,588 | 648,035 | 202,069 | 531,263 | 3,544,965 | 648,035 | 64.5MB |
| 17p1 | 17,994,785 | 2,054,172 | 706,131 | 1,683,050 | 13,588,316 | 2,054,172 | 244.5MB |
| 17q1 | 9,430,923 | 963,103 | 303,130 | 738,625 | 4,403,891 | 963,103 | 88.3MB |
| 17q2 | 1,365,626 | 148,058 | 41,648 | 195,915 | 925,266 | 148,058 | 14.2MB |
| 17q3 | 7,210,982 | 767,500 | 234,079 | 589,059 | 3,379,624 | 767,500 | 65.6MB |
| 17q4 | 37,501,231 | 4,054,901 | 1,252,598 | 2,949,271 | 22,785,855 | 4,054,901 | 432.0MB |
| 18p | 15,208,998 | 1,600,531 | 490,490 | 1,271,875 | 11,525,869 | 1,600,531 | 189.4MB |
| 19p | 24,099,769 | 2,396,917 | 797,814 | 1,981,628 | 17,134,593 | 2,396,917 | 284.5MB |
| 20p | 26,047,320 | 2,934,752 | 858,060 | 2,093,350 | 16,711,709 | 2,934,752 | 304.1MB |
| 20q1 | 32,982,757 | 3,783,588 | 1,153,190 | 2,752,320 | 21,697,573 | 3,783,588 | 403.6MB |
| 21q1 | 8,464,467 | 988,743 | 326,829 | 827,212 | 6,195,030 | 988,743 | 110.7MB |
| 21q2 | 2,426,829 | 304,942 | 109,770 | 342,957 | 2,007,819 | 304,942 | 34.0MB |
| 22q1 | 1,671,038 | 147,455 | 53,852 | 220,890 | 1,352,677 | 147,455 | 19.5MB |
| 22q2 | 2,147,855 | 223,988 | 75,972 | 261,365 | 1,448,721 | 223,988 | 23.4MB |
| Total | 206,798,151 | 23,224,299 | 7,314,624 | 16,806,547 | 141,284,185 | 23,224,299 | 2.5GB |

**Table S5:**
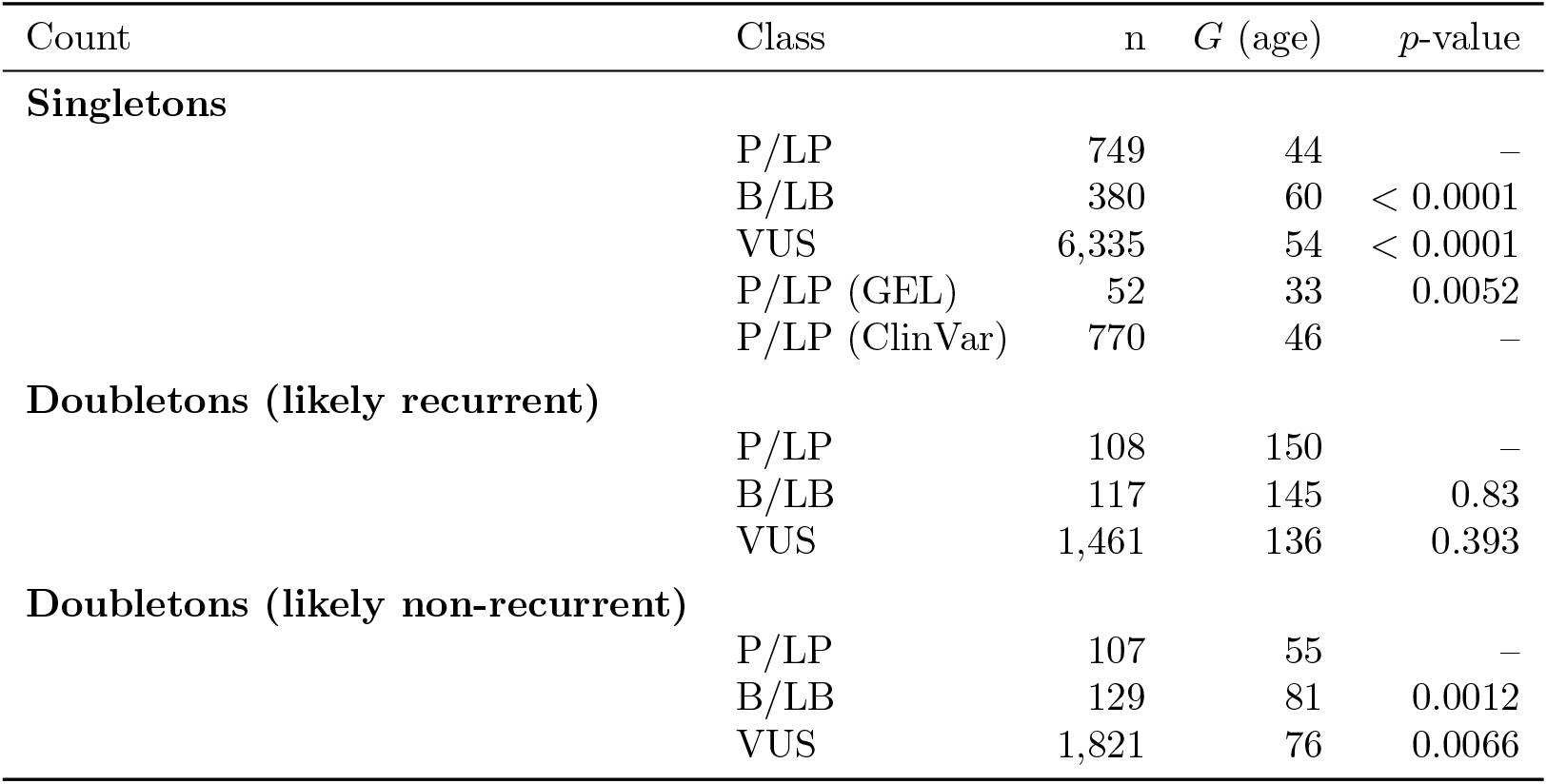
Allele age comparisons by clinical class. *G* denotes the geometric mean allele age (in generations). For doubletons, results are shown for variants estimated to be likely recurrent and likely non-recurrent using a genealogical consistency filter (Methods §23). Statistical tests evaluate whether pathogenic or likely pathogenic (P/LP) variants are younger than the comparison class using permutation tests.

| Count | Class | n | $G$ (age) | $p$-value |
| --- | --- | --- | --- | --- |
| **Singletons** |  |  |  |  |
|  | P/LP | 749 | 44 | – |
|  | B/LB | 380 | 60 | < 0.0001 |
|  | VUS | 6,335 | 54 | < 0.0001 |
|  | P/LP (GEL) | 52 | 33 | 0.0052 |
|  | P/LP (ClinVar) | 770 | 46 | – |
| **Doubletons (likely recurrent)** |  |  |  |  |
|  | P/LP | 108 | 150 | – |
|  | B/LB | 117 | 145 | 0.83 |
|  | VUS | 1,461 | 136 | 0.393 |
| **Doubletons (likely non-recurrent)** |  |  |  |  |
|  | P/LP | 107 | 55 | – |
|  | B/LB | 129 | 81 | 0.0012 |
|  | VUS | 1,821 | 76 | 0.0066 |

**Table S6:** Example of estimated ancient doubletons (DAC=2) classified as variants of uncertain significance (VUS) in the GEL dataset. All 6 are classified as Missense variants. Estimated age is reported in generations before the present.

| Chr | Pos | rsID | Gene | Gene MOI | Ggroup | Age | 95% CI |
| --- | --- | --- | --- | --- | --- | --- | --- |
| 21 | 44,525,821 | rs139112987 | TSPEAR | Recessive | AFR | 35,340 | 6,148–89,540 |
| 22 | 20,090,255 | rs777348779 | DGCR8 | Dominant | EAS | 21,657 | 3,028–58,215 |
| 16 | 580,192 | rs1361495767 | PIGQ | Recessive | NFE | 9,465 | 728–29,108 |
| 17 | 75,842,559 | rs551855408 | UNC13D | Recessive | AFR | 5,596 | 527–16,503 |
| 19 | 5,110,736 | rs1466512977 | KMD4B | Dominant | NFE | 5,369 | 333–17,197 |
| 17 | 77,488,314 | rs762543093 | SEPTIN9 | Dominant | NFE | 4,828 | 1,209–10,903 |

**Table S7:** Per-region tree sequence statistics for the 1000 Genomes Project dataset. The regions are a subset of those detailed in Tab. S2. See Tab. S4 for details on the columns shown.

| region | length (bp) | sites | trees | nodes | edges | mut | file size |
| --- | --- | --- | --- | --- | --- | --- | --- |
| 17p | 17,994,697 | 596,776 | 265,521 | 528,258 | 3,482,633 | 596,776 | 218.5MB |
| 17q1 | 9,163,785 | 263,697 | 105,366 | 203,601 | 1,171,136 | 263,697 | 82.0MB |
| 17q2 | 1,337,198 | 40,576 | 15,856 | 37,064 | 198,920 | 40,576 | 14.0MB |
| 17q3 | 7,230,866 | 211,258 | 80,931 | 163,430 | 933,648 | 211,258 | 65.6MB |
| 17q4 | 37,497,391 | 1,162,183 | 465,421 | 937,537 | 5,941,940 | 1,162,183 | 389.8MB |
| 20p | 25,524,955 | 849,268 | 350,460 | 661,927 | 4,065,850 | 849,268 | 273.9MB |
| 20q | 34,481,295 | 1,087,525 | 441,943 | 847,444 | 5,102,012 | 1,087,525 | 347.0MB |
| Total | 133,230,187 | 4,211,283 | 1,725,498 | 3,379,261 | 20,896,139 | 4,211,283 | 1.4GB |

**Table S8:**
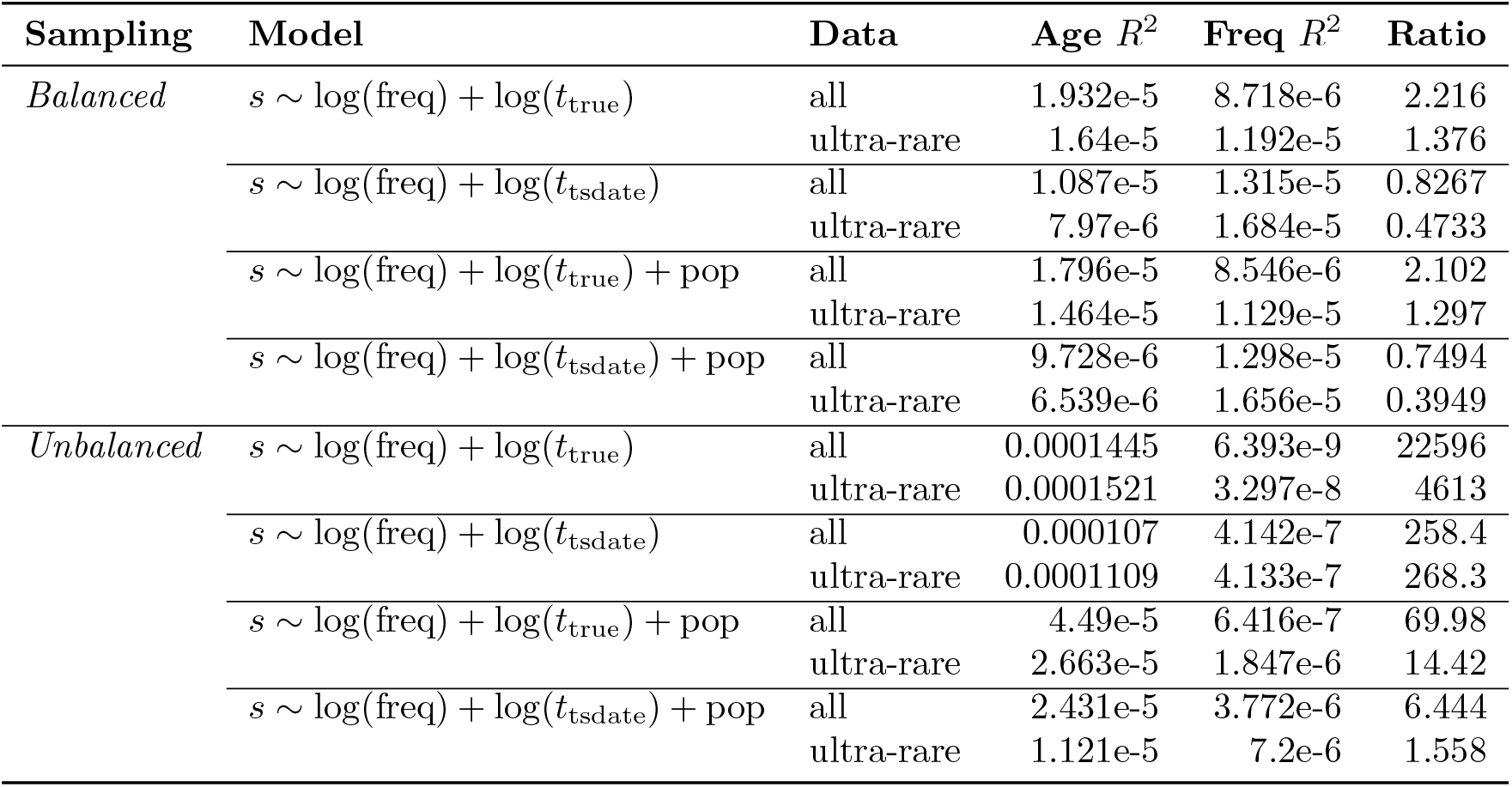
Variance explained by mutation age versus allele frequency in predicting negative selection. Proportional reduction in deviance (pseudo *R*^2^) when age or frequency terms are removed from linear models that predict the selection coefficient (*s*). Two simulated datasets were used, comprising 60,000 samples of a 40Mb region of human chromosome 17.

| Sampling | Model | Data | Age $R^2$ | Freq $R^2$ | Ratio |
| --- | --- | --- | --- | --- | --- |
| *Balanced* | $s \sim \log(\text{freq}) + \log(t_{\text{true}})$ | all | 1.932e-5 | 8.718e-6 | 2.216 |
|  |  | ultra-rare | 1.64e-5 | 1.192e-5 | 1.376 |
| | $s \sim \log(\text{freq}) + \log(t_{\text{tsdate}})$ | all | 1.087e-5 | 1.315e-5 | 0.8267 |
|  |  | ultra-rare | 7.97e-6 | 1.684e-5 | 0.4733 |
| | $s \sim \log(\text{freq}) + \log(t_{\text{true}}) + \text{pop}$ | all | 1.796e-5 | 8.546e-6 | 2.102 |
|  |  | ultra-rare | 1.464e-5 | 1.129e-5 | 1.297 |
| | $s \sim \log(\text{freq}) + \log(t_{\text{tsdate}}) + \text{pop}$ | all | 9.728e-6 | 1.298e-5 | 0.7494 |
|  |  | ultra-rare | 6.539e-6 | 1.656e-5 | 0.3949 |
| *Unbalanced* | $s \sim \log(\text{freq}) + \log(t_{\text{true}})$ | all | 0.0001445 | 6.393e-9 | 22596 |
|  |  | ultra-rare | 0.0001521 | 3.297e-8 | 4613 |
| | $s \sim \log(\text{freq}) + \log(t_{\text{tsdate}})$ | all | 0.000107 | 4.142e-7 | 258.4 |
|  |  | ultra-rare | 0.0001109 | 4.133e-7 | 268.3 |
| | $s \sim \log(\text{freq}) + \log(t_{\text{true}}) + \text{pop}$ | all | 4.49e-5 | 6.416e-7 | 69.98 |
|  |  | ultra-rare | 2.663e-5 | 1.847e-6 | 14.42 |
| | $s \sim \log(\text{freq}) + \log(t_{\text{tsdate}}) + \text{pop}$ | all | 2.431e-5 | 3.772e-6 | 6.444 |
|  |  | ultra-rare | 1.121e-5 | 7.2e-6 | 1.558 |

**Figure S1:**
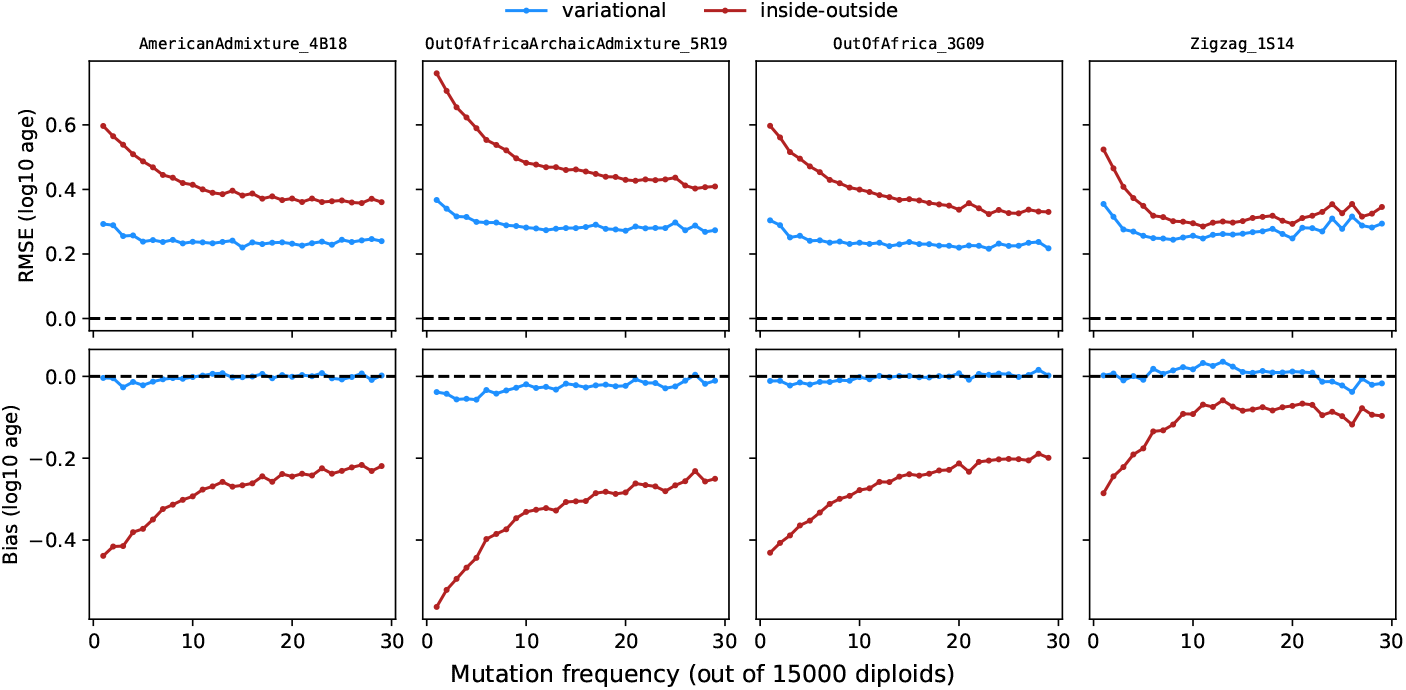
Accuracy of the algorithm described in this study (“variational”) compared to the previous version in tsdate (“inside-outside”). Accuracy was assessed by comparing true (simulated) and inferred (tsinfer and tsdate) ARGs across four human stdpopsim models, over a 40 Mb section of chromosome 17 with 15,000 diploids sampled evenly across populations. Root mean square error (RMSE) and bias are calculated for log_10_ mutation ages (estimated as edge midpoints), separately for mutations at various frequencies in the sample.

**Figure S2:**
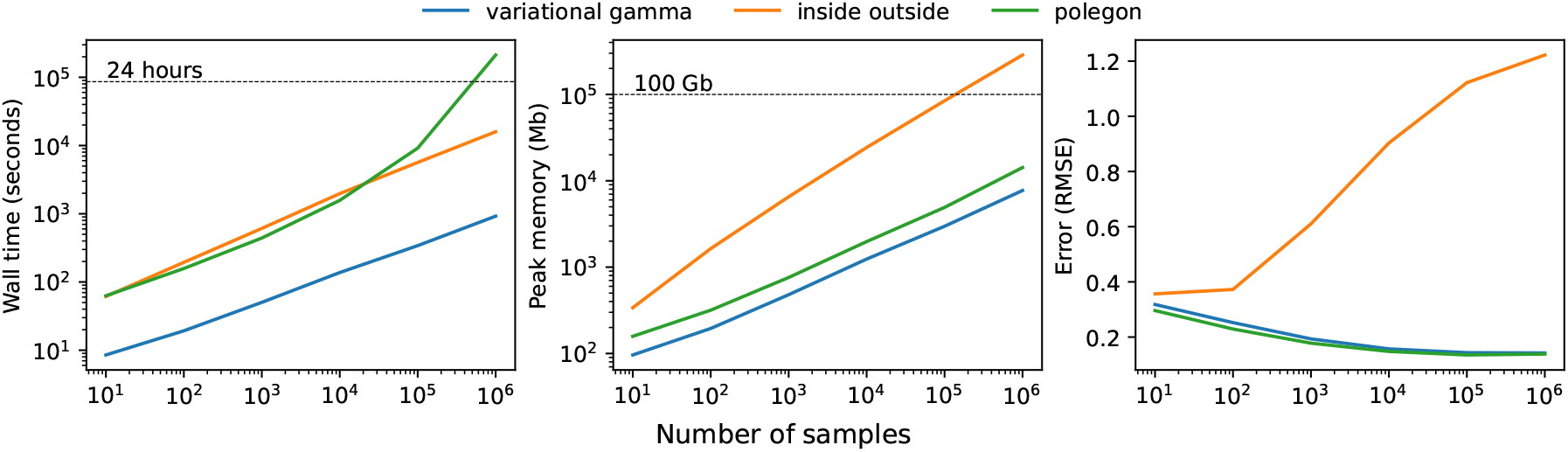
Benchmarks of the `variational gamma` algorithm, alongside the original `inside outside` algorithm in tsdate and POLEGON. All three methods were applied to a realistic, pedigree-based simulation of human chromosome 22, randomly subsampled to various numbers of samples From left to right are (median of three replicates) wall time in seconds, peak memory usage in Mb, and root mean squared error of estimated log_10_ node ages.

**Figure S3:**
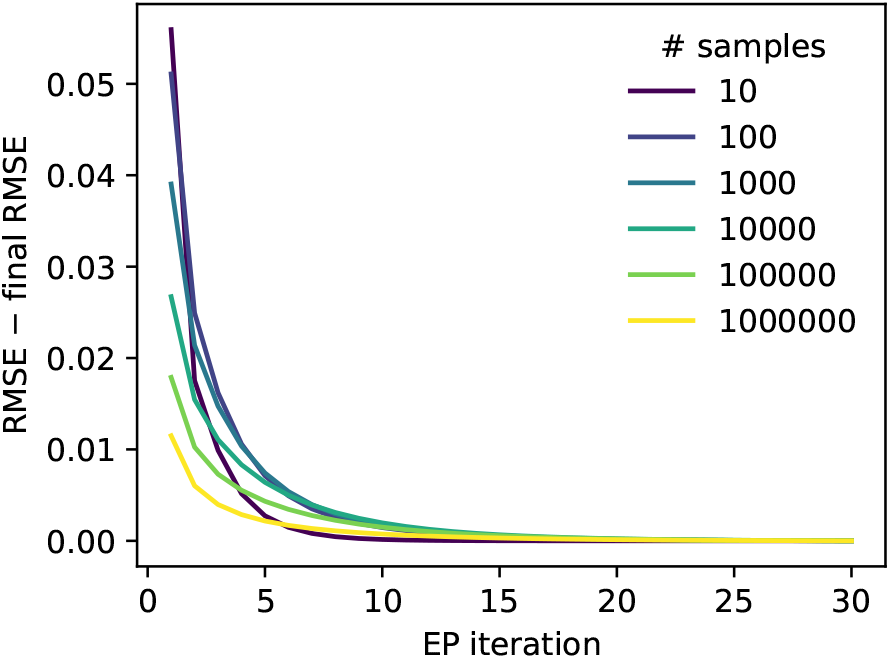
Convergence of the tsdate `variational gamma` algorithm with iteration of the expectation propagation scheme. Convergence was measured as root mean square error (RMSE) in node ages at a given iteration, minus the RMSE at the final iteration, averaged over three replicate ARGs per sample size. Each replicate was generated by randomly subsampling a large, pedigree-based simulation of human chromosome 22.

**Figure S4:**
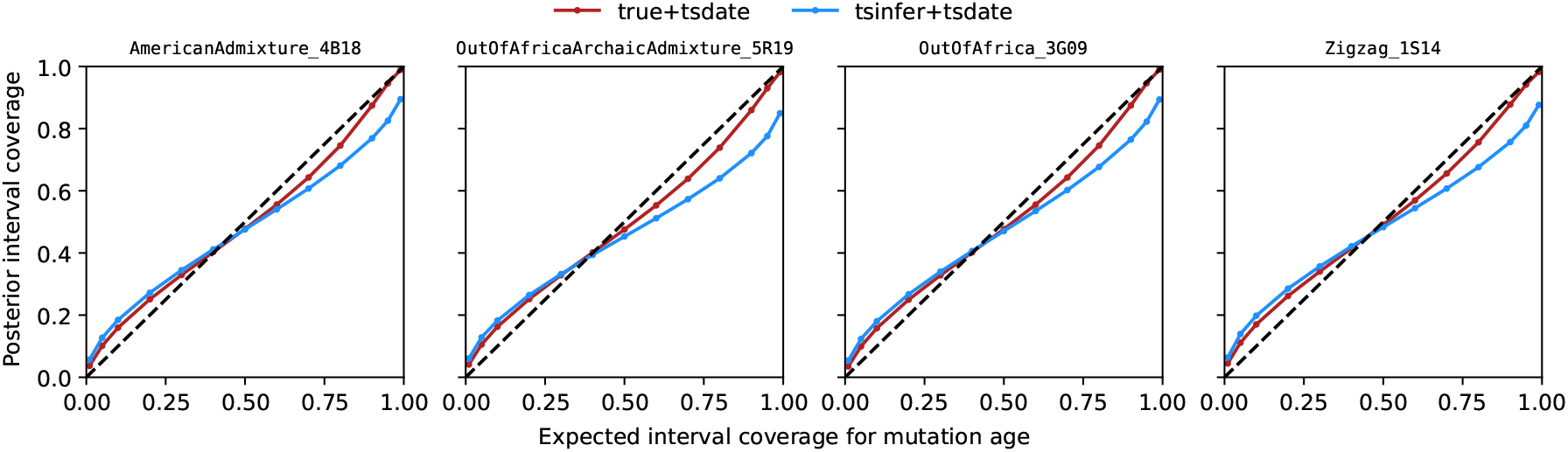
Calibration of the approximate posteriors for mutation ages produced by the ts-date `variational_gamma` algorithm. tsdate was applied to both true (simulated) and inferred (tsinfer) ARGs across four human stdpopsim models, over a 40 Mb section of chromosome 17 with 15,000 diploids sampled evenly across populations. Calibration can be quantified by comparing the proportion of true ages that fall within posterior intervals to what is expected given the interval widths, a perfectly calibrated method would fall on the *y* = *x* line in these plots. The approximate posteriors from the true ARGs provide reasonable estimates of uncertainty, especially for the widest intervals, despite the assumption of a particular parametric form. In contrast, the approximate posteriors from the inferred ARGs underestimate uncertainty – for instance, a 99% interval is roughly equivalent to a 90% interval. This is expected, because the uncertainty reported by tsdate does not model the additional error from the tsinfer topology inference.

**Figure S5:**
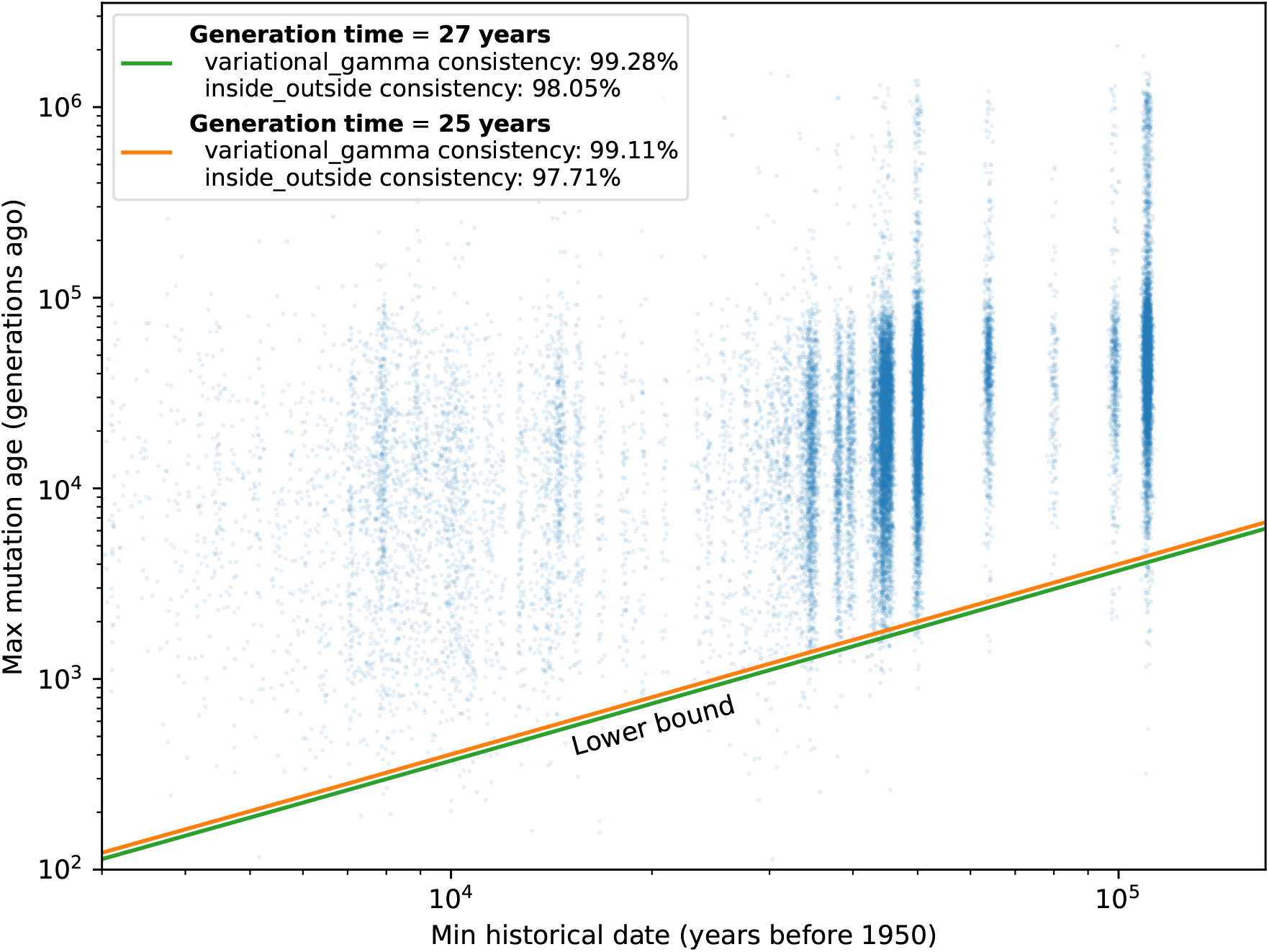
Allelic ages inferred by tsinfer+tsdate from the 1000 Genomes dataset (chromosome 20) compared to minimum variant ages from the Allen Ancient DNA Resource. See Methods §8 for details. Ancient DNA dates range from 0.1ka to 111ka.

**Figure S6:**
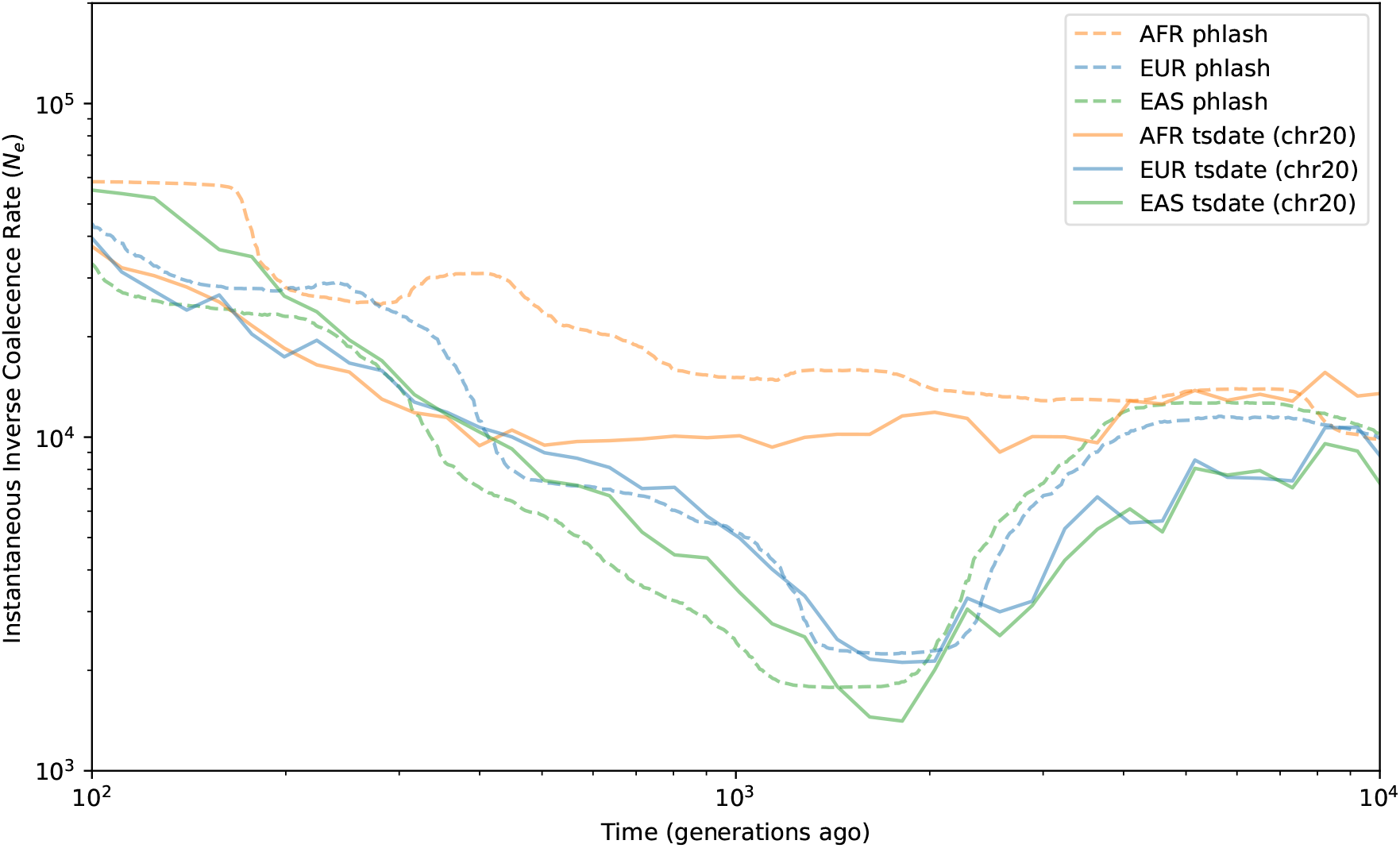
Inverse instantaneous coalescence rates (*N*_*e*_) inferred by tsinfer+tsdate from the 1000 Genomes dataset (chromosome 20), focussing on the proposed time period for the Out of Africa bottleneck. Previously published estimates from PHLASH are also shown.^51^ See Methods §9 for details. The plot spans the time period 2.7ka to 270ka (assuming a generation time of 27 years).

**Figure S7:**
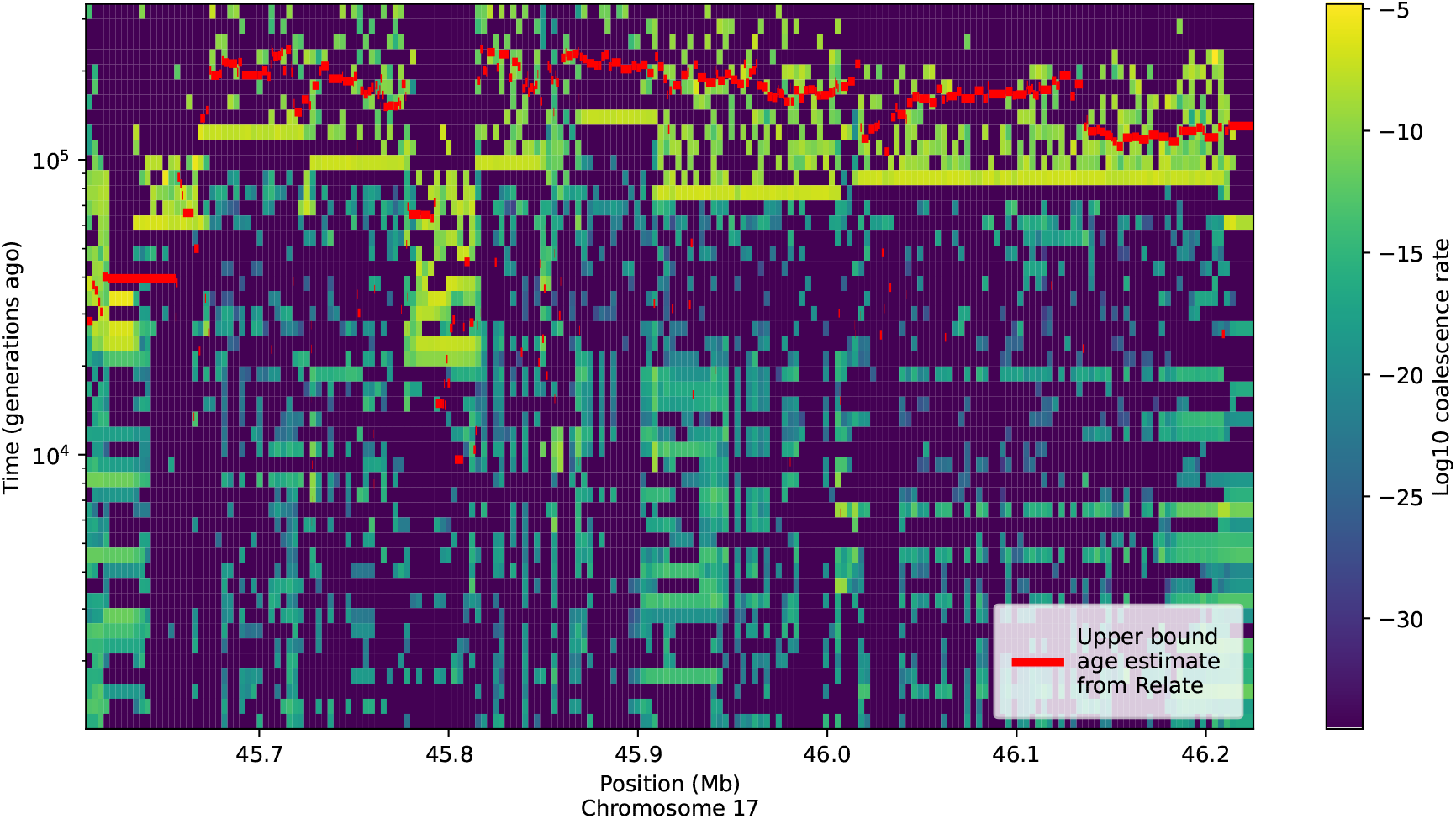
Dated cross coalescence rates inferred by tsinfer+tsdate between carriers and non-carriers of a known ancient inversion on chromosome 17 of the 1000 Genomes dataset. The upper bounds for the inversion from a published estimate which range from 160 ka to 6417 ka (assuming a generation time of 27 years), are shown in red. See Methods §10 for details.

**Figure S8:**
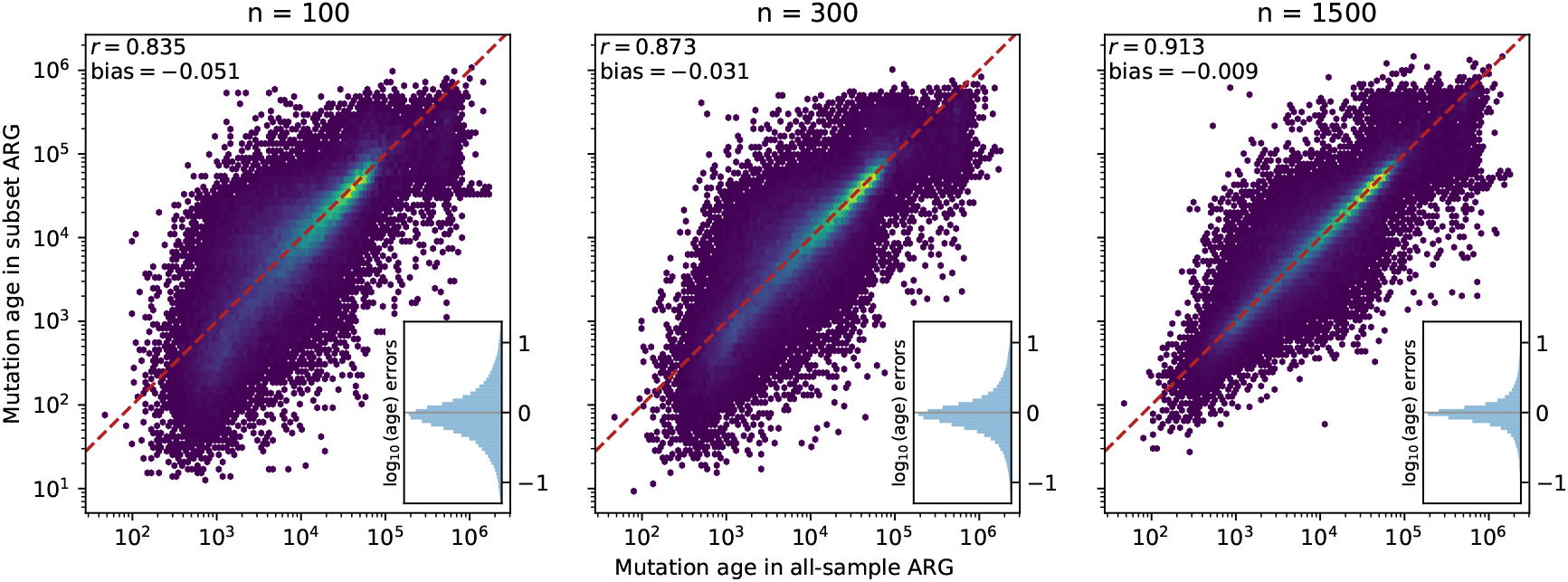
Dependence of mutation age estimates on sample size in the 1000 Genomes Project data. Mutation age estimates based on ARGs inferred from three subsets (100, 300 and 1500 samples) compared to estimates from the complete dataset. See Methods §7 for details.

**Figure S9:**
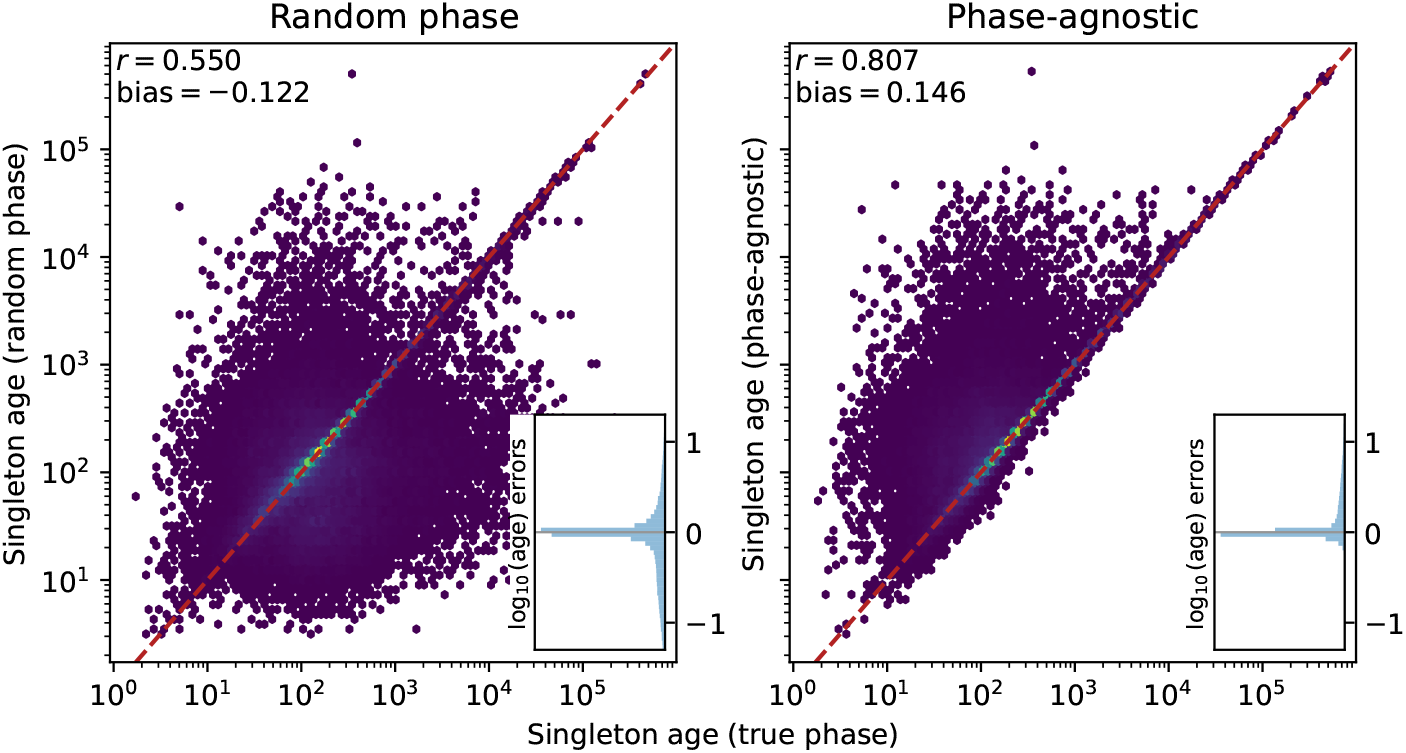
Validation of phase-agnostic singleton mutation dating using the 1000 Genomes Project data. We used 106,863 sites that are singletons in an ARG inferred from a subset of 1,500 individuals, but of higher frequency in the all-sample ARG. Age estimates of singletons obtained using their true phase are compared to those using a randomly assigned phase (left) and the phase-agnostic dating algorithm (right). See Methods §7 for details.

**Figure S10:**
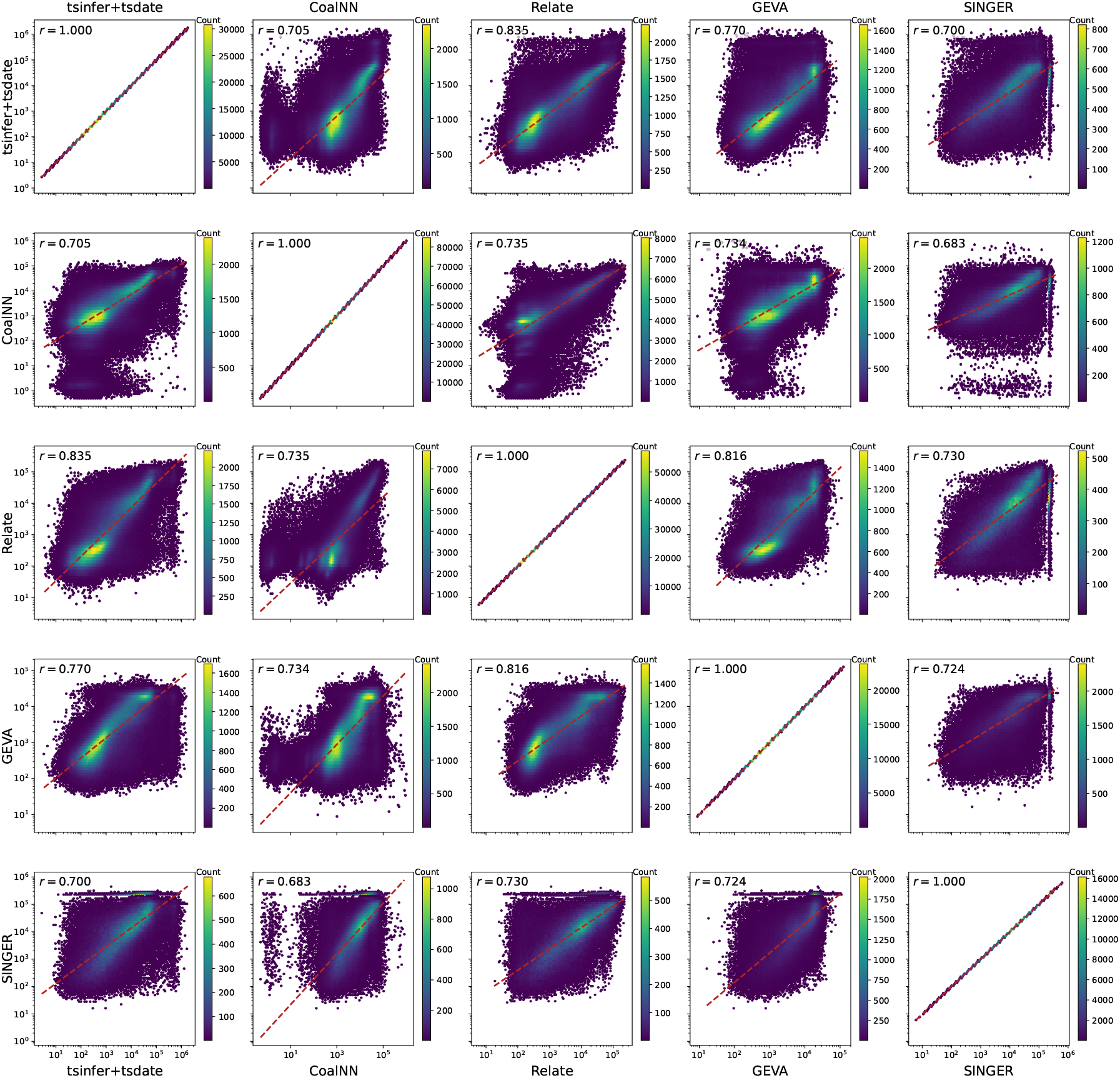
Pairwise comparison of allele age estimates obtained from five methods using chromosome 20 of the 1000 Genomes Project data. Results are restricted to a subset of 162,556 sites that are present in all datasets and have identical allele polarisation. See Methods §11 for details.

**Figure S11:**
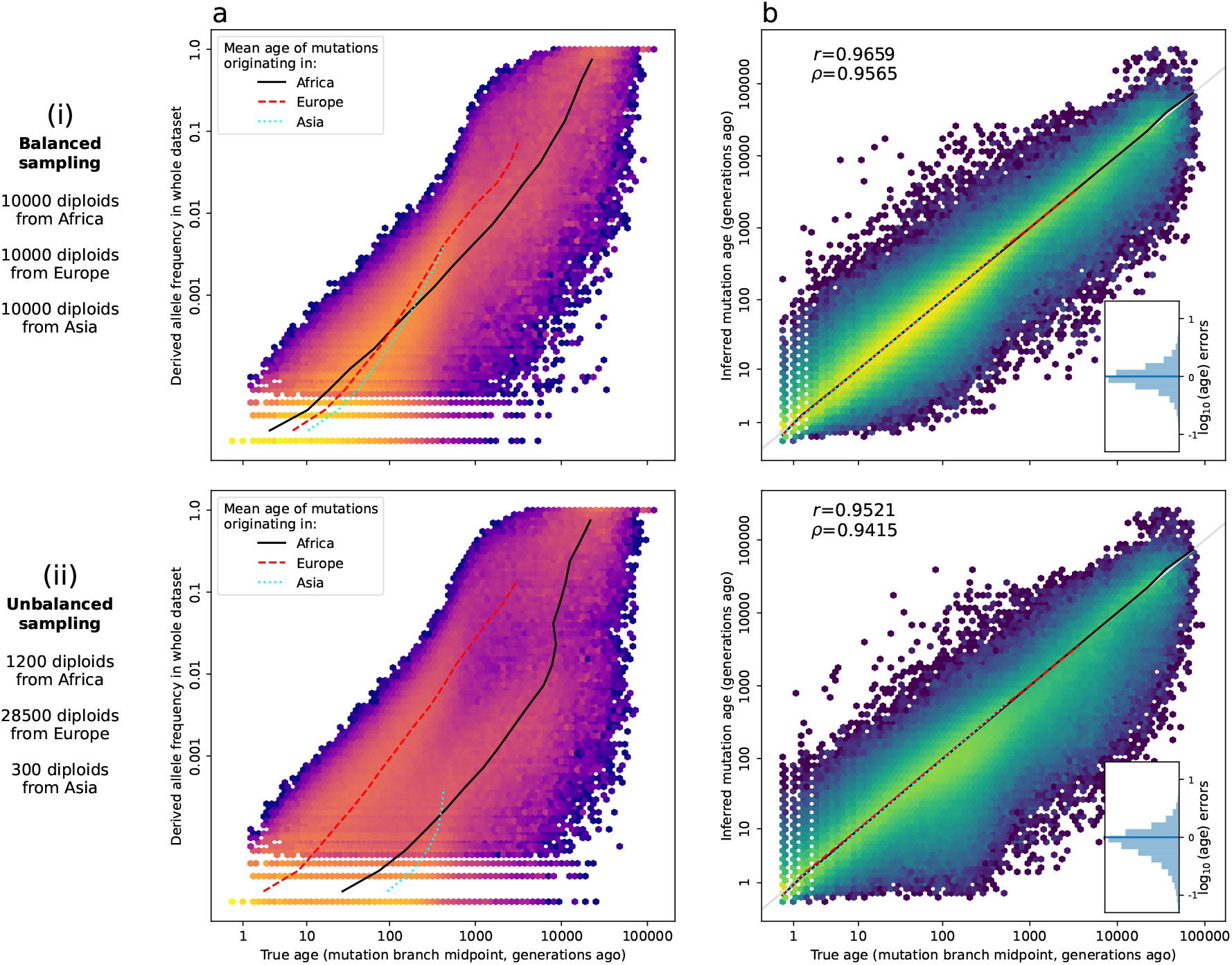
Robustness to unbalanced sampling. Distributions of mutation ages and frequencies in a 40 MB segment of human chromosome 17 from a 3-population Out-of-Africa simulation with negative selection under balanced (top row) and unbalanced (bottom row) sampling regimes. Data is restricted to the 98% of polymorphic sites with only one mutation. (left plots) Allele frequency against true age; solid/dotted/dashed lines lines indicate binned mean mutation ages for the 3 populations in 20 evenly-spaced log-frequency bins. (right plots) Inferred age against true age; lines as in (i), and residual inferred ages in inset histograms. Although the relationship between age and frequency breaks down with unbalanced sampling (compare bottom-left to top-left plot), age estimation is not affected (compare bottom-right and top-right plots).

**Figure S12:**
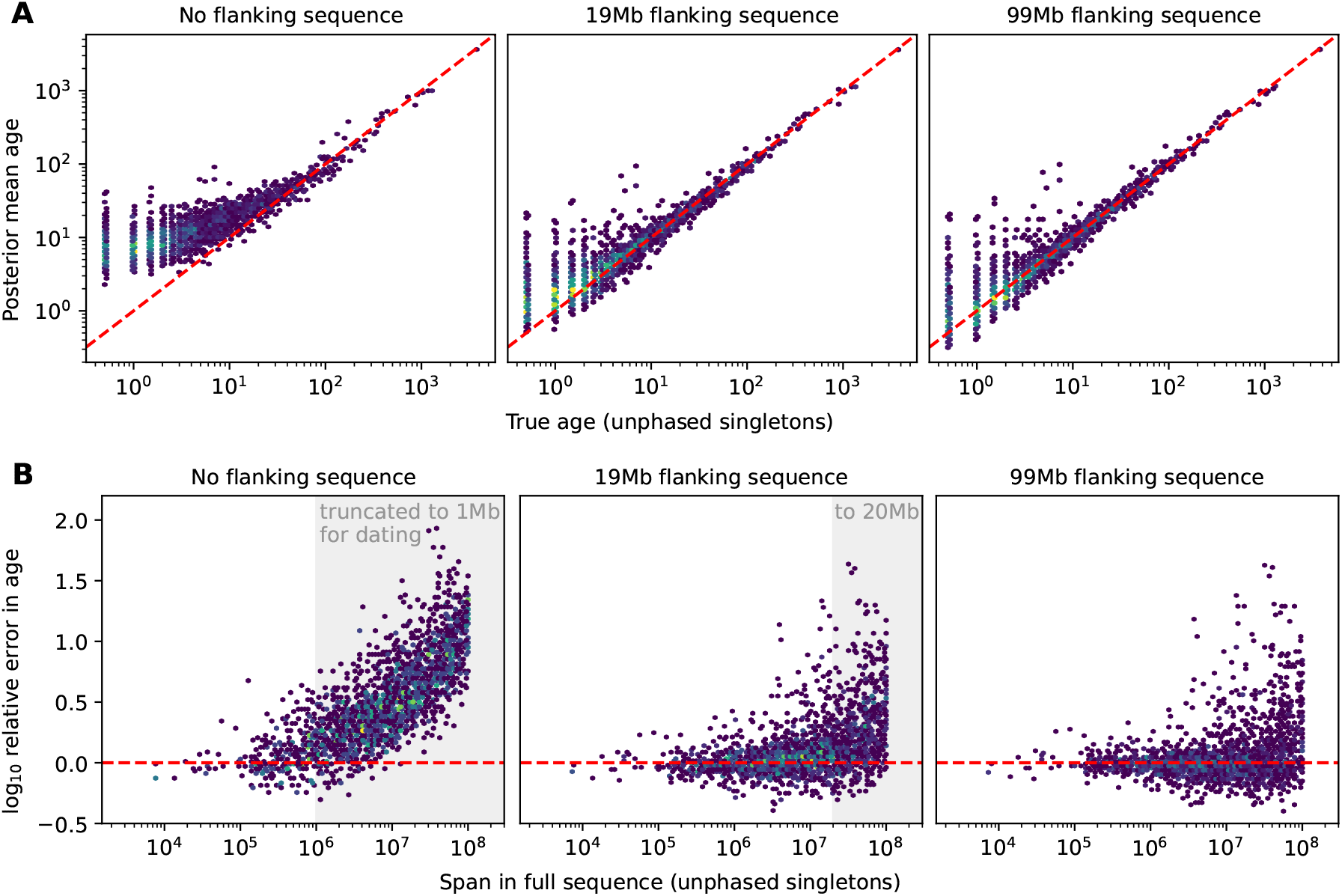
ARGs with short sequence lengths lead to bias in the estimated ages of recent mutations. (A) True versus estimated ages of unphased singleton mutations, from a 1Mb interval in the center of a 100Mb simulated tree sequence with 40k haploids. Each panel shows the same mutations, dated on subsegments of the tree sequence of progressively larger lengths (1Mb, 20Mb, and 100Mb). Substantial upwards bias in the youngest mutation ages is evident for shorter sequences. (B) The error in estimated ages versus the span of the edge. The singletons in the gray region are on edges that must have been truncated by removing the flanking regions prior to dating. Thus, truncating long, recent haplotypes leads to substantial dating error.

**Figure S13:**
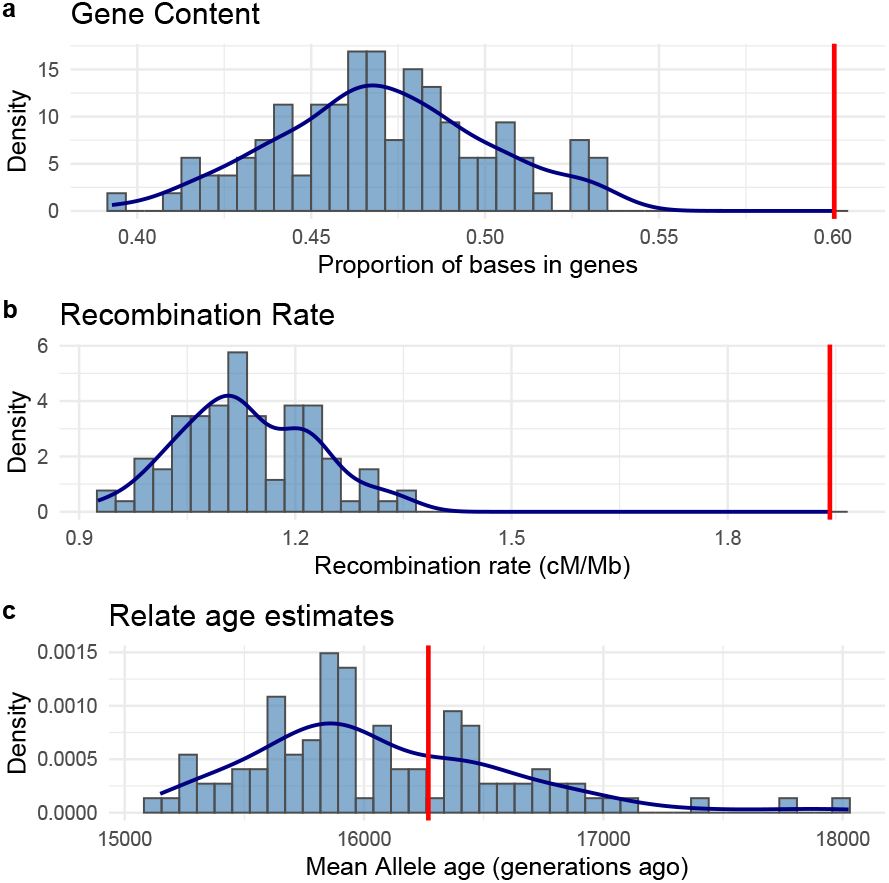
Comparison of genomic properties between the GEL ARG inference regions (red line; Table S2) and matched random autosomal regions (blue bars). The GEL estimate aggregates values across all 15 inference regions. Each blue bar corresponds to one of 100 independent replicates, each comprising 15 regions matched to the inference regions by size. Panels show (a) gene content (fraction of bases within ENSEMBL protein-coding genes), (b) average recombination rate (cM/Mb; HapMap II), and (c) mean allele age inferred from the Relate Yoruba (YRI) dataset.

**Figure S14:**
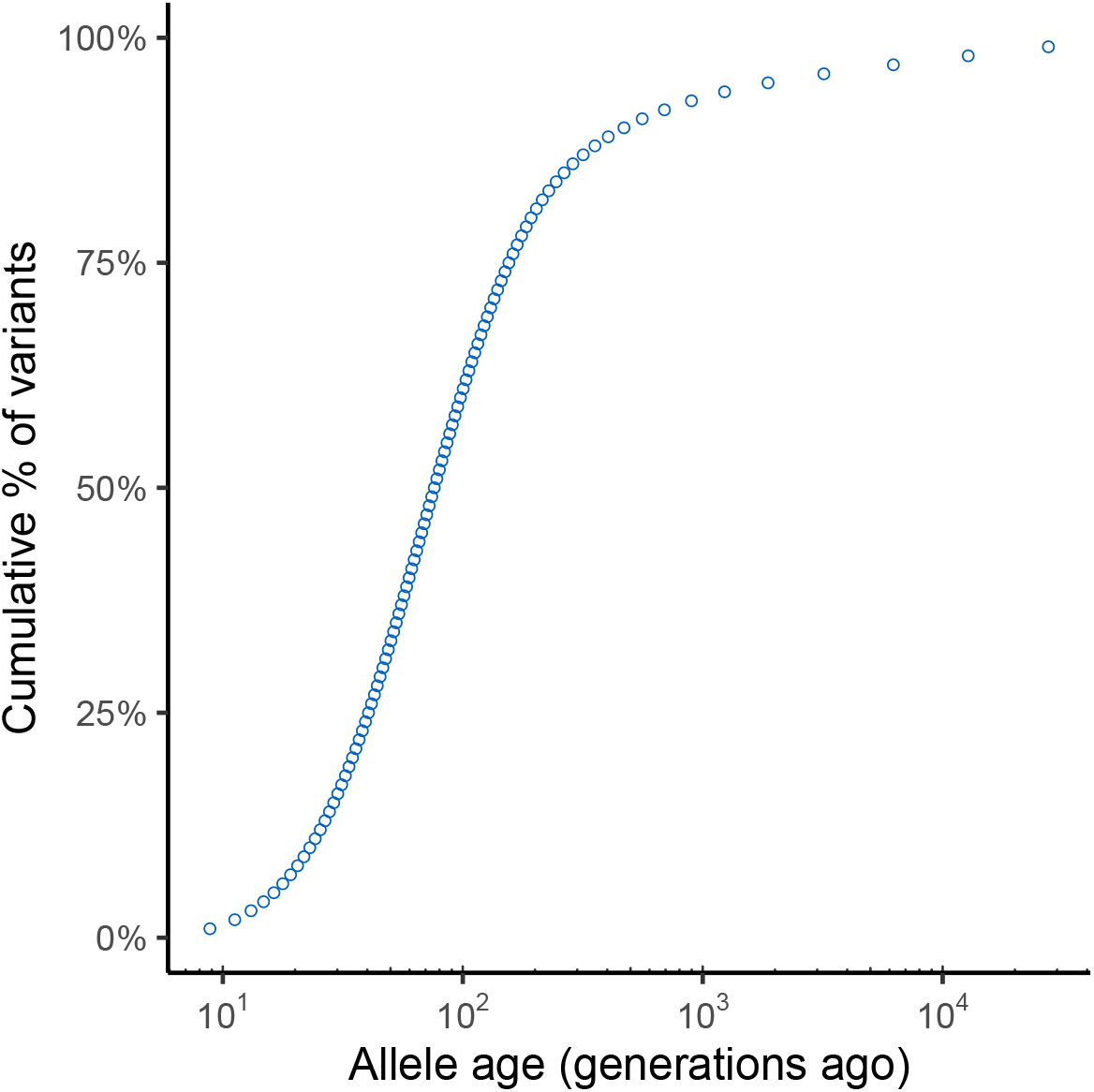
Cumulative distribution function of estimated variant ages in the GEL dataset.

**Figure S15:**
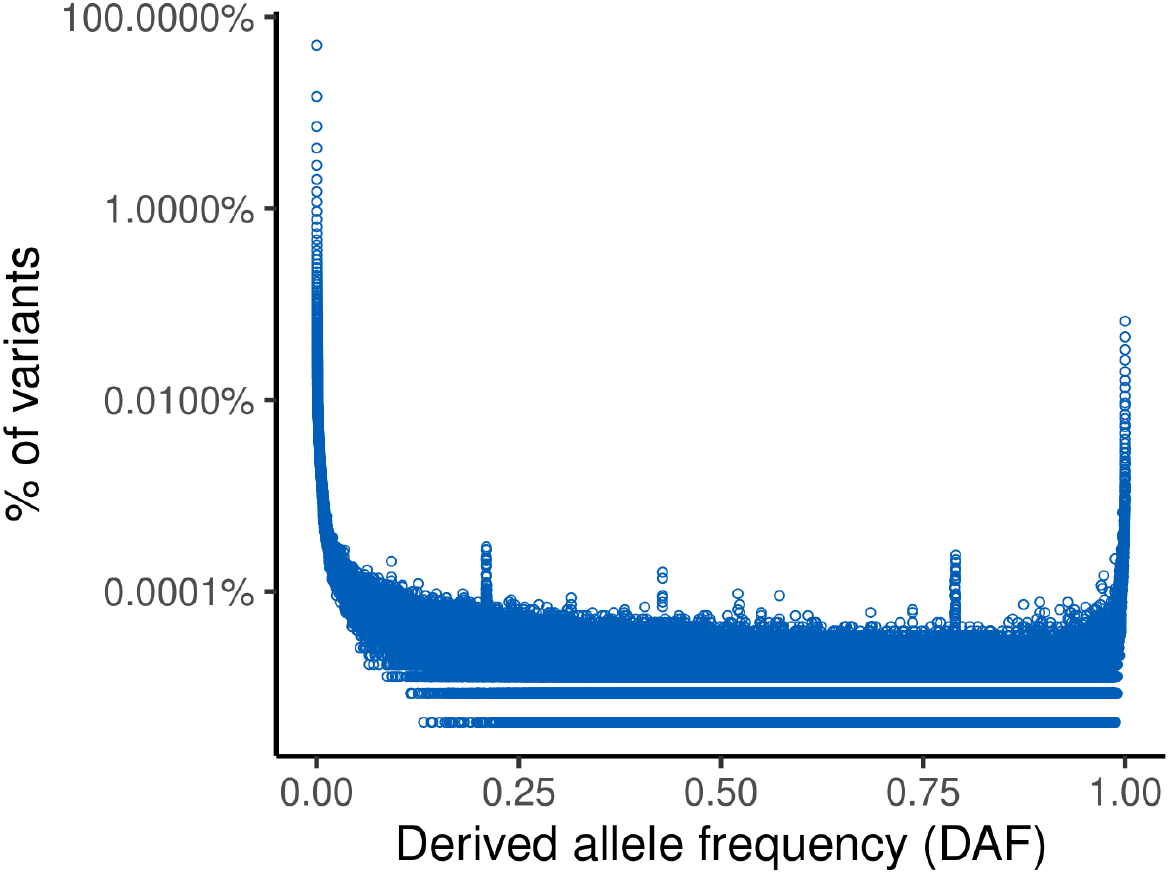
Derived allele frequency spectrum (AFS) in the GEL dataset.

**Figure S16:**
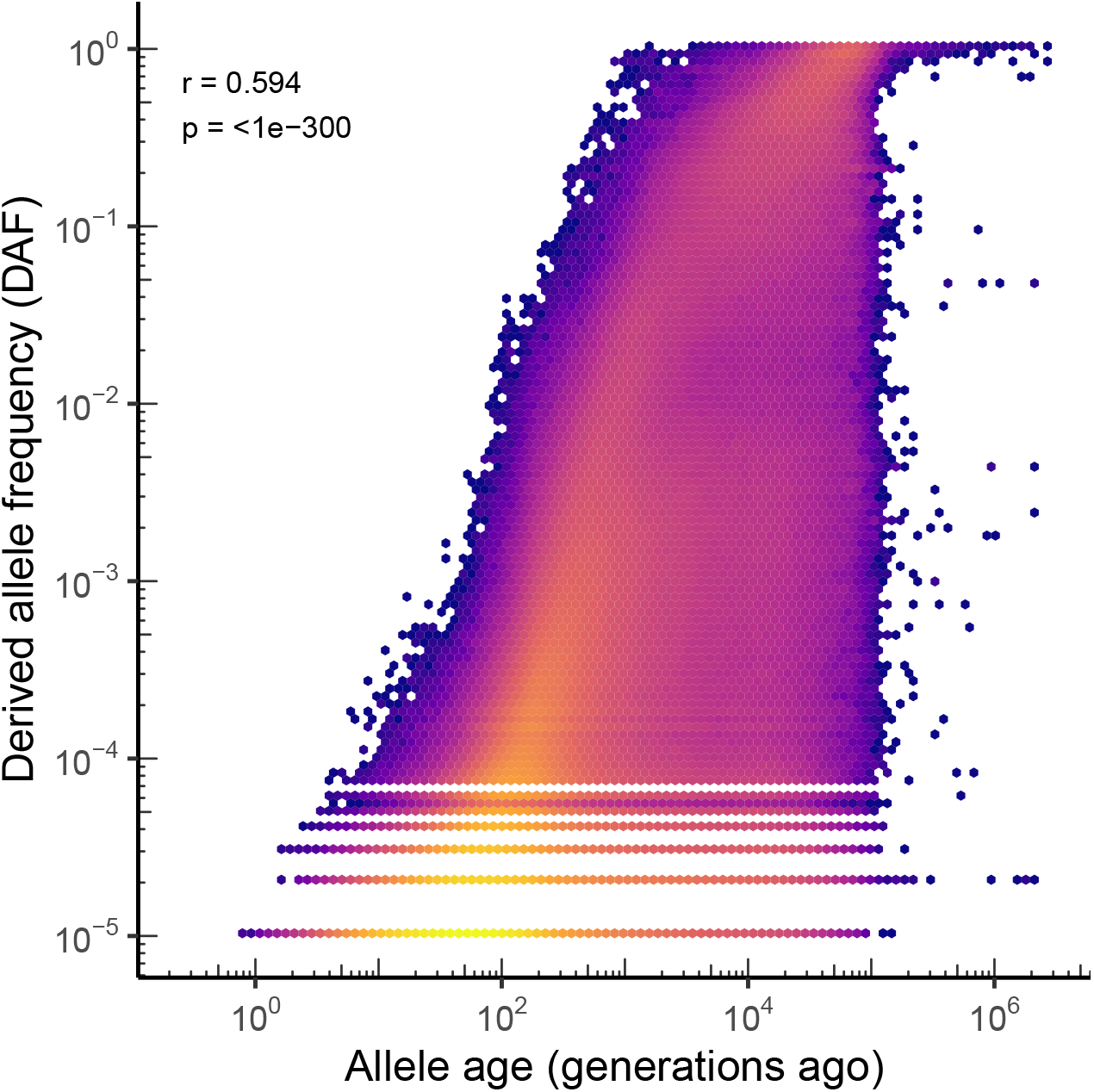
Log–log relationship between allele age (generations ago) and derived allele frequency (DAF), showing a positive correlation (*r* = 0.594, *p* < 10^−300^), consistent with younger variants tending to segregate at lower frequency.

**Figure S17:**
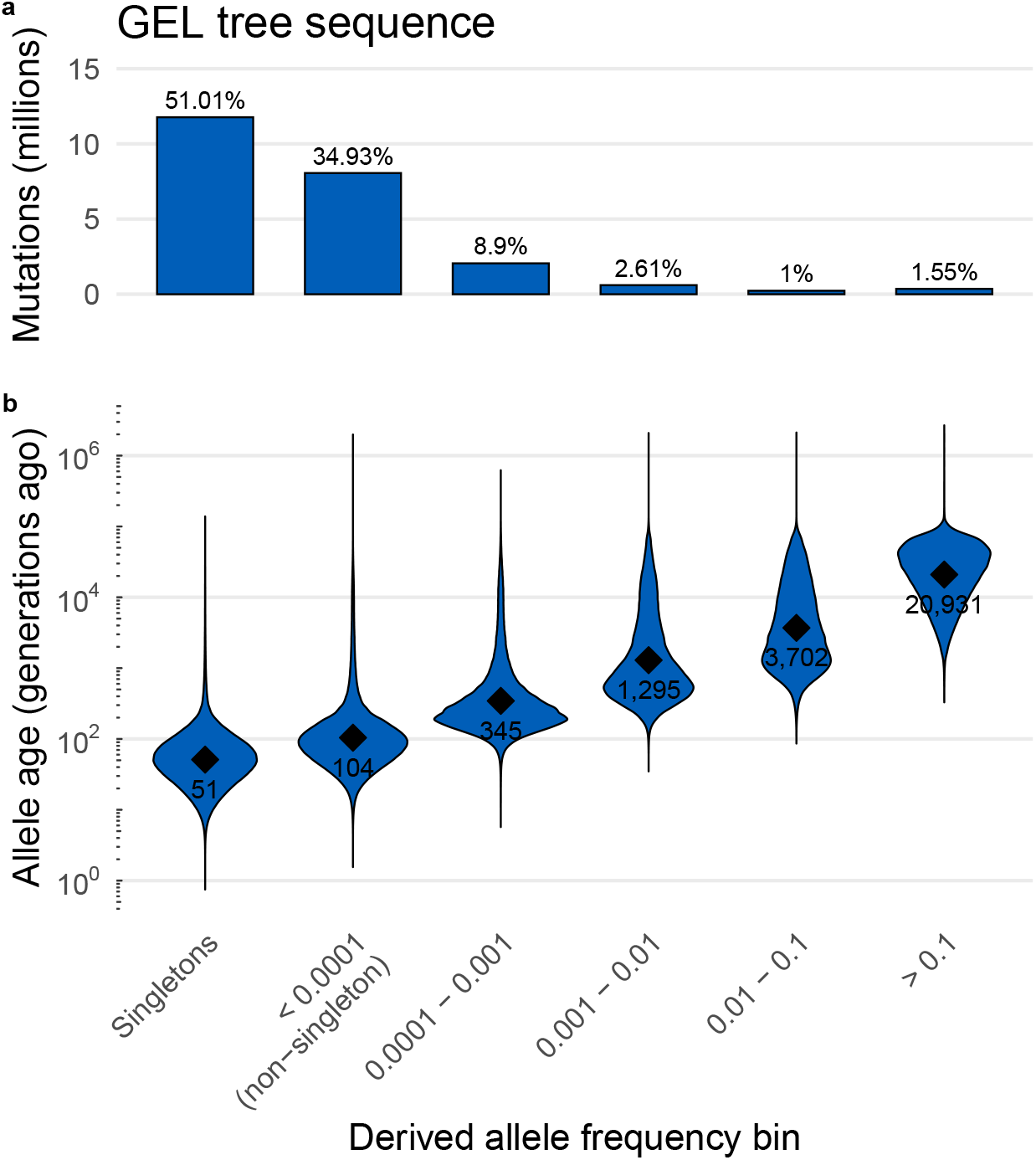
Allele age versus derived allele frequency in the GEL dataset. (a) Number of mutations per frequency bin (millions). (b) Violin distributions of allele age (generations ago) across the same bins, showing younger ages at lower frequencies.

**Figure S18:**
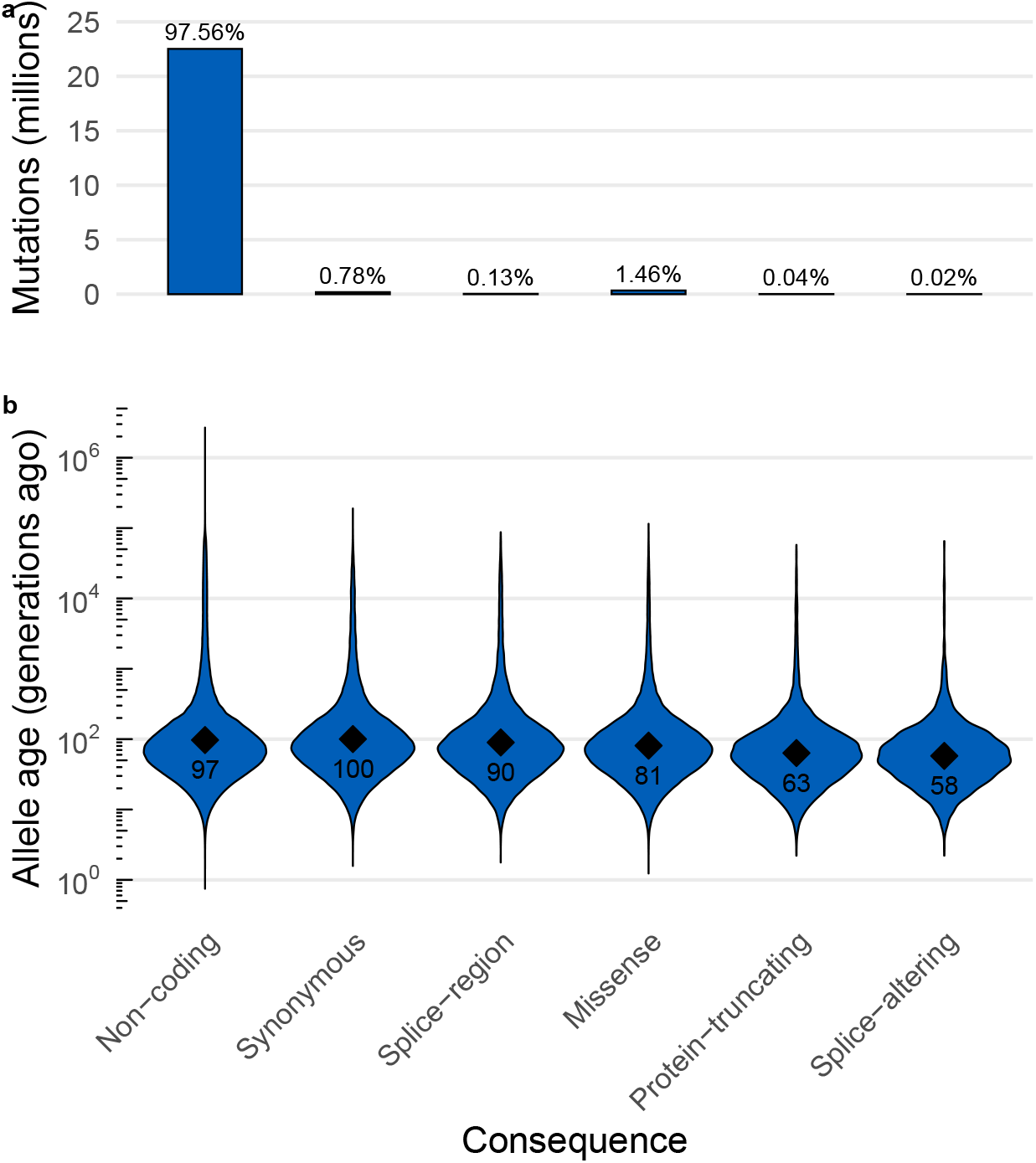
Allele age versus mutation consequence in the GEL dataset. (a) Number of mutations per consequence category (millions), ordered by severity. (b) Violin distributions of allele age (generations ago) across the same bins, showing younger ages more severe consequence types; points and labels show geometric mean ages. Categories shown (left to right): Non-coding, Synonymous, Splice-region, Missense, Protein-truncating, and Splice-altering as estimated using Ensembl VEP.

**Figure S19:**
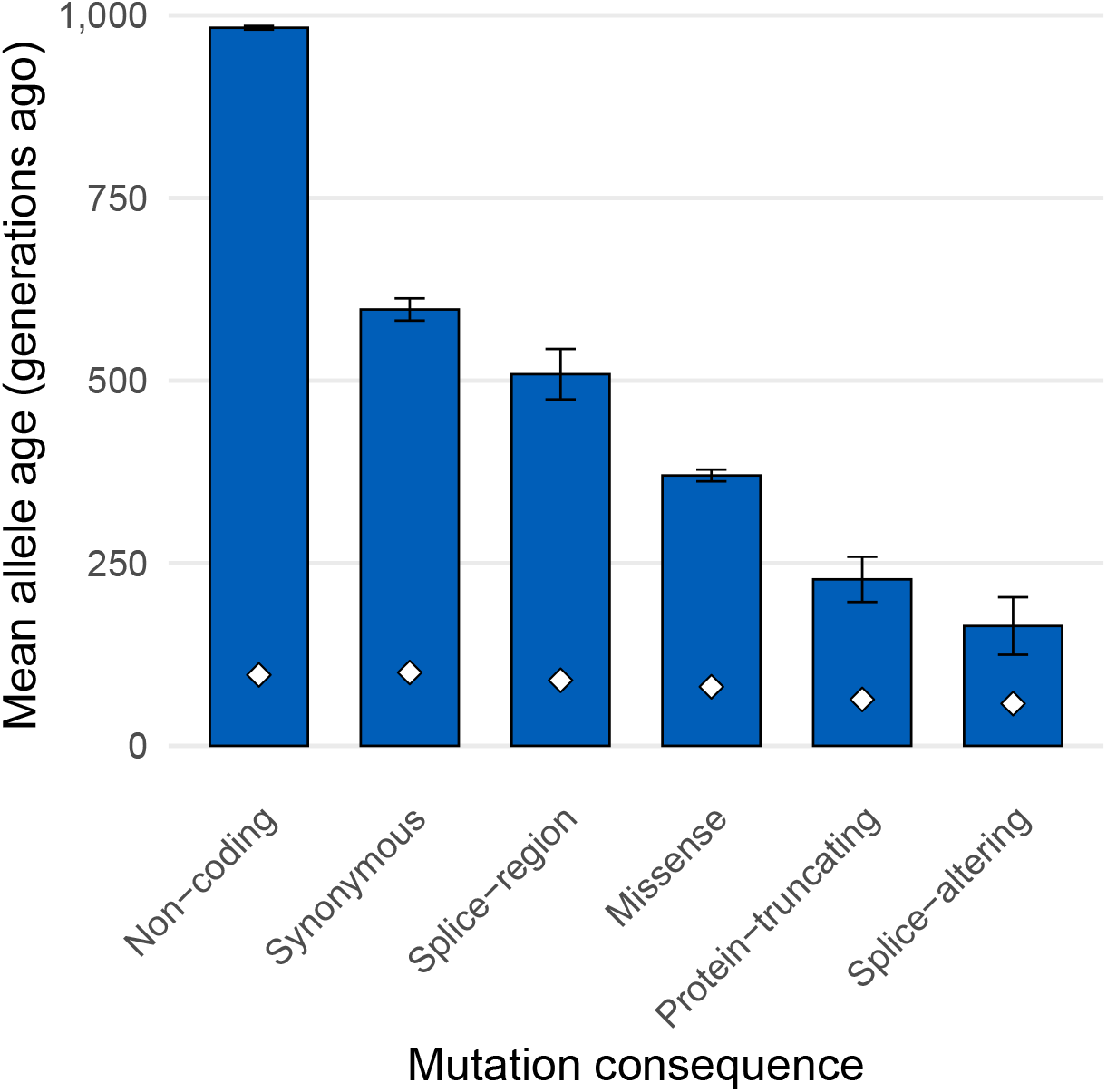
Mean allele age by mutation consequence in the GEL dataset. Bars show the arithmetic mean per category; error bars are 95% confidence intervals from a *t*-distribution (`mean cl t`). White diamond markers denote the geometric mean. Categories shown (left to right): Non-coding, Synonymous, Splice-region, Missense, Protein-truncating, and Splice-altering as estimated using Ensembl VEP. As expected from theory, the majority of all mutations across all categories are young (recent geometric mean ages), but a much smaller proportion of putatively deletrious alleles are old (strong decrease in arithmetic mean ages).

**Figure S20:**
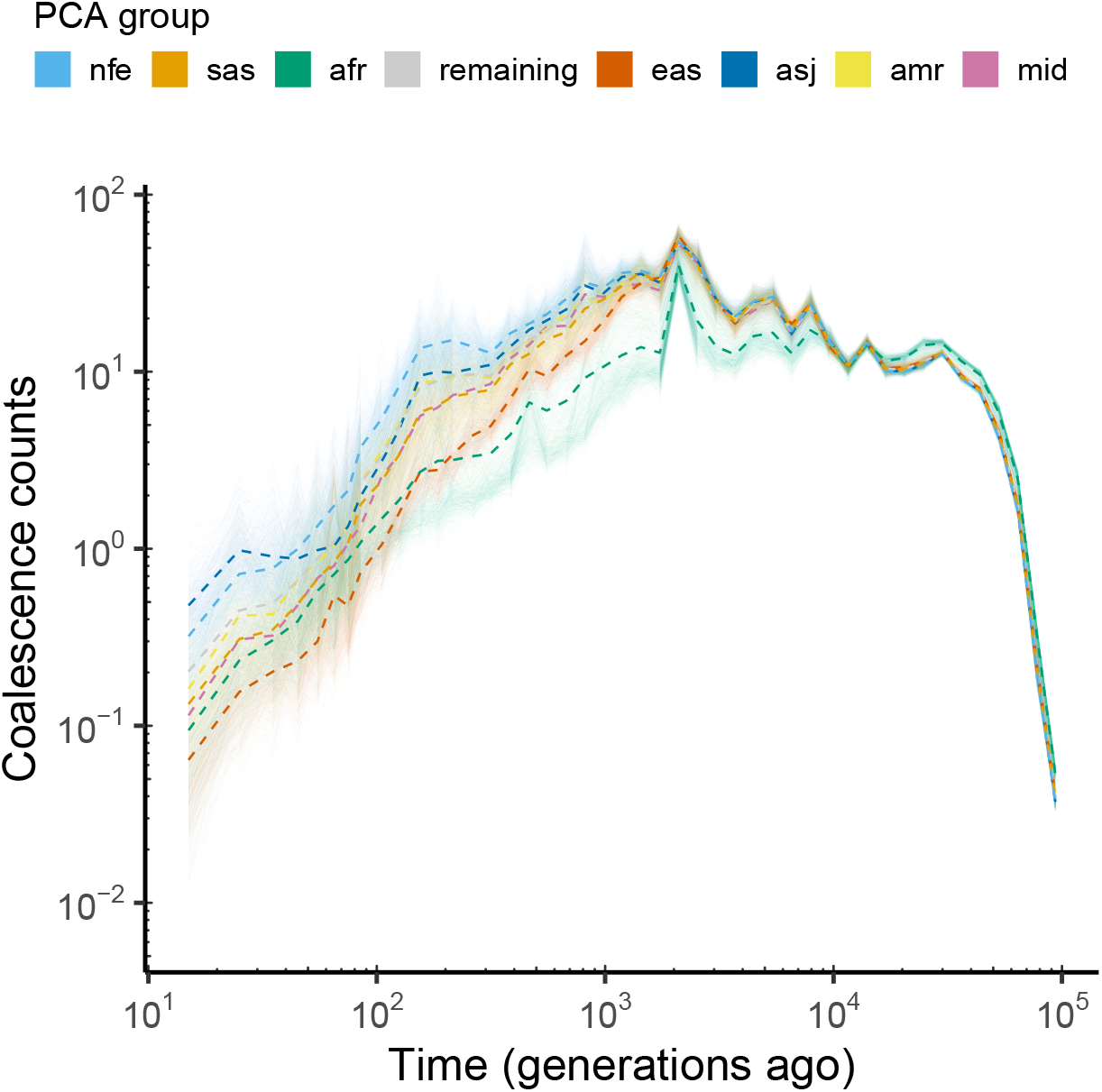
Pairwise coalescence counts between focal individuals and all others in the inferred ARG, coloured by PCA group. Each faint line represents a single focal individual (up to 500 per group, or all when *n* < 500), showing the number of coalescence events through time across the largest contiguous inferred ARG segment on chromosome 17q. Dashed lines indicate the mean for each PCA group. Both axes are shown on a log scale.

**Figure S21:**
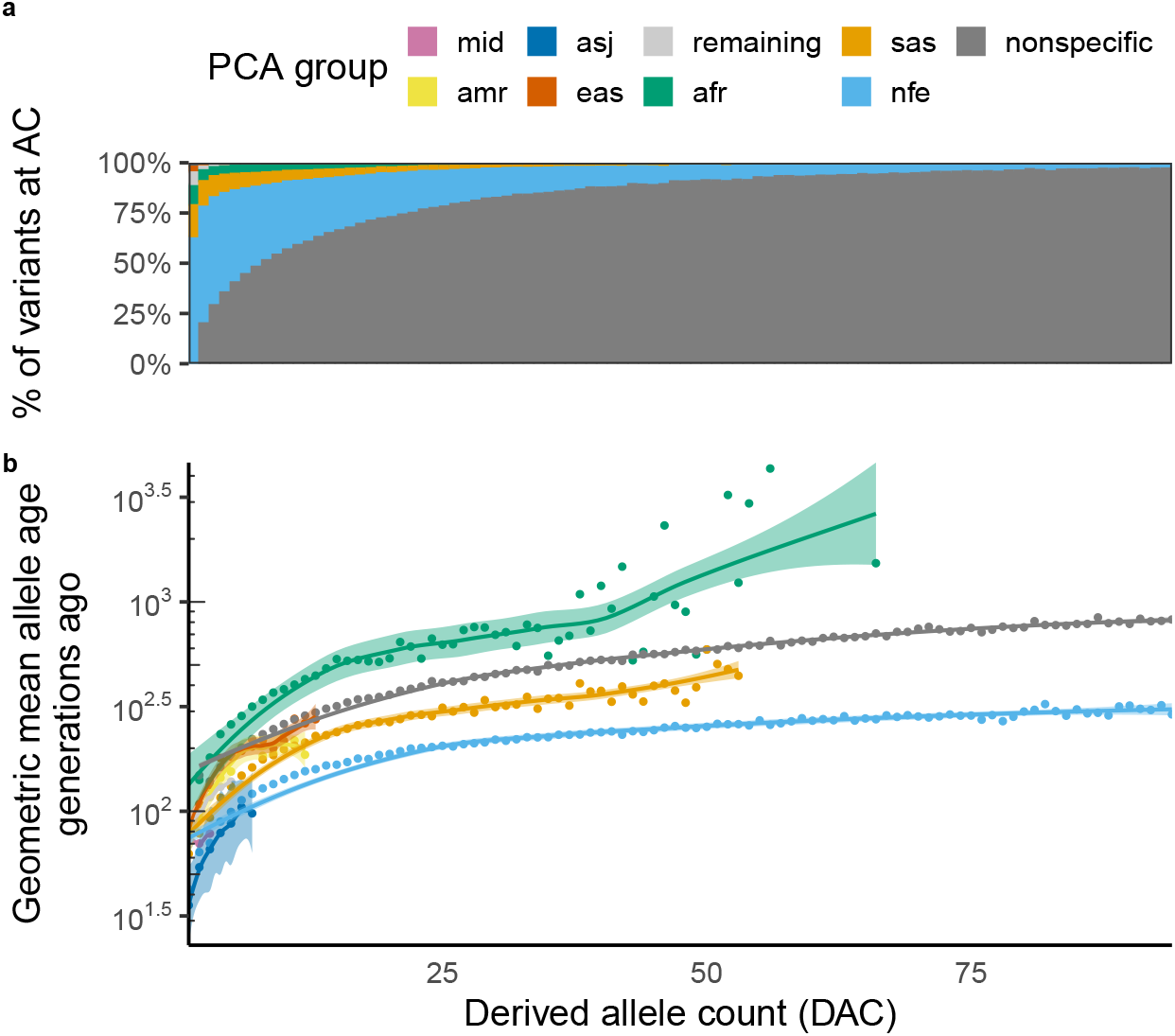
PCA group-stratified allele age versus derived allele count (DAC). **(A)** Proportion of variants per DAC bin across the cohort. **(B)** Geometric mean allele age (generations ago) by DAC, stratified by PCA groups (AFR, AMR, EAS, SAS, NFE, ASJ, MID) plus remaining individuals (unassigned to any one group) and a *nonspecific* category aggregating variants observed across multiple groups. Geometric means are restricted to groups with > 10 alleles in a given DAC bin.

**Figure S22:**
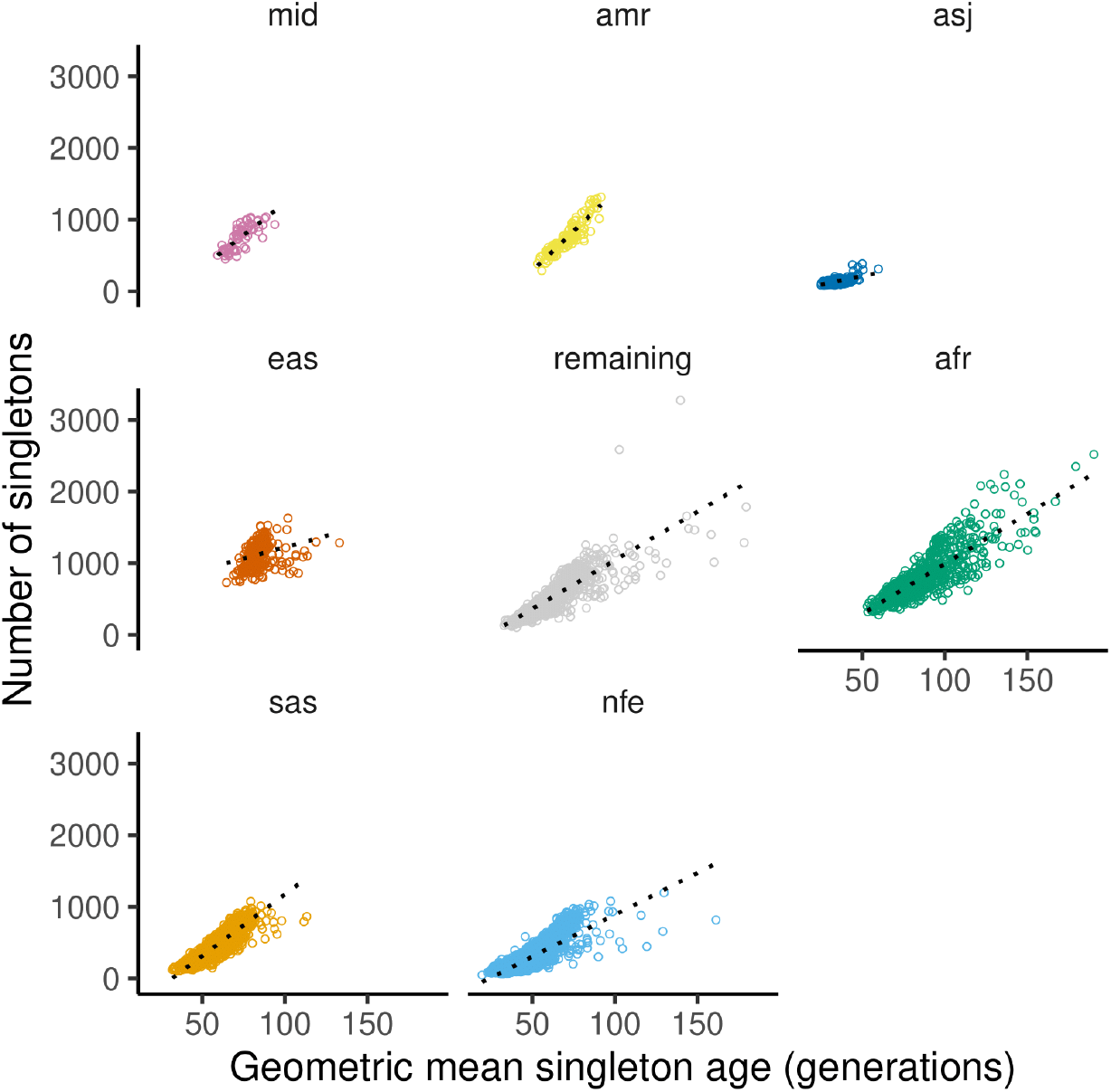
Number of singletons (GEL dataset-wide DAC = 1) per individuals vs the geometric mean age of those singletons, stratified by PCA groups (AFR, AMR, EAS, SAS, NFE, ASJ, MID) plus remaining individuals (unassigned to any one group). Analyses were restricted to a subset of 42,686 individuals without a < 3 degree relative in the GEL dataset.

**Figure S23:**
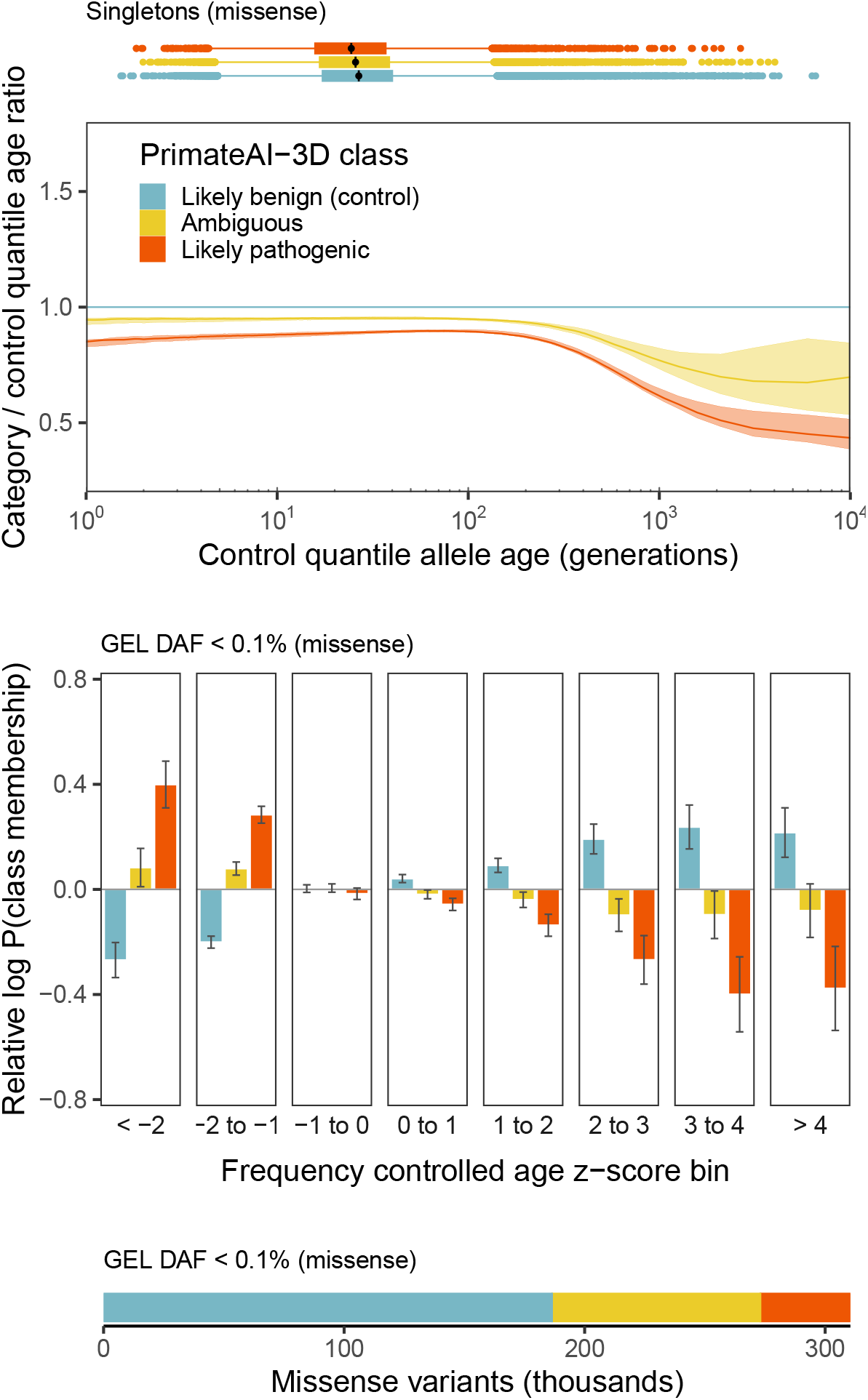
Allele age patterns associated with PrimateAI-3D classifications for ultra-rare missense variants. (i) Quantile–quantile analysis of singleton missense variants (DAC=1) comparing *ambiguous* and *likely pathogenic* classes against *likely benign* (control), with bootstrapped 95% confidence bands; younger ages (left) indicate stronger purifying selection. (ii) For ultra-rare missense variants (GEL DAF < 0.1%), the relative probability (odds ratios) of a variant being classified as *likely pathogenic* or *ambiguous* across frequency-controlled allele-age *z*-score bins (youngest < −2 to oldest > 4), estimated via logistic regression with 95% confidence intervals. (iii) Counts of ultra-rare missense variants by PrimateAI-3D class.

**Figure S24:**
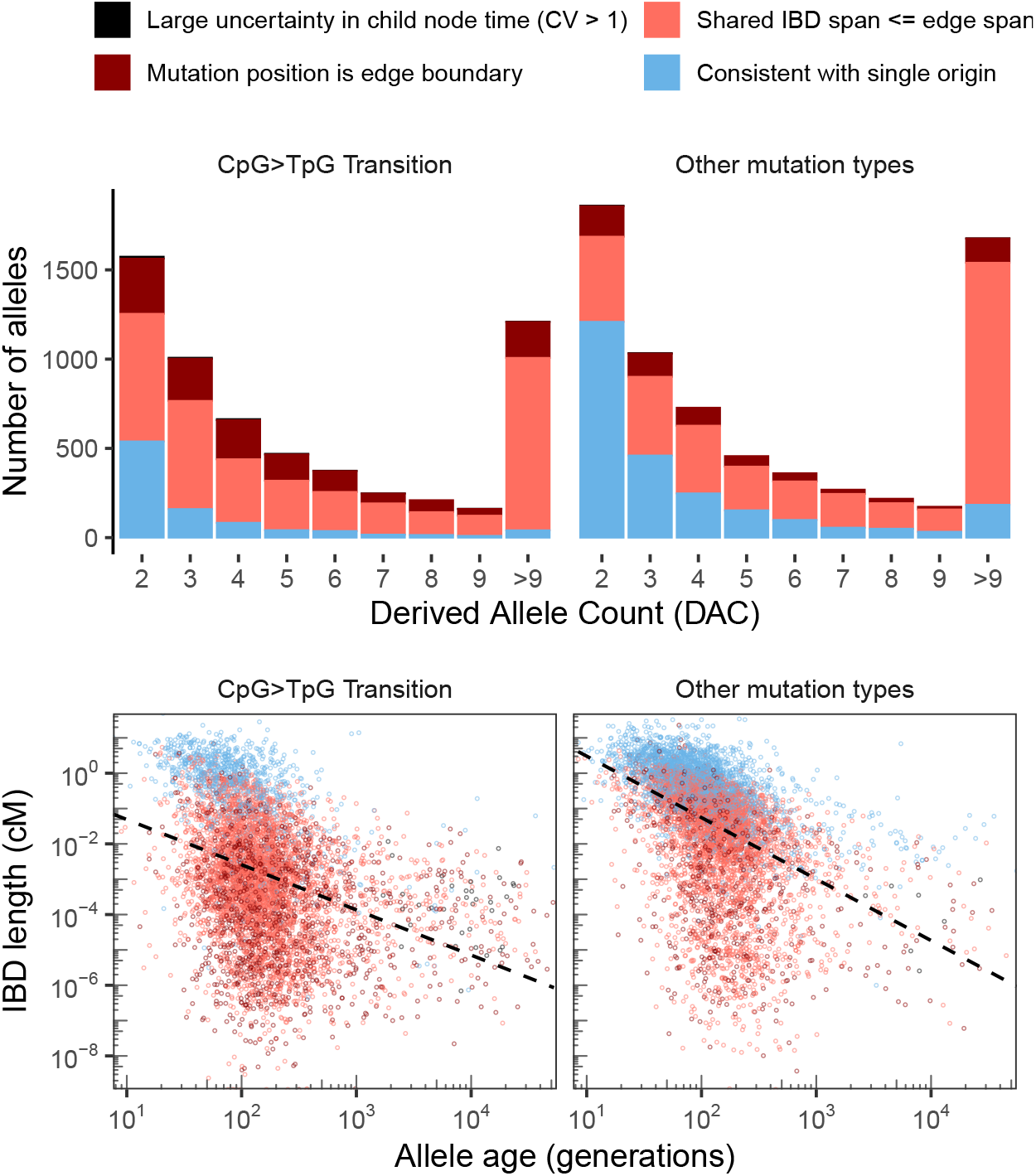
ARG-informed heuristics identify likely recurrent mutations and reveal expected relationships with allele age and IBD span. **Top:** Stacked bar plots showing the number of clinically classified variants (DAC > 1, DAF < 0.1%) stratified by derived-allele-count (DAC) bin and mutation type (left: CpG → TpG transitions; right: other mutation types). Colours indicate the outcome of the recurrence heuristic: large uncertainty in child-node time (CV > 1), mutation at edge boundary, shared IBD span ≤ edge span, or consistent with a single origin. CpG → TpG transitions show a higher fraction flagged as likely recurrent and the proportion of flagged variants increases with DAC, consistent with an elevated probability of independent mutation events at higher allele counts. **Bottom:** Relationship between derived allele age and IBD length (cM) on a log–log scale for CpG → TpG transitions (left) and other mutation types (right).

**Figure S25:**
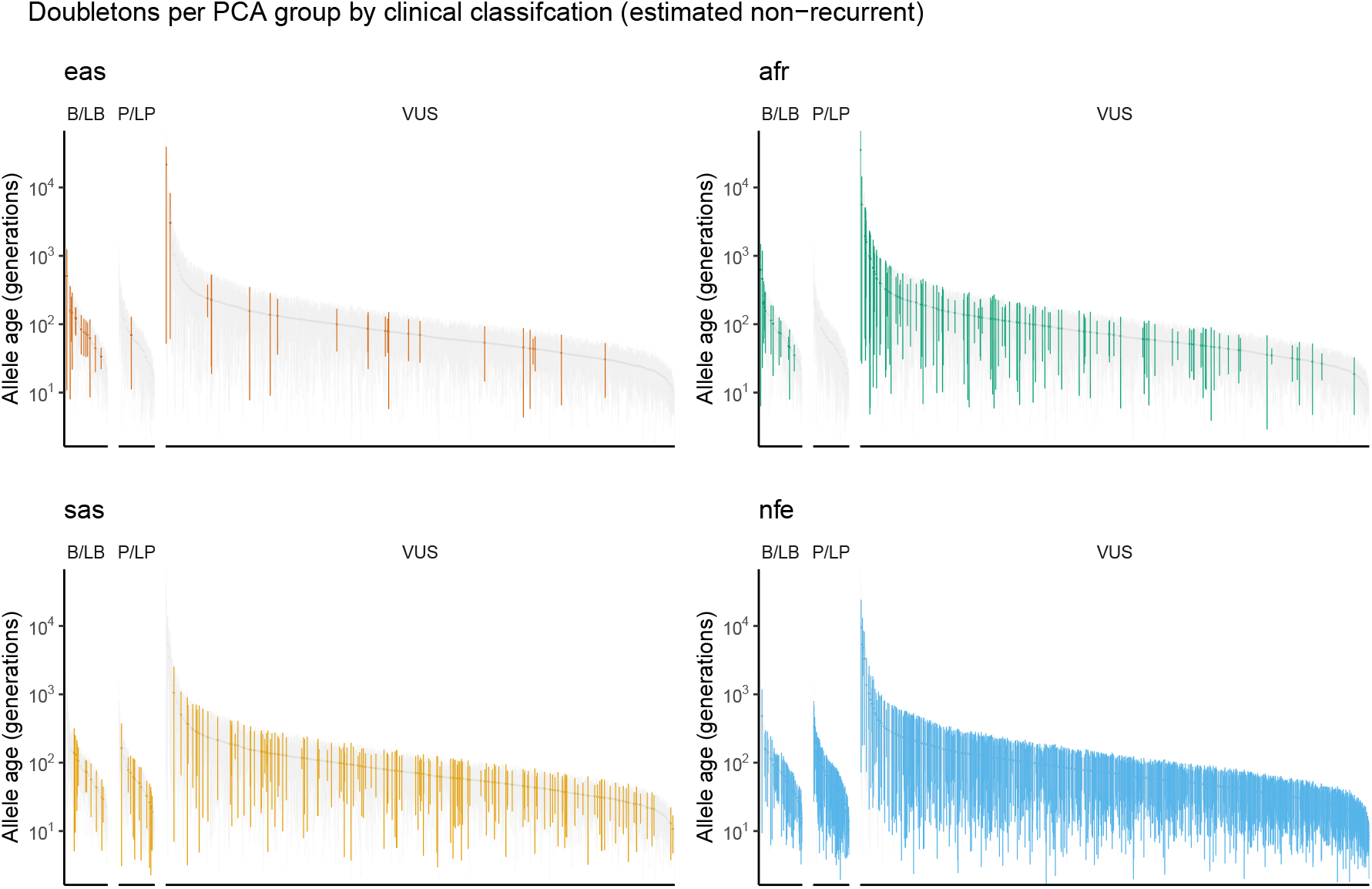
Allele ages for doubleton variants coloured by PCA group (AFR, EAS, SAS, NFE) and clinical classification in the GEL dataset. Points mark the estimated mutation age (generations ago); vertical intervals show the bracket between the ages of the ancestral and descendant nodes immediately above and below the mutation in the ARG. For comparison, the same variants are shown in all panels, with only those found in the corresponding PCA group colored. For each PCA group, panels correspond to the classes benign or likely benign (B/LB), pathogenic or likely pathogenic (P/LP), and variants of uncertain significance (VUS), as defined by the internal 100,000 Genomes Project rare disease clinical scientist questionnaires or ClinVar (22). Only doubletons inferred as non-recurrent (23) are shown.

**Figure S26:**
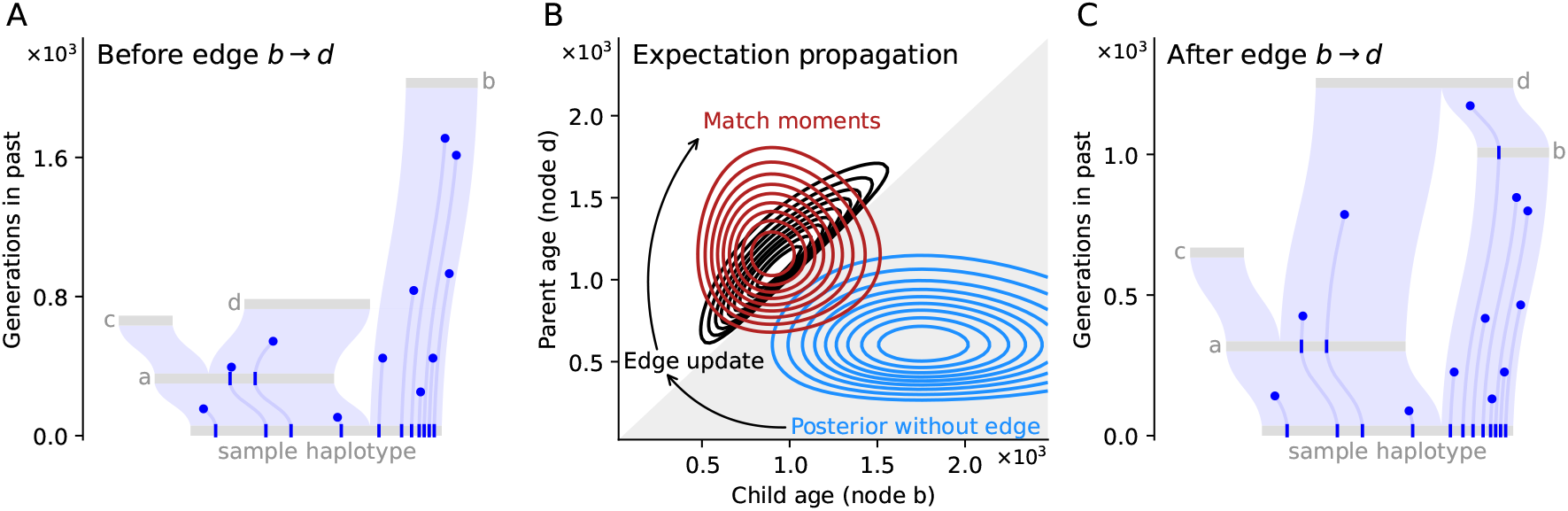
An example of an expectation propagation (EP) update, shown for edge *b* → *d* in Fig. 1. From left to right: (A) The dated ARG after iterating over all edges *except b* → *d*, before the update that incorporates information from this edge into the variational posteriors of nodes *b* and *d*. (B) EP maintains an approximating factor per edge that contributes to the variational posteriors of the attached parent and child nodes. Each factor is updated by first removing it from the variational approximation; multiplying the downdated approximation by the exact (Poisson) mutational likelihood for the edge (the “surrogate”); and then reparametrizing the held-out factor such that the updated approximation has the same moments as the surrogate. Incorporating edge *b* → *d* dramatically alters the posteriors of parent (*d*) and child (*b*) as the former must be older than the latter. (C) Iterating this moment matching condition propagates information across nodes, and results in variational posteriors that reflect both mutational density on edges and the topological constraints intrinsic to the ARG.

**Figure S27:**
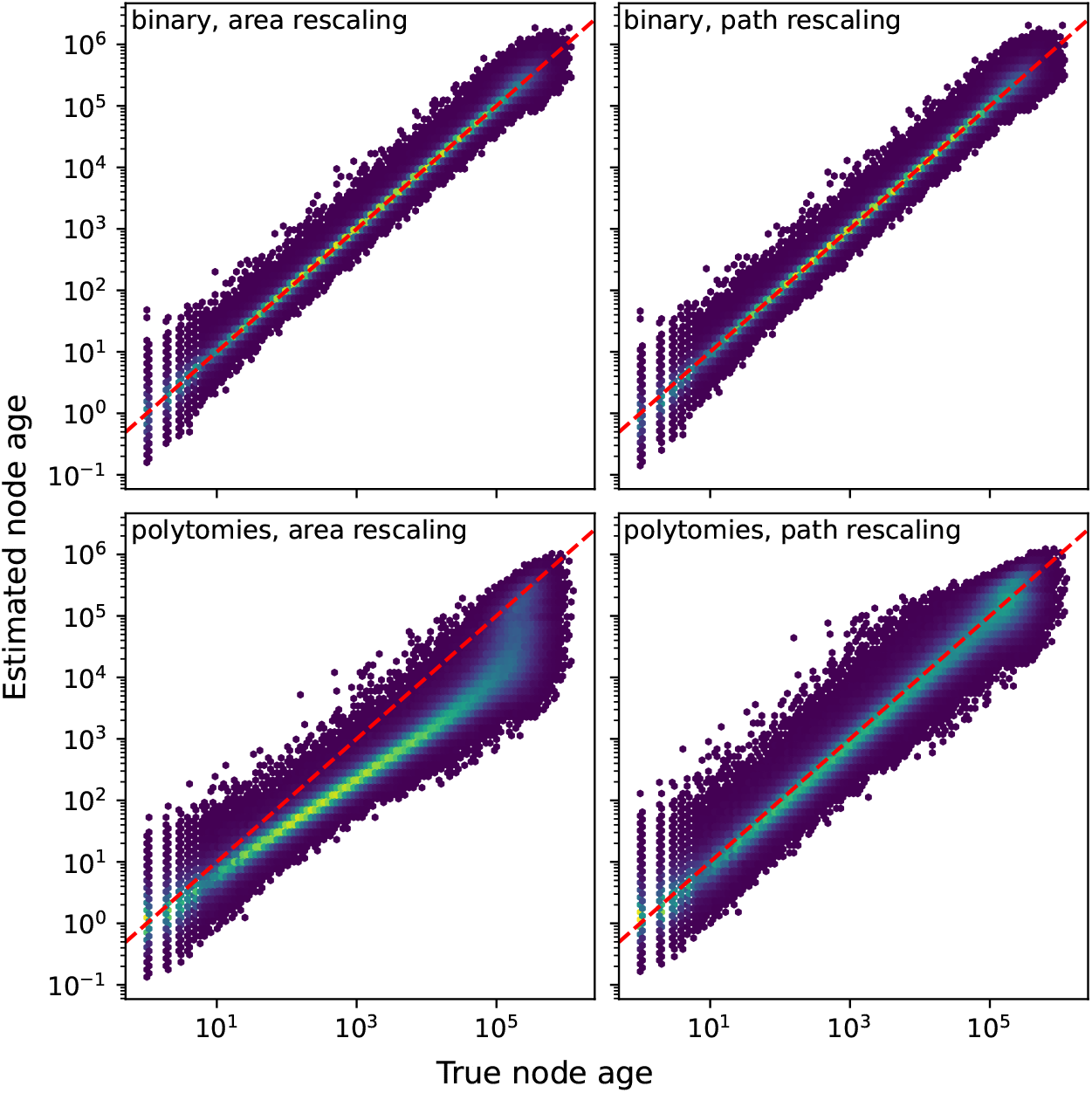
ARGs with artefactual polytomies are biased when rescaled by mutational area. Top row: When an ARG containing only binary trees are rescaled by either mutational area (left) or root-to-leaf path length (right), both approaches result in similarily well-calibrated time scales. Bottom row: The same ARG but with edges unsupported by mutations collapsed into polytomies, which introduces spurious mutational area. In this case, rescaling by mutational area will introduce bias and noise (left), but rescaling by root-to-leaf path length (right) will mitigate the bias. The ARG was generated via a discrete-time Wright-Fisher simulation of a randomly mating population of size 40,000 and 20,000 diploid samples; with 100Mb of sequence given a recombination rate of 10^−8^ and a mutation rate of 1.29 × 10^−8^.

**Figure S28:**
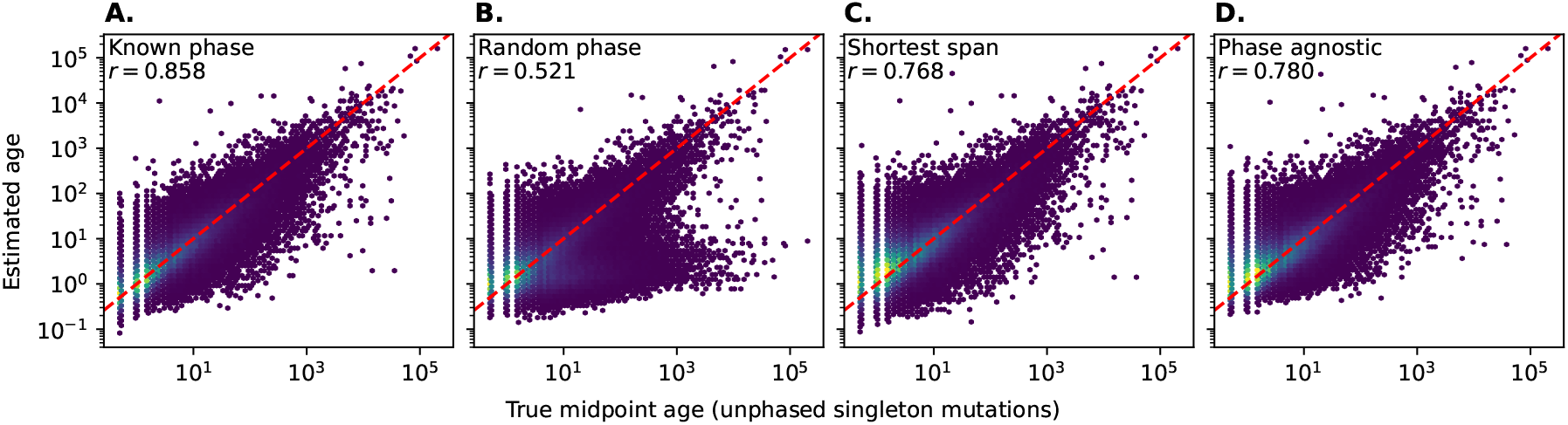
Phase-agnostic dating of singleton mutations. From left to right: (A) singletons dated with known phase result in unbiased age estimates (of the midpoint of the branch to which they are mapped). (B) If the phase is randomly flipped, some proportion of singletons will be mapped to longer or shorter branches and have erroneous ages. As a result, many older singletons will appear to be young. (C) Choosing singleton phase based on shorter edge span mitigates this error to a large extent. (D) The phase-agnostic dating algorithm described in SI §5 adapts this heuristic into a principled model, and propagates the uncertainty from the choice of phase into the posteriors for nodes and mutations. The ARG was generated via a discrete-time Wright-Fisher simulation of a randomly mating population of size 40,000 and 20,000 diploid samples; with 100Mb of sequence given a recombination rate of 10^−8^ and a mutation rate of 1.29 × 10^−8^, and subsequently inferred via tsinfer.

